# Woody species do not differ in dormancy progression: differences in time to budbreak due to forcing and cold hardiness

**DOI:** 10.1101/2021.04.26.441354

**Authors:** Al Kovaleski

## Abstract

Budbreak is one of the most observed and studied phenological phases in perennial plants. Two dimensions of exposure to temperature are generally used to model budbreak: accumulation of time spent at low temperatures (chilling); and accumulation of heat units (forcing). These two effects have a well-established negative correlation: the more chilling, the less forcing required for budbreak. Furthermore, temperate plant species are assumed to vary in amount of chilling required to complete endodormancy and begin the transition to breaking bud. Still, prediction of budbreak remains a challenge. The present work demonstrates across a wide range of species how bud cold hardiness must be accounted for to study dormancy and accurately predict time to budbreak. Cold hardiness defines the path length to budbreak, meaning the difference between the cold hardiness buds attain during the winter, and the cold hardiness at which deacclimated buds are predicted to open. This distance varies among species and throughout winter within a species. Increases in rate of cold hardiness loss (deacclimation) measured throughout winter show that chilling controls deacclimation potential – the proportion of the maximum rate response attained at high chill accumulation – which has a sigmoid relationship to chilling accumulation. For forcing, rates of deacclimation increase non-linearly in response to temperature. Comparisons of deacclimation potential show a dormancy progresses similarly for all species. This observation suggests that comparisons of physiologic and genetic control of dormancy requires an understanding of cold hardiness dynamics and the necessity for an update of the framework for studying dormancy and its effects on spring phenology.

## Introduction

Spring phenology defines the long-term survival and fitness of perennial plants within environments with below-freezing winter temperatures (1,2). Spring kills of breaking buds are widely regarded as a major aspect influencing species distribution (3), and are expected to increase in frequency due to climatic changes (4), making accurate predictions key to understanding adaptation (5). Yet accurate predictions still remain a major challenge (6,7), and climate change may create environmental conditions in colder climates that are not analogous to those in presently warmer regions (8). Artificial warming experiments have been used in order to study possible future conditions (9,10), but results from these experiments do not match observations of advanced budbreak over the last few decades of climate warming (6). This mismatch demonstrates that we still lack a predictive understanding of the budbreak process, and thus empirical studies are required to understand the relationships that define the time to budbreak in woody species (7).

Considerable prior research has attempted to predict the effects of climate change on phenology (3,6,11–20). To date, this work has been limited to assessing the interactions between chilling (accumulation of time spent in low temperatures) and forcing (accumulation of thermal time, growing degree-days) as temperature effects on budbreak, with reference to shifts in their interaction in a changing climate (6,11,17,21). However, a missing key component of winter survival, the dynamic and changing cold hardiness of buds, has remained unexplored as the starting point for estimation of the time to achieve any phenological stage. Instead, most phenological models fail to account for species specific differences in cold hardiness for buds of all temperate species, at all times during the dormant season, and in any location or climate, despite existing empirical evidence that artificial acclimation increases time to budbreak (22). It is also widely known that species have different temperature thresholds for tissue damage during bud swell, budbreak, and leafout (5,18), as well as in midwinter (18,22,23). This indicates that the amount of cold hardiness buds need to lose to transition from their cold hardy state to budbreak (the deacclimation path length) differs among species and climates, and throughout the dormant season. The influence of cold hardiness has remained largely unexplored on these processes, despite information existing on the effects on budbreak (22,24).

Another critical source of uncertainty in predictions of phenology is our poor understanding of dormancy. Time spent in low temperatures (chilling) is known to promote the transition from a dormancy phase that is non-responsive to growth-conducive temperatures (i.e., endodormancy, rest) to a responsive phase (ecodormancy, quiescence) (25). Some species, or genotypes within species, are understood to have lower chilling requirements than others based on faster budbreak. This results in comparative classifications of “low chill” and “high chill requirement” plants. However, there are still questions about which temperatures promote chilling and how much chilling they provide (26,27), leading to multiple methods for estimation of chill accumulation based on temperature (28–32). This is in part due to the fact that a mechanism for dormancy has yet to be resolved, despite a growing list of dormancy related genes and chemicals that promote dormancy transitions (33–38). Recent work has shown that rates of cold hardiness loss (deacclimation) increase with chilling accumulation, which can be used to study dormancy progression (24). By standardizing the rates to the maximum observed at the end of the season, this measurement is referred to as deacclimation potential – *Y_deacc_* – which varies from 0 to 1 [or 0% to 100% (24)]. The *Y_deacc_* is the increase in responsiveness observed as chill accumulates, and is a quantitative measurement of dormancy. In analogous terms, *Y_deacc_* = 0 would mean entirely endodormant (non-temperature responsive buds), and *Y_deacc_* = 1 would mean entirely ecodormant buds (maximum temperature responsiveness), acknowledging a quantitative progress in dormancy rather than a qualitative transition.

Three sources of variation in time to budbreak are thus investigated in the present study: the initial cold hardiness of buds (the departure point, *CH_0_*), the cold hardiness at budbreak (*CH_BB_*), and the effective rate of cold hardiness loss (deacclimation; 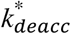). The relationship of these variables is used here as:

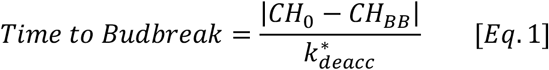

where |*CH_0_* – *CH_BB_*| defines the path length from the cold hardy, fully dormant state to budbreak. An example of this relationship is shown in **Fig. 1**: two rates of deacclimation, 4 and 1 °C day^−1^, are given for a common path length of 20 °C. These two deacclimation rates result in 5 and 20 days to budbreak, respectively. This relationship (**Eq. 1**) demonstrates the uneven effect of path length on time to budbreak: if the path length increased by 5 °C for both, to a total of 25 °C (e.g., buds from a region with lower minimum temperatures having a lower *CH_0_*), time to budbreak would increase by 1 and 5 days at the same rates of deacclimation (hypothetical scenarios are presented more extensively in **SI Appendix Notes, SI Appendix Figs. S1-S5**). Moreover, it is expected that different species will vary in each of these three aspects: different levels of cold hardiness throughout the winter, different thresholds to attain budbreak, and different rates of deacclimation. Because timing of budbreak is affected by chill accumulation and spring temperatures, the hypothesis is that rates of deacclimation will also be affected by both chill accumulation and ambient spring temperature within the same species.

**Figure 1.**
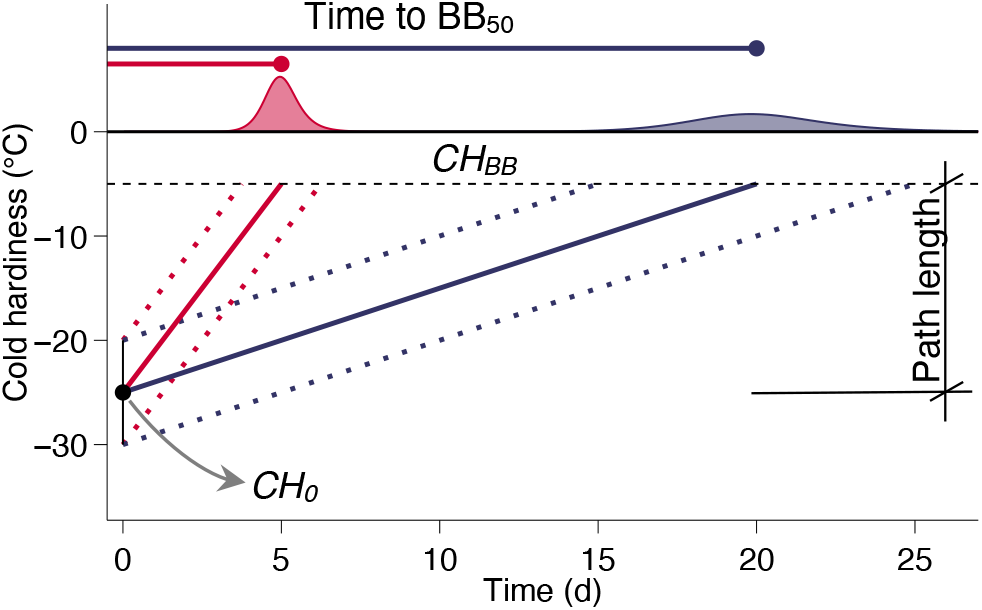
Sources of variation in time to budbreak. In order to reach budbreak, buds must lose their initial cold hardiness (*CH_0_*) and reach the cold hardiness at budbreak (*CH_BB_*). This distance is the path length (in °C). For the same path length, time of budbreak is then defined by how fast the path is traveled: the effective rate of deacclimation 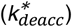 A higher rate of deacclimation (red) leads to concentrated and earlier budbreak, whereas a lower rate (blue) means budbreak occurs later and more sporadic (density curves).

To study these effects, over 40,500 cold hardiness measurements and 8,000 cuttings for budbreak observation from 15 different species collected over two dormant seasons were used as proxy for whole-plant responses (39). The 15 species represent both Gymnosperms and Angiosperms, deciduous and evergreen, from eight different families (Cornaceae, Ericaceae, Fabaceae, Fagaceae, Oleaceae, Pinaceae, Rosaceae, and Sapindaceae), from three different continents and many climates of origin, of horticultural and ecological importance, with both regular and naked buds. The phenotype that unites all species studied is supercooling of water as a method for bud survival (40). In general, four species are presented in the main figures as examples for the concepts (*Abies balsamea*, *Cercis canadensis*, *Forsythia* ‘Meadowlark’, and *Larix kaempferi*), but results for all species are presented in Supporting Information.

## Results

The species studied differed in their cold hardiness determined weekly throughout the winter (**Fig. 2, SI Fig. S6**). However, the same pattern is observed: cold hardiness is gained in the fall, maintained during the winter, and lost before budbreak in the spring (density curves show field budbreak distribution in **Fig. 2, SI Fig. S6**). This response generally follows air temperature (**SI Appendix, Fig. S7**). For every date of cold hardiness determination, buds were also placed in a growth chamber at 22 °C to observe time to budbreak. Similar patterns were also observed across species in terms of time to budbreak: buds take a long time to break in the fall, whereas time to budbreak decreases exponentially starting in late November (**Fig. 2**, **SI Appendix, Fig. S8**).

**Figure 2.**
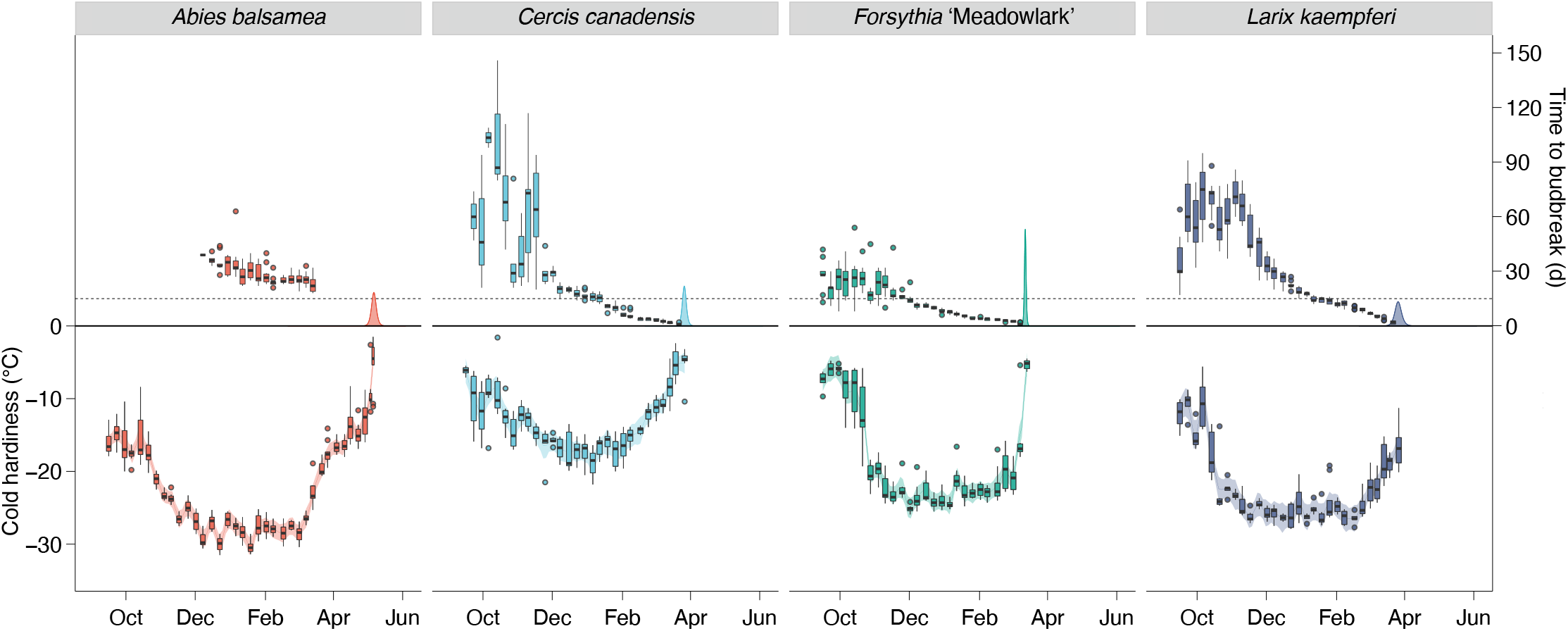
Cold hardiness and time to budbreak. Cold hardiness of buds measured weekly throughout the 2019-2020 dormant seas (bottom). At each collection, time to reach budbreak under forcing was also measured (top). Density curves above 0 line represent observed budbreak in the field (see **Extended Data Figure 8**). Dashed line shows 15 days threshold used for chilling requirement determination.

### Effect of chilling accumulation (dormancy) on rates of deacclimation

Besides weekly field cold hardiness and time to budbreak, loss of bud cold hardiness was tracked between weekly field collection and budbreak for each collection at a single temperature. These buds were in the same growth chamber as those where budbreak was evaluated (22 °C). From these data, effective rates of deacclimation 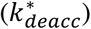 were determined for each collection in °C of cold hardiness lost per day (e.g., **Fig. 3A**; adjusted r^2^ = 0.79, *F_1588,30425_* = 76.8, *P*<0.001, for all species all collections combined). The 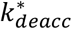 (slope of the linear regressions in **Fig. 3A)** increased with the accumulation of chill during winter for all species (**Fig. 4A, SI Appendix Fig. S9**). Based on deacclimation and budbreak at different chill accumulations, it is notable that budbreak (presented along the 0 °C line in **Fig. 3A**) is linked to the dynamics of deacclimation: lower 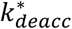at 23 chill portions leads to erratic, delayed budbreak, as expected based on **Fig. 1**. In turn, budbreak occurs much more synchronously with higher 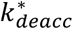 at 84 chill portions (i.e., height of density curves in **Fig. 3A**). This effect is also observed in species comparisons: at 84 chill portions, buds break earlier and with more uniform timing in *Cercis canadensis* than in *Abies balsamea*.

**Figure 3.**
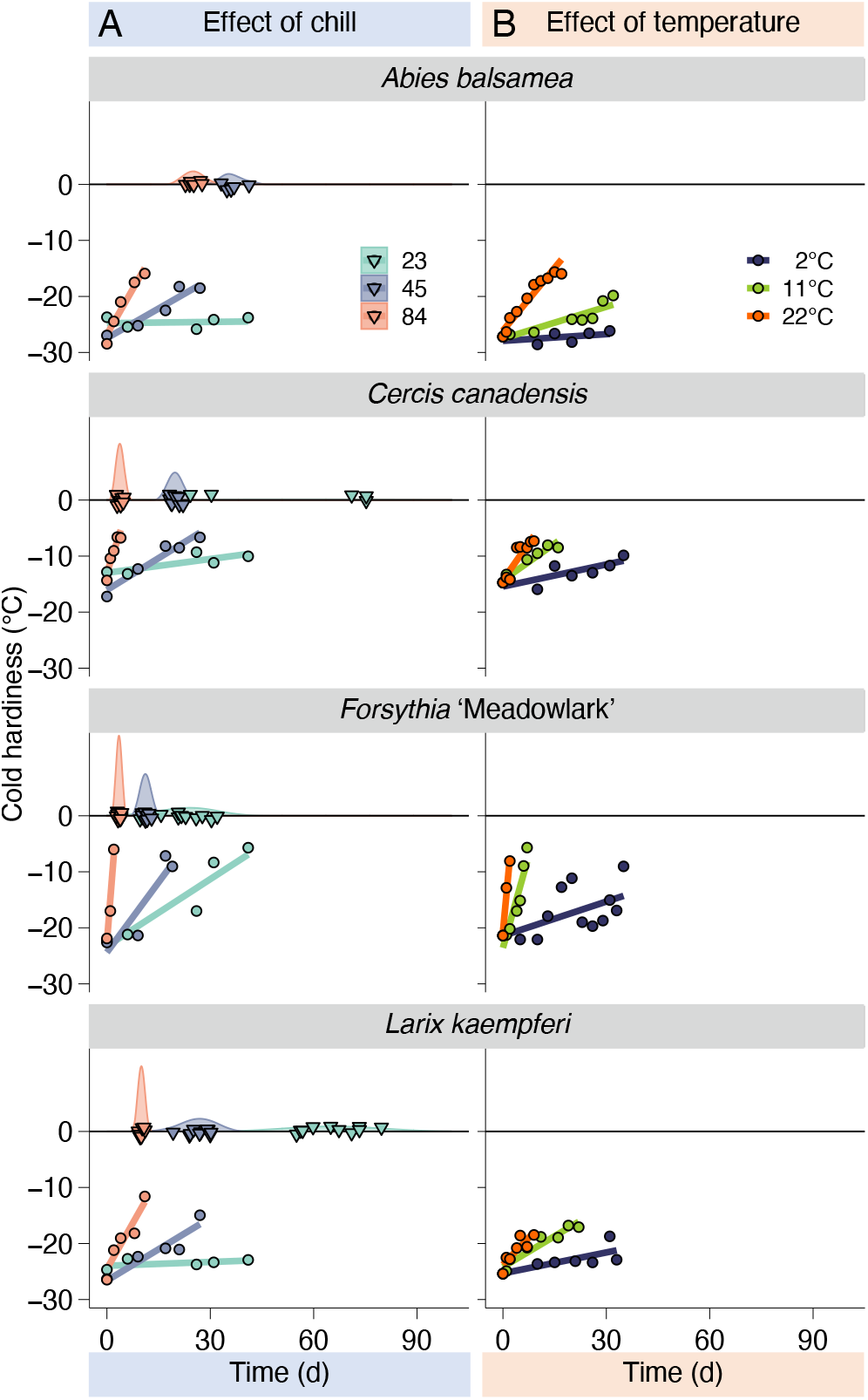
Different deacclimation rates and budbreak at different chill accumulations and temperatures. Loss of cold hardiness measured at different levels of chill accumulation, but same temperature (*A*) and at different temperatures but same chill accumulation (*B*). At different levels of chill accumulation, budbreak was also measured (upside down triangles and density curves at 0 °C line, *A*).

**Figure 4.**
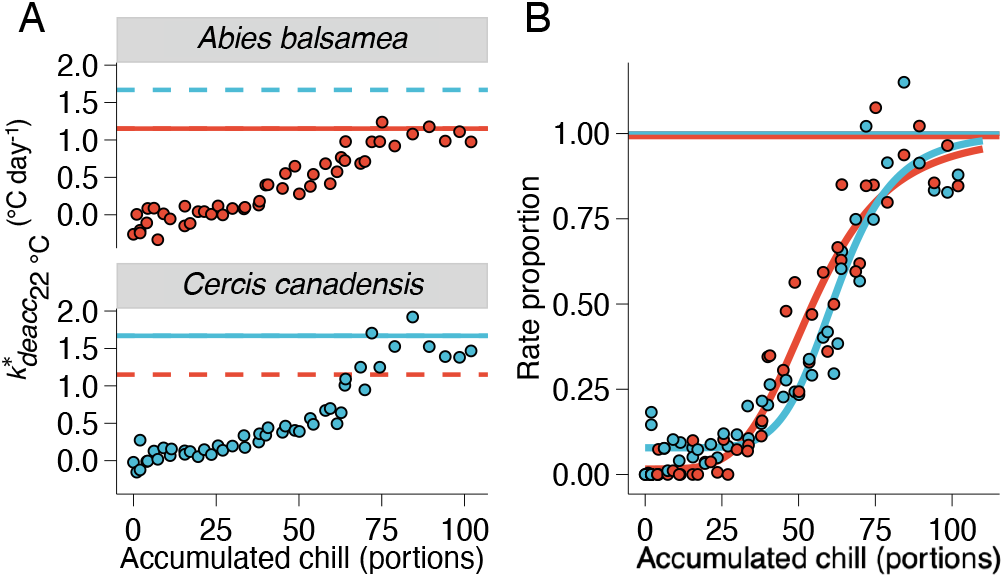
Increase in deacclimation rates in response to chill accumulation. Rates of deacclimation increased with chill accumulation (*A*). Full line shows maximum rate of deacclimation at 22 °C as the average of the four highest deacclimation rates for each species. Dashed line shows the maximum rate of the other species for comparison. When normalized to the maximum rate of deacclimation for each species, a normalized rate proportion is produced (*B*).

The maximum rate of deacclimation observed at 22 °C 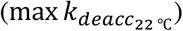 differed among species (**SI Appendix Fig. S10**). The 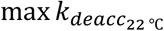 ranged from 0.6 °C day^−1^ (*Rhododendron calendulaceum*) to 9.5 °C day^−1^ (*Forsythia* × ‘Meadowlark’). Eleven out of the 15 species studied exhibited deacclimation rates between 0.5 and 2.0 °C day^−1^. Effective rates of deacclimation 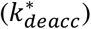 from each species were then normalized to their corresponding 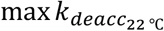, generating rate proportion measurements (*Y_deacc_*) and isolating the effect of chilling accumulation (**Figs. 4A and 4B, SI Appendix Fig. S11;** see **SI Appendix Notes, Section 4, for further explanation**). Here, *Y_deacc_* was modeled as a three-parameter log-logistic:

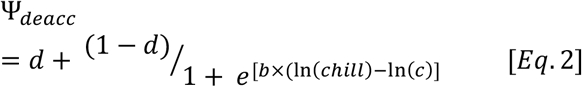

where the potential for deacclimation is a function of chilling accumulation (*chill*, here in portions). Three estimates are thus obtained: *d* is the minimum deacclimation observed at all times (analogous to an intercept); *b* is the slope associated with the log-logistic curve, where negative values indicate increases in *Y_deacc_* as chill accumulation increases, and greater magnitudes indicate a higher slope; and *c* is the inflection point of the log-logistic curve.

Based on deacclimation potential, dormancy progressed extremely similarly across this wide range of species of different field phenology (**Fig. 5A, SI Appendix Fig. S12**). The inflection point *c* of these sigmoid curves varied between 55 and 70 chill portions (**Fig. 5B**; between 52 and 76 chill portions for all 15 species). In stark contrast, the currently standard method of determining chilling requirement based on an arbitrary choice of time for 50% budbreak [here chill required for 50% budbreak within 15 days was used (dashed line in **Fig. 2**)] produced values ranging from 37 to 96 chill portions. Based on these observations, *Forsythia* ‘Meadowlark’ would be considered to have a low chill requirement, and become ecodormant in early winter in the location of this study; whereas *Abies balsamea* would be a high chill requirement species, and become ecodormant only in the beginning of spring. The chill requirement based on date of budbreak is highly negatively correlated with the 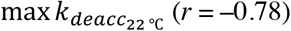. This means that as chill accumulates over the progression of winter, species with faster maximum rates will reach the rate necessary to lose cold hardiness and break bud within 15 days at lower chill accumulation than those with slower maximum rates. However, deacclimation rates continue to increase in response to chilling. The estimated effective rate of deacclimation at 22 °C at any point in chilling accumulation is thus:

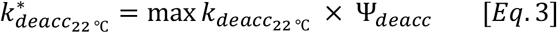

**Figure 5.**
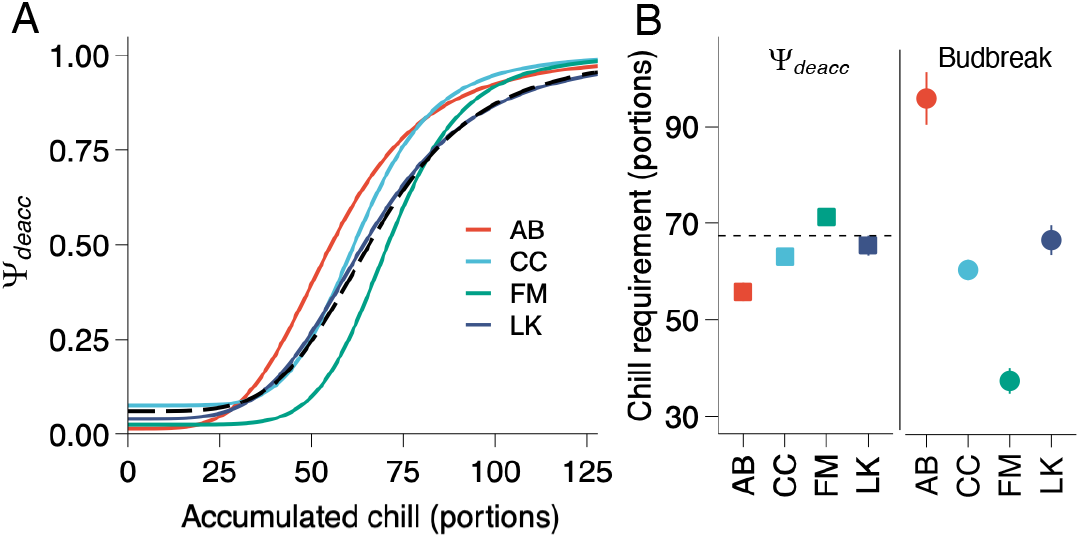
Dormancy progression based on deacclimation rates and time to budbreak. Normalized rates of deacclimation produce the deacclimation potential for all species (*A*; 51≥*n* ≥49). Inflection points of the deacclimation potential (*c* term from **Eq. 2**; curves in *A*) are very similar across species, unlike chill requirement based on budbreak (*B*). Dashed lines (*A* and *B*) show the average based on deacclimation potential of all 15 species studied. Error bars, when visible, represent standard deviation. AB – *Abies balsamea*; CC – *Cercis canadensis*; FM – *Forsythia* ‘Meadowlark’; LK – *Larix kaempferi.*

However, different temperatures can result in different maximum rates of deacclimation.

### Effect of temperature on rates of deacclimation (forcing)

To evaluate how temperatures affect the rates of deacclimation, buds were collected from 13 of the 15 species at 54 and 82 portions of chill accumulation (not all 13 were collected at both times). These buds were deacclimated at 7 different temperatures between 2 and 30 °C. As expected, lower temperatures result in slower deacclimation as compared to higher temperatures (**Fig. 3B**). The slopes for each temperature were extracted as deacclimation rates (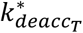, the effective rate of deacclimation at temperature *T*) using a linear model (*adjusted r^2^* = 0.66, *F_287,7384_* = 53.9, *P*<0.001, for all temperatures and genotypes combined). Because these collections did not occur at chilling accumulations where *Y_deacc_* = 1, the values were normalized based on the rates measured at 22 °C at each collection and 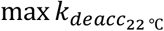 in order to isolate the effect of temperature. This correction transforms values of the measured, effective rates of deacclimation at a given temperature 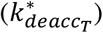 below maximum chill accumulation (*Y_deacc_* < 1) into the maximum possible rate at that temperature 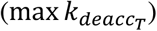, based on **Eq. 3**.

The corrected rates were then used to model the response of rates of deacclimation to temperature (**Fig. 6; SI Appendix Fig. S13**). While a linear model produces a good fit for 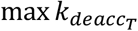 as a function of temperature (*adjusted r^2^* = 0.95, *F_25,89_* = 87.8, *P*<0.001, for the 13 species combined), it is clear that the response to temperature increases faster at lower temperatures and tapers off at warmer temperatures for some species (e.g., Forsythia × ‘Meadowlark’ in **Fig. 6**). Therefore, a polynomial of the third order with intercept = 0 to allow for these curvatures was used. The effective rate of deacclimation at any temperature and chill accumulation thus becomes:

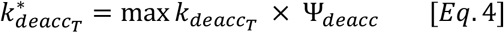

where the effective rate of deacclimation at temperature *T* is a function of both temperature (through 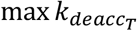) and chill accumulated (through *Y_deacc_*).

**Figure 6.**
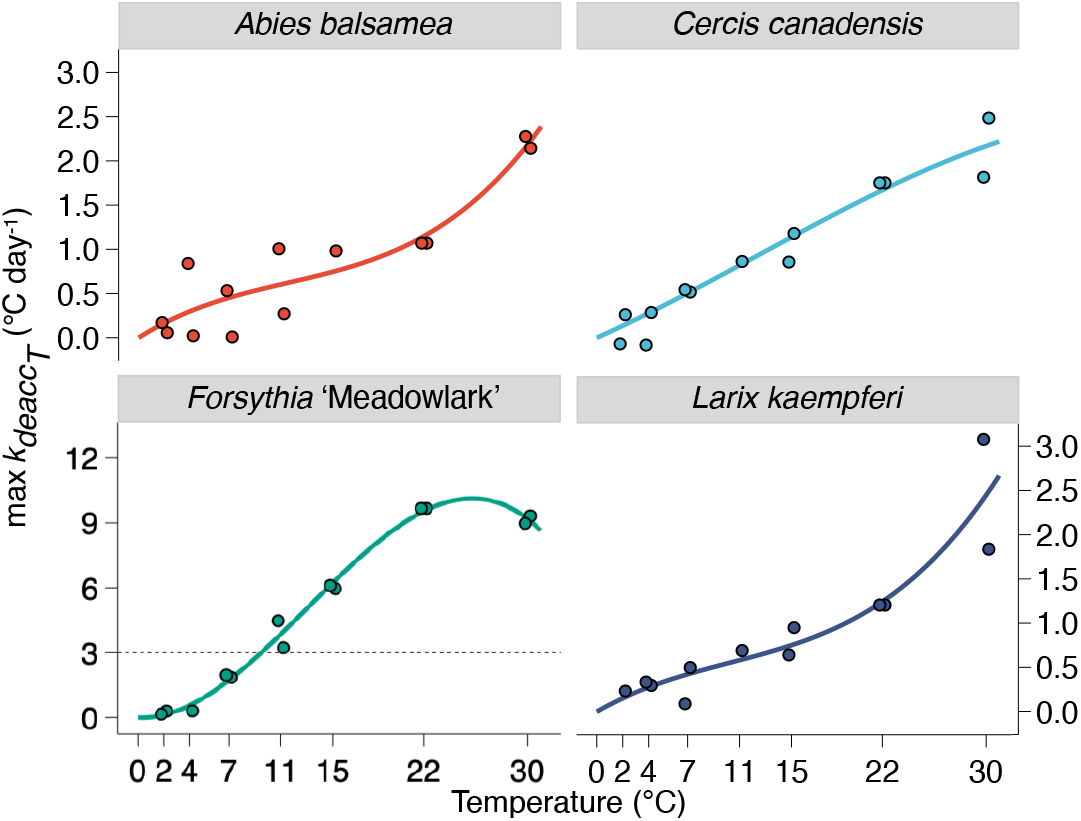
Deacclimation rates in response to temperature. Deacclimation rates increase with increasing temperatures for all species (*n* = 14). A different scale was used for *Forsythia* ‘Meadowlark’, but dashed line shows maximum value of scale for other species.

Yet despite accounting for the effects of forcing temperature and prior chilling on the rate of cold hardiness loss, differences in time to budbreak can still arise from how much cold hardiness a bud must lose for budbreak to be perceived (see **Fig. 1**, **Eq. 1**).

### Effect of cold hardiness (path length)

To obtain the path length from a cold hardy state to budbreak for each species in field conditions, two measurements are required: the maximum cold hardiness (*CH_max_*, used as *CH_0_*) and the cold hardiness at budbreak (*CH_BB_*) (**Fig. 1**). *CH_max_* was obtained by averaging the cold hardiness at the four timepoints during the winter where cold hardiness in the field was maximum for each species (**Fig. 7A, SI Appendix Fig. S14**).

**Figure 7.**
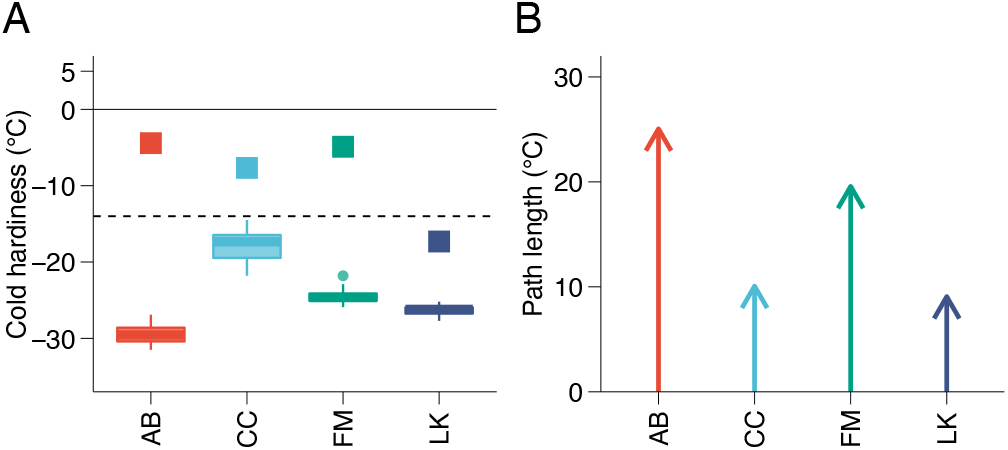
Cold hardiness parameters for each species. Maximum cold hardiness (*CH_max_*) observed in the 2019-2020 season (boxplots) and cold hardiness at budbreak (*CH_BB_*; squares) for each species (dashed line shows minimum temperature observed in the season for an idea of safety margins; *A*). Path length (|*CH_max_* - *CH_BB_*|) to budbreak for each species (*B*). AB – *Abies balsamea*; CC – *Cercis canadensis*; FM – *Forsythia* ‘Meadowlark’; LK – *Larix kaempferi.*

To estimate *CH_BB_*, the deacclimation curve at each collection was projected from all initial cold hardiness measurements to the time at which each bud broke and averaged (see **SI Appendix Notes, Fig. S1**). *CH_BB_* values for the four main species ranged from −17 °C to −4 °C (**Fig. 7A**). With values of *CH_BB_* and *CH_max_* determined, the path length is thus obtained as |*CH_max_* − *CH_BB_*|, and values ranged from 9 °C to 25 °C (**Fig. 7B**). Note that although *Larix kaempferi* buds are about 10 °C more cold hardy than *Cercis canadensis*, the two species have very similar path lengths to budbreak because of different cold hardiness at budbreak.

### Combining effects of chilling, temperature, and path length can predict field budbreak

The balance between acclimation and deacclimation results in the cold hardiness at any given moment during the dormant season (**Fig. 2**), and thus the path length to budbreak at any moment during the season. In the fall, although warm temperatures are still present 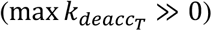, chill accumulation is very low (Ψ_*deacc*_ ≅ 0), therefore very little deacclimation can occur (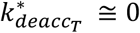, see **Eq. 4**). As chill accumulates during the winter, the potential for deacclimation increases (Ψ_*deacc*_ > 0). However, winter temperatures are generally too low for significant levels of cold hardiness loss to occur 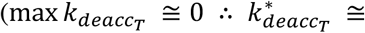 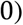. But once both deacclimation potential is high enough (Ψ_*deacc*_ > 0) and spring temperatures are in place 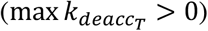, buds are able to deacclimate 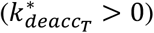. Based on this, the field cold hardiness can be described as:

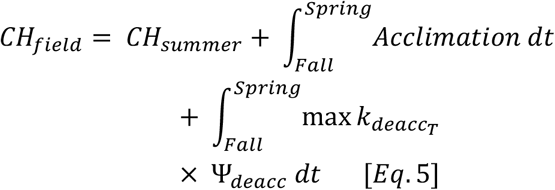

where *CH_summer_* is the cold hardiness of buds as they form in the late summer and fall, likely a function of species or genotype; *Acclimation* is a function of temperature where generally low temperatures promote greater gains in cold hardiness, but not part of this study; and 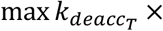 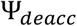 describes any deacclimation occurring. Both acclimation and deacclimation portions are integrated over time (*t*). Through these relationships, cold hardiness throughout the winter can be predicted: in the fall, acclimation predominates leading to gains in cold hardiness; in the spring, deacclimation predominates leading to loss of cold hardiness. Additionally, field budbreak is predicted to occur in the spring once *CH_field_* = *CH_BB_*, therefore connecting the dormant and growing seasons.

To test whether prediction of field budbreak was possible, the time to run the path from maximum cold hardiness in the field until the cold hardiness at budbreak was predicted based on the temperature and chill relations established in growth chamber experiments. For this, *CH_max_* and *CH_BB_* were used for each species, and their growth chamber-determined forcing 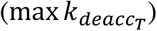 and chilling (*Y_deacc_*) responses. In addition, a set acclimation rate (*k_acc_*) was used for all species (limited to not increase observed *CH_max_* – see **Materials and Methods**). Therefore, cold hardiness in the field is described here as:

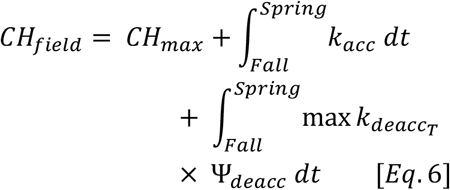

where the trajectory predicted was only that of the loss of cold hardiness in the spring, and the predicted day of budbreak occurred when *CH_field_* = *CH_BB_*. In this way, ***Eq. 6*** predicts the path of cold hardiness loss in late winter and spring and, based on this information, the date when budbreak happens can be inferred (**Fig. 8, SI Appendix Fig. S15**).

The resulting relationship between predicted (using ***Eq. 6***) and observed budbreak for all 15 species used in this study combined is 1.98 + 1.01 × *BB_observed_* (**Fig. 9**; *adjusted r^2^* = 0.91, *F_1,13_* = 148.1, *P*<0.001), where *BB_observed_* is the day of 50% budbreak in the field. It is important to note that: (i) the only field estimated parameter is *CH_max_*; (ii) there was no optimization of parameters obtained from growth chamber experiments (no training of the model); and (iii) the same acclimation rate was used for all species.

**Figure 8.**
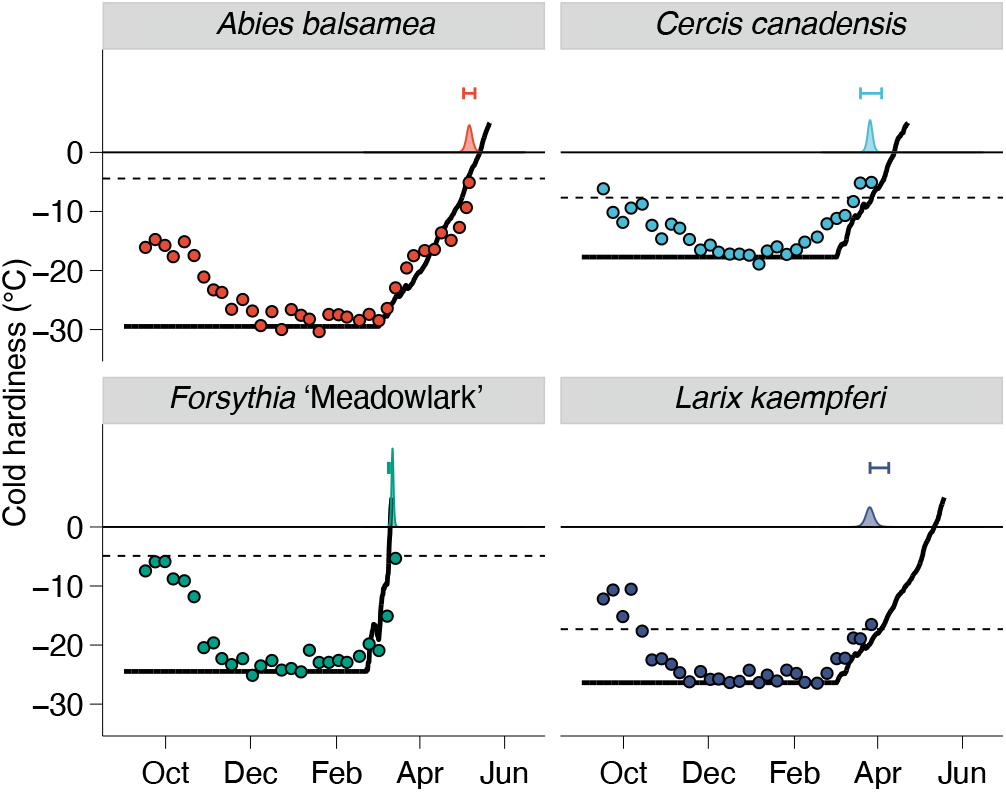
Predicting budbreak based on cold hardiness dynamics. Average cold hardiness [circles, no error for simplicity (see **Fig. 2**)] during the entire season are shown, as well as field budbreak (density plots at 0 °C). Predicted cold hardiness started at maximum cold hardiness (*CH_max_*) and runs until +5 °C based on species-specific deacclimation parameters (solid line). Budbreak is predicted to occur when prediction line crosses the cold hardiness at budbreak (*CH_BB_*; dashed lines) for each species. The interval is produced by adding error of ± 2.5 °C to *CH_max_*.

**Figure 9.**
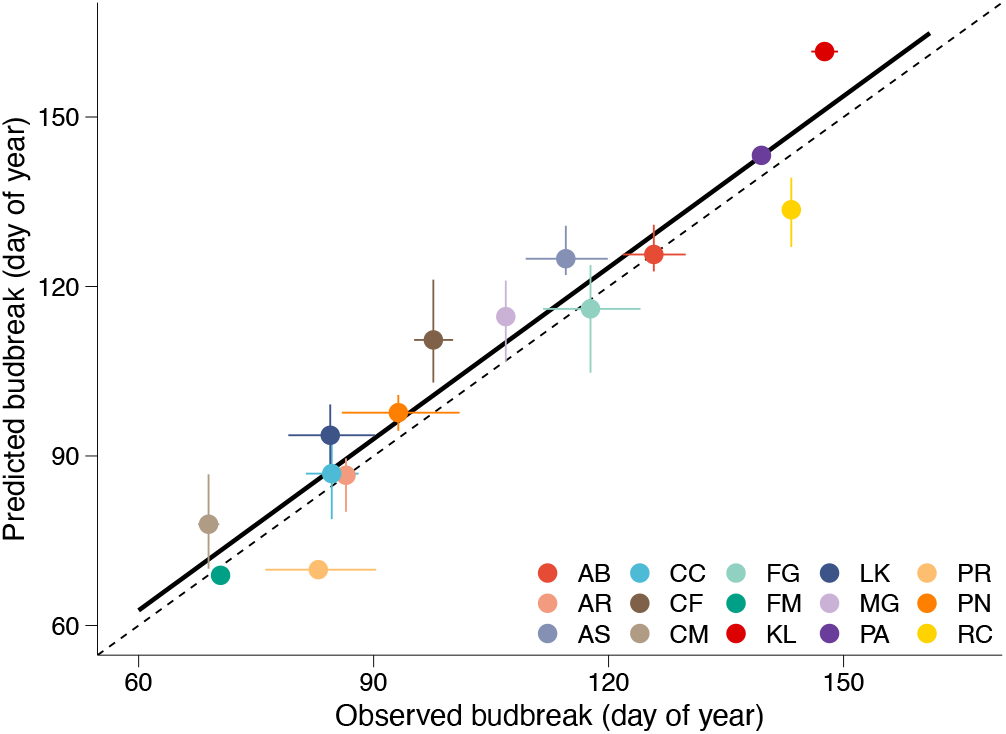
Evaluation of budbreak predictions based on cold hardiness dynamics. Relationship between observed and predicted budbreak based on deacclimation dynamics. Dashed line represents 1:1 relationship; full line shows calculated relationship: BB_predicted_ = 1.98 + 1.01 × BB_observed_, adjusted *r^2^* = 0.91, *P*<0.001. Horizontal error bars are distribution of field budbreak from 5% to 95%; vertical error bars are estimated error of prediction based on changes in initial cold hardiness (*CH_max_* ± 2.5 °C). AB – *Abies balsamea*; AR – *Acer rubrum*; AS – *Acer saccharum*; CC – *Cercis canadensis*; CF – *Cornus florida*; CM – *Cornus mas*; FG – *Fagus grandifolia*; FM – *Forsythia* ‘Meadowlark’; KL – *Kalmia latifolia*; LK – *Larix kaempferi*; MG – *Metasequoia glyptostroboides*; PA – *Picea abies*; PR – *Prunus armeniaca*; PN – *Prunus nigra*; RC – *Rhododendron calendulaceum.*

## Discussion

As the world becomes progressively warmer, spring phenology has continued to advance in time. However, the pace of advance in time to budbreak appears to be decreasing. Climate warming has led to shifts in chill accumulation (8), where decreases in chill have been speculated to counterbalance increasingly warmer springs (21,41,42). An effect that is largely ignored, however, is that warmer winter temperatures can also lead to less cold hardy buds. Here, changes in cold hardiness are demonstrated to affect the path to budbreak, and thus an important but neglected dimension of spring phenology. Most importantly, through the investigation of cold hardiness dynamics, dormancy progression across a wide range of diverse woody perennial species spanning the seed plant phylogeny is very similar. This suggests classifications of low or high chill requirement are based mostly on forcing response.

### Acknowledging cold hardiness in phenological models

This study uses aspects of cold hardiness dynamics to make inferences about budbreak phenology. This demonstrates the existence of a phenotype that is measured easily, and much more so than determination of internal development of buds [e.g., (43,44)], and that can be measured prior to any external development in budbreak progression. In addition, cold hardiness dynamics clearly demonstrate the negative relationship between chilling and forcing (as plasticity; 11,13,21), indicating this is not a measurement artifact (2).

Here, plasticity is demonstrated by the 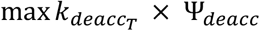 interaction (**SI Appendix Fig. S16**). As a result, throughout the winter, growing degree-day-like accumulation changes: the same temperature has its effect modulated by how much chill has accumulated at any point in winter. This has previously been reported in deacclimation rates for *Cornus sericeae* (45) and several grapevine species (24), though with much less detail. An additional contribution to plasticity can come from differences in cold hardiness. The observation that cold hardiness is the starting point from which buds progress toward budbreak, deacclimation would be accounted for by accumulation of commonly used growing degree-days, but any reacclimation (46) occurring would have to be accounted for by negation of previously accumulated growing degree-days.

Based on the inclusion of cold hardiness modeling in plasticity of plant responses, some inferences can be made about how timing of budbreak can change. Previously reported variations in budbreak timing and chill requirement for the same species (especially if same genotype) in different latitudes, or different climates (11,13,16,47), can arise from two different factors: the cold hardiness that is reached at any given location (48) (see **SI Appendix Fig. S4**) and differences in chill accumulation (8). Alternatively, when different genotypes are compared within the same environment (1,11), differences in time to budbreak can arise from different levels of cold hardiness or different rates of deacclimation. In warming experiments (6,9), higher temperature may lead to expected higher rate of deacclimation in response temperature and a lower cold hardiness during the dormant season (smaller path length), but this is somewhat balanced by lower chill accumulation (see **SI Appendix Notes**).

The objective of this work is not to produce a phenology prediction model, but rather to demonstrate how cold hardiness dynamics link the dormant and growing seasons. Therefore, no quantitative comparisons are made to other models. In addition, the equations used here are only suggested fits for the measured experimental data, rather than arbitrary expectations of what responses should be. Other models are available for prediction of cold hardiness, but those use different sets of estimates for endo- and ecodormancy phases, and are mostly based on measurements of field cold hardiness or on a few experimental determinations of temperature effects (45, 49–54). Here, no assumptions regarding the phase of bud dormancy were made. Instead, exhaustive measurement of how cold hardiness is lost through weekly determinations during two dormant seasons are used to show a quantitative progression of dormancy (i.e., deacclimation potential – *Y_deacc_*), rather than a simple division into endo- and ecodormancy phases.

Nonetheless, the parameters obtained in this study for temperature and chill responses resulted in good prediction of budbreak in the field in a single year of observation (**Fig. 9**). This demonstrates the robustness of this empirically based approach. Some error still arises from these parameters when compared to the cold hardiness path they predict (e.g., *Cercis canadensis* in **Fig. 8**). However, knowledge of the shape of responses and general magnitude of the parameters makes it possible to optimize these to produce accurate field predictions for both cold hardiness and budbreak (i.e., training of the model). It is also possible that parameters obtained here are correct, but the actual temperature experienced by buds in the field is different than that of air temperature. At night, radiative cooling can decrease bud temperature below that of air; and during the day, solar radiation may increase temperature, while wind may work to keep buds in equilibrium with air (55,56). Different effects of radiation on bud temperature may be especially relevant for studies comparing species with buds several orders of magnitude difference in size, as in the present case.

### Considerations on forcing

Linear responses to temperature for phenological responses have been widely used in the literature ever since the concept of growing degree-days was invented in the 18^th^ century (57). Here, the curvature in the response of rate of deacclimation to temperature results in residuals that are similarly skewed for most species if a linear relationship is used (as is generally the case): rates are underestimated at low temperatures and overestimated at moderate temperatures (**SI Appendix, Fig. S17**). While previous works have modeled this non-linear response as a combination of an exponential and a logarithmic (24), or as a logistic response (2), here a polynomial of third degree is used.

Non-linearity of the forcing response has recently been suggested as a source of error in studies of spring phenology (58). Based on the relationship described here in ***Eq. 1***, deacclimation rates are a relative measurement of thermal sensitivity (2,16,47,48,59). Observations of declines in thermal sensitivity (42) could therefore be arising from the seemingly exponential increase in deacclimation rates response to low temperatures, leading into a linear phase of temperature response. This nonlinearity of response to temperature in deacclimation appears to be very clear in some genotypes (e.g., *Forsythia* ‘Meadowlark’, **Fig. 6**). Measurements of deacclimation rates also show contributions towards growth of low, but above freezing temperatures that would generally not be permitted to contribute to growing degree-day calculations [below base temperature; e.g., (10)]. The non-linearity in response to temperature demonstrates that using only the change in temperature (Δ*T*; 11,12,48,60) to predict future conditions may not be appropriate, though it is possible that temperature fluctuations around where responses are over- and underpredicted may result in accurate predictions.

The combination of differential cold hardiness in the field and diversity in rates of cold hardiness loss also results in uneven effects across species. The 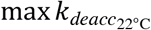 for *Forsythia* ‘Meadowlark’ is 9.5 °C day^−1^, whereas it is 0.6 °C day^−1^ for *Rhododendron calendulaceum*. This means that, in a warm winter, if all plants were 6 °C less cold hardy but still reached maximum chill accumulation, budbreak would change by less than a day for *F.* ‘Meadowlark’ while decreasing by 10 days for *R. calendulaceum* at an air temperature of 22 °C. These effects are further exacerbated at common spring temperatures: the decrease in time to budbreak with 6 °C less cold hardiness at an air temperature of 7 °C would lead to only 5 days decrease for *F.* ‘Meadowlark’ 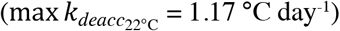 while that decrease would be 25 days for *R. calendulaceum* 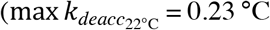 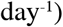 Therefore, although both species are responding to changes, their “responsiveness” measured by change in time to budbreak would be perceived as different (61).

### Including cold hardiness can improve our understanding of chilling

Chilling models are known to produce mixed results across regions. Here, the chill portions (dynamic) model (31,32) was used for convenience and based on evidence of better transferability of findings across environments (62,63). Furthermore, calculated chill portions and Utah chill units (29) have a correlation statistic higher than 0.995 in both years studied. However, many questions still remain regarding the temperatures at which chilling occurs and which therefore contribute to advancing budbreak. Most chilling accumulation models do not account for chilling at below freezing temperatures, but there is increasing evidence that they should (26,27). It is possible that in some experiments, negative temperatures have led to increases in cold hardiness [e.g., (18,22)], and thus longer paths to budbreak. Therefore, despite an increase in rate of deacclimation, the time to budbreak may not decrease. Even when negative temperatures are shown to contribute to chilling, it is possible that this effect is still underestimated because of these gains in cold hardiness.

Warm temperatures are also sometimes considered to contribute to chilling [e.g., 15 °C (17)]. In this case, their effects may be overestimated. While warmer temperatures may contribute to chill accumulation, they do promote greater loss of cold hardiness in buds than lower temperature (e.g., 11 °C vs. 2 °C in **Fig. 3B**). Therefore, they decrease the path length to observe budbreak in these assays. Determining the cold hardiness as buds are removed from chilling chambers in these experiments can assign variation in time to budbreak to increased chilling vs. decreased path length in assays.

Different temperatures have also been shown to contribute differentially in terms of chilling across species (26,27), which would suggest that models should be species-specific. In contrast, here evidence is provided that all diverse species included in the study going through dormancy at a surprisingly similar pace (**SI Appendix Fig. S12**). Different temperatures treatments imposed by Baumgarten and colleagues (27) may have had different effects in cold hardiness of each species during the chilling, which in turn could have constitute the source of interspecific variation observed. Alternatively, variations in temperature in the field around optimum temperatures for each species may have resulted in homogeneous chilling for all species studied in the present work.

Considering all three scenarios (chilling by negative temperatures, chilling by warm temperatures, and different effects across species), it is clear that coupling cold hardiness measurements of buds with chilling assays can help parse out error in models resulting in non-translatable results across regions. This demonstrates the necessity for more empirical studies of dormancy (7), though combining cold hardiness with phenology. Future experiments assessing effects of chilling treatments on cold hardiness in controlled environments or determining the extent of cold hardiness in different environments will help elucidate different chilling requirements observed across regions and years for the same species and even genotypes within species.

### Implications of a conserved dormancy progression

The pace in our understanding of plant dormancy has been very slow. The requirement of low temperatures to promote budbreak was first presented early in the 19^th^ century (64). However, to this day experiments are being conducted to understand the contributions of temperatures to chilling in order to improve chill accumulation models. This is also true in terms of the molecular understanding of dormancy: the mechanism behind this important phenotype remains elusive (33,34). Even the mechanism of chemicals known to help overcome dormancy and promote budbreak (i.e., hydrogen cyanamide) is yet to be described (35). It is possible that this is due to the fact that most studies compare species and genotypes with “low chill” and “high chill” requirement based on time to budbreak [e.g., (36–38)]. The results presented here suggest that those chill requirements are mostly a result of a genotype’s response to forcing (*r* = −0.78) and not different stages of dormancy. Therefore, genes related to those phenotypes would actually be promoting slower or faster growth [e.g., (65)].

The single dormancy transition for all plants included in the present work does not change results seen in studies of local adaptation. Instead, the findings presented here provide empirical evidence that the response to temperature, or forcing, is more important than chilling in driving perceived differences in phenology across species within the same environment (2,11,19,24,47). Environments with low chilling accumulation tend to have warmer springs, and thus the compensatory effects between chilling and forcing come into effect 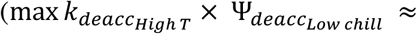 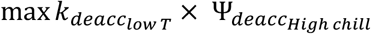, **SI Appendix, Fig. S16**). Therefore, ambient spring temperatures and rates of deacclimation must be high enough to compensate for the lack of chill accumulation. However, if chilling is too low, budbreak may take too long even at high temperatures. This can be especially important for agricultural settings where concentrated budbreak is required. However, as shown here, the absolute time to budbreak from these assays (compared to an arbitrary threshold) actually provides little information by itself in terms of the physiological stage of dormancy.

Several considerations can be made based on the sigmoid shape of the dormancy response, measured here in the form of *Y_deacc_*. The effect of chilling is quantitative, and therefore simply using qualitative determinations of endodormancy and ecodormancy (25) for buds cannot accurately describe the progression of dormancy. Although budbreak can and does occur at low chill accumulation, especially in those species with faster deacclimation rates, there is a critical state of chilling required before proper budbreak starts occurring (2) [e.g., taking almost 100 days to break bud at 22 °C (**Fig. 2**, **SI Appendix Fig. S8**) would mean forfeiting most of the growing season]. Locations with mild chill accumulation, such that they lie in the linear portion of the sigmoid curve, will be most affected by climate warming, whereas regions with high chill accumulation (8) may take much longer before decreases in chilling produce sensible results: a decrease from 100 to 75 chill portions would decrease *Y_deacc_* by 0.2, whereas a decrease from 75 to 50 chill portions reduces *Y_deacc_* by 0.4 for the average of all species in this study, resulting in uneven decreases in temperature sensitivity. Within a winter, this progression can lead to different, increasing magnitudes in the effect of unseasonal mid-winter warmings (winter weather whiplash; 66): the same amount of warming at 75 chill portions (*Y_deacc_* = 0.68) would cause more than double the deacclimation than at 50 chill portions (*Y_deacc_* = 0.25; a four to five-week difference in time at the Arnold Arboretum in both years studied). Even if these warming events do not lead to budbreak (<u>*CH*_*fiel*_</u>_*d*_<*CH_BB_*), they could leave buds at risk for “invisible” mid-winter damage should temperatures drop below cold hardiness of buds.

The same progression of dormancy for all plants observed here suggests two potential qualities for the molecular regulation of dormancy. One is that the chilling clock is an extremely conserved mechanism or an extreme case of convergent evolution (67): in the panel evaluated here, both gymnosperms and angiosperms, which diverged about 300 million years ago, are represented and showed almost no difference in deacclimation potential in response to chilling (**Fig. 5A**). The second potential aspect is that the molecular mechanism is simple in the sense that little to no variation is possible within dormancy itself. There has been some evidence presented in this sense: single or few gene changes are able to completely disrupt dormancy from plants (68,69). Instead of variation in dormancy, the results presented here suggest timing of budbreak in plants is regulated only by differences in their cold hardiness and maximum rates of cold hardiness loss. This suggests evolution acting on these traits, but not on the dormancy mechanism.

Contrary to the trend of studying plants in warmer environments, information for future adaptation was drawn here from how plants survive the cold. Using the cold hardiness dynamics phenotypes presented here, new clarity should arise from the relationships of temperature and dormancy in future studies. It is clear that temperature affects three, and not only two, aspects related to dormancy: chilling, forcing, and cold hardiness. Plants in colder climates become more cold hardy (48) and therefore have a longer path to budbreak than those in warmer climates. This status interacts with the level of chill accumulation reached prior to spring and the spring temperatures promoting forcing to result in budbreak. Using *Y_deacc_*, investigation of chilling models may yield better knowledge as to how different temperatures contribute to chilling accumulation and dormancy transitions in plants, and the overlapping temperature ranges of chilling and forcing (2). Considering the potential for consistent dormant transitions across a wide range of species, contrasts between genotypes with low and high chilling requirements may need to be reevaluated, as these likely reflect inherently slower vs. faster deacclimation or less vs. more cold hardiness, a combination of both, but not different chilling requirements. The new framework presented here should be used to further investigate cold hardiness, chilling, and forcing responses of different plant species (24), genotypes within a species (1,24), or even other taxa beyond plants (15,59). However, awareness of a single woody plant response to chill accumulation should facilitate estimation of other parameters for modeling of spring phenology. Based on the broad range of species used, the impact of cold hardiness dynamics on phenology is likely universal across plants presenting cold season dormancy: a keystone phenotype defining the dormant season that has been largely ignored but must be included in future studies.

## Materials and Methods

### Study site and material collected

All material was collected from plants from the Arnold Arboretum of Harvard University, located in Boston, MA, USA (42°17’57” N 71°07’22” W). The climate is a humid subtropical climate (Koeppen’s Cfa), USDA Plant Hardiness Zone 6b [average annual extreme minimum temperature −17.8 to −15 °C (70)]. The 15 species collected were balsam fir (*Abies balsamea*), red maple (*Acer rubrum*), sugar maple (*Acer saccharum*), Eastern redbud (*Cercis canadensis*), flowering dogwood (*Cornus florida*), cornelian cherry (*Cornus mas*), American beech (*Fagus grandifolia*), forsythia (*Forsythia* × ‘Meadowlark’), mountain laurel (*Kalmia latifolia*), Japanese larch (*Larix kaempferi*), dawn redwood (*Metasequoia glyptostroboides*), Norway spruce (*Picea abies*), apricot (*Prunus armeniaca* ‘Mikado’), Canadian plum (*Prunus nigra*), and flame azalea (*Rhododendron calendulaceum*). Buds were collected from individual plants, except for: *Forsythia* × ‘Meadowlark’, for which multiple clones are available; and *Fagus grandifolia*, *Kalmia latifolia* and *Rhododendron calendulaceum*, which resulted in collections from multiple individuals at random located in the same mass due to large number of buds required. Plant IDs are available in **Extended Data Table 1**, and further details can be found using the Arboretum Explorer website (https://arboretum.harvard.edu/explorer/).

Collections for deacclimation and budbreak assays occurred at weekly intervals in the field during two dormant seasons: from mid-September 2019 through mid-March 2020, and mid-September 2020 through end of February 2021 (**Extended Data Table 2**) and were processed on the same day of collection. Cuttings were separated in two sets: 10 cuttings from each species were separated for budbreak observations, whereas the additional material was used for cold hardiness measurements. The cuttings were placed in a growth chamber at 22 °C, 16h daylight in cups of water; the water was changed weekly. In the first season, assessments of field cold hardiness continued between mid-March 2020 and budbreak for each species, but lab access restrictions due to COVID-19 did not allow for deacclimation and budbreak assays. Additional cuttings were collected in mid-winter [mid-February (82 chill portions) in the first season, mid-January (54 chill portions) in the second season], from 13 of the 15 species in both years combined (all but *Acer saccharum* and *Metasequoia glyptostroboides*) to investigate the effect of different temperatures in deacclimation. The cuttings were also placed in cups of water, and into growth chambers at seven different temperatures (2, 4, 7, 11, 15, 22 and 30 °C) to establish the relationship of deacclimation rate to temperature. With the exception of the 22 °C treatment (16h daylight), all other temperatures were combined with 0h daylight [see (24) for similar description of methods].

### Cold hardiness measurements

Field cold hardiness was measured on the same day of collections. The material in the growth chambers (at 22 °C from weekly collections and at different temperatures for two collections) was then used to evaluate loss of cold hardiness (deacclimation) over time in semi-regular intervals. Cadence of sampling to determine rates of deacclimation was based on equipment availability and expected deacclimation rates for each species. In general, a minimum number of time points for cold hardiness measurements were sought such as to determine the two parameters of the linear relationship: species with higher deacclimation rates (e.g., Forsythia × ‘Meadowlark’ and *Prunus armeniaca*) were generally sampled at smaller intervals than those with lower deacclimations rates (e.g., *Picea abies*). However, at high chill accumulations, extreme rates of deacclimation in some species led to full deacclimation within a day, and therefore rates being based on only two time points [field (day 0) and day 1]. For the temperature experiment, cold hardiness was generally assessed at smaller intervals in higher temperatures than lower temperatures. Measurements for all species at 22 °C in every collection are available in **SI Appendix Figs. S18 to S47**. For temperature responses, measurements are available in **SI Appendix Figs. S48 and S49**.

Cold hardiness of buds was measured using differential thermal analysis [DTA; (71)]. In DTA measurements, high temperature exotherms (HTEs) represent non-lethal freezing events and low temperature exotherms (LTEs) represent lethal freezing within buds. Buds were excised from stems and placed in thermoelectric modules (TEMs) in plates connected to a Keithley data logger (Tektronix, Beaverton, OR). The plates were placed in a programmable environmental chamber (Tenney, New Columbia, PA), in which temperature was decreased to −5 °C for 5h to promote HTEs and then cooled at −4 °C hour^−1^ to −55 °C. Due to COVID-19 related restrictions, starting 19 March 2020 until the end of the first season, the freezing was done in a common household freezer, and therefore the rate of cooling was not controlled and the minimum temperature reached was approximately −22 °C. Temperature and changes in voltage due to temperature differentials between the two surfaces of the TEMs are recorded, and therefore the release of heat due to freezing of supercooled water in buds results in voltage peaks. A maximum of 10 values for cold hardiness were recorded each time. Many species became less cold hardy in the field in mid-March 2020. While it is possible that the lower limit of detection (at approximately –22 °C in a household freezer) caused these errors, this shift also overlaps with considerable increase in air temperature and even full deacclimation and budbreak of some species.

### Budbreak

Observations for budbreak in the 22 °C chamber occurred daily and was noted when budbreak occurred in any bud in cuttings with multiple buds (all species except *Cornus florida* and *Rhododendron calendulaceum*, for which cuttings with single buds were used). Generally, in early collections this resulted in a single bud breaking in cuttings with multiple buds, whereas at later collections multiple buds would break synchronously within the same cutting – resulting in an underestimation in overall time to budbreak at low chill accumulations. Due to differences in bud morphology for each species, the budbreak phenotype was different and is further described in **Extended Data Table 1**. In general, very early signs of budbreak were used (e.g., bud scales begin to separate and leaf tips are visible) rather than leafout. In species where both vegetative and reproductive budbreak was observed, vegetative budbreak comprised most of the early observations, whereas at later collections budbreak was mostly of reproductive buds. Budbreak was also observed in the field in the spring of 2020 (**SI Appendix Fig. S50**). Observations were done in daily intervals and the same phenotype for budbreak was used as that of controlled environment observations, noted as percent of total buds presenting the phenotype.

### On-site meteorological data

Weather data were obtained from the Weld Hill meteorological station (HOBO RX3000 Station, Hobolink), available through the Arnold Arboretum website (arboretum.harvard.edu). The most distant plant (*Acer rubrum*) is located ~1,700 m from the weather station. Hourly air temperature was used to compute chilling accumulation at each collection point. Chilling accumulation was calculated as chill portions based on the Dynamic Model (31,32). Temperature and chilling accumulation are shown in **SI Appendix Fig. S7**.

### Statistical analysis

All analyses were conducted and figures produced using R [v. 4.0.4 (72)] within RStudio [v. 1.3.1093 (73)] and multiple packages therein (74–82). Deacclimation rates were determined using ordinary linear regressions, where cold hardiness is the dependent variable, and time in days is the independent variable for each species in each collection date:

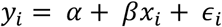

where the intercept *a* represents the cold hardiness at collection, and regression slope *b* is the effective deacclimation rate 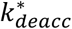 in °C day^−1^. Because of detection limits at above ~−3 °C (**SI Appendix Fig. S17**), and the general trend of cold hardiness measurements to have a minimum cold hardiness of −5 °C (18), adjacent timepoints with similar cold hardiness measurements at warmer temperatures were manually removed to prevent reductions in the deacclimation rates due to this artifact. Datapoints with studentized residuals greater than 3 were removed (1.3% of measurements) and regressions were re-fit. Some gains in cold hardiness were observed in earlier collection points, which is expected to be a result of buds maturing within the warm temperature chamber.

To compare the effect of chilling among species, the rates of deacclimation from weekly collections subjected to 22 °C were normalized within each genotype, resulting in the potential for deacclimation at each collection:

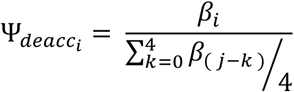

where b_i_ is the deacclimation rate at the *i*^th^ collection, and b_(j)_ is the *j^th^* highest deacclimation rate, such that the rate is normalized to the average of the highest 4 measured deacclimation rates. The average of the 4 highest deacclimation rates is also presented as 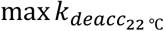 A three-parameter log-logistic function, with upper limit 1, was fit to these data:

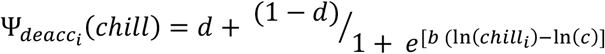

where the potential for deacclimation at the *i^th^* collection is a function of chilling accumulation at the *i^th^* collection. Three estimates are thus obtained: *d* is the minimum deacclimation observed at all times (analogous to an intercept); *b* is the slope associated with the log-logistic curve, where negative values indicate increases in *Y_deacc_* as chill accumulation increases, and greater magnitudes indicate a higher slope; and *c* is the inflection point of the log-logistic curve. To establish chilling requirements based on time to 50% budbreak at 22 °C, the threshold of 15 days was used. The chilling requirement was calculated as the mean chilling accumulation of the four consecutive points where the average time to budbreak was closest to 15 days, using both years of data combined. For species in which the 15 days threshold was not crossed, the last four timepoints were used.

The maximum cold hardiness (*CH_max_*) of each species was determined as the average of measurements from the four dates where the cold hardiness for each genotype were maximum. To estimate the cold hardiness at budbreak (*CH_BB_*), the field cold hardiness dataset was merged with the budbreak dataset, such that each cold hardiness measurements had n budbreak times associated with it – n being the number of buds that broke within a collection. This dataset was subset using a threshold of >25 chill portions accumulated in order to select informative points where deacclimation rates were high enough and budbreak occurred with more regularity. Each cold hardiness point was then predicted to lose cold hardiness at the measured deacclimation rate until each time of budbreak (see **SI Appendix Fig. S1**). The average of all measurements then provided an estimated parameter for cold hardiness at budbreak. Because *Rhododendron calendulaceum* had very poor break bud in controlled environment, a determined cold hardiness at budbreak was used for this species based on literature [*CH_BB_* = −5 °C (46,83)].

The response of deacclimation rate to temperature was fit using a third-degree polynomial to acknowledge the fast increases in rate at low temperatures and lower increases in rate at high temperatures for the nine species in which different temperatures were tested. The slopes from each temperature for each species were corrected based on the ratio of the rate measured at 22 °C at this time and 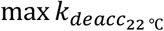 (**Extended Data Fig. 4a**). Corrected rates were then used to calculate the function that describes *k_deacc_* for each species:

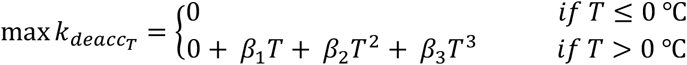

where T is the air temperature. For those genotypes in which the response to different temperatures was not estimated, a linear response was used with a 0 intercept; and 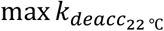 divided by 22 as b_1_.

To evaluate time to budbreak in relation to field cold hardiness, a rate of acclimation (*k_acc_*) was required. The linear rate of acclimation used was:

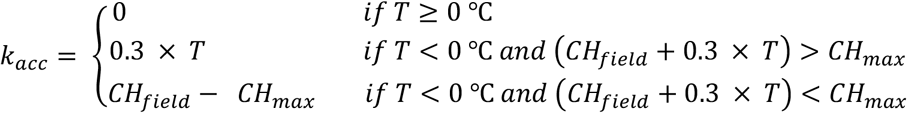

where acclimation does not occur at above 0 °C, and is linear otherwise. A threshold was imposed such that the predicted cold hardiness would not surpass maximum measured cold hardiness. The estimate of 0.3 was used based on three iterations (0.1, 0.2, 0.3) for acclimation rates of similar magnitude to those in the literature (49,50).

Estimates, sample sizes (n), and appropriate fitness statistics (e.g., *r^2^*s and *P*-values) for each portion of the data analyses are available in **SI Appendix Tables S3–S5**.

## Notes

https://www.github.com/apkovaleski/CH_Budbreak_ArnoldArb

### Competing Interest Statement

The authors have declared no competing interest.

## References

1. E. Thibault, R. Soolanayakanahally, S. R. Keller, Latitudinal clines in bud flush phenology reflect genetic variation in chilling requirements in balsam poplar, Populus balsamifera. Am. J. Botany 107, 1–9 (2020).

2. I. Chuine, A unified model for budburst in trees. J. theor. Biol. 207, 337–347 (2000).

3. C. J. Chamberlain, B. I. Cook, I. Morales-Castilla, E.M. Wolkovich, Climate change reshapes the drivers of false spring risk across European trees. New Phyt. 229, 323–334 (2020).

4. L. Gu et al., The 2007 Eastern US spring freeze: increased cold damage in a warming world? BioScience 58:253–262.

5. C. J. Chamberlain, B. I. Cook, I. G. Cortázar-Atauri, E. M. Wolkovich, Rethinking false spring risk. Glob. Chang. Biol. 25, 2209–2220 (2019).

6. E. M. Wolkovich et al., Warming experiments underpredict plant phenological responses to climate change. Nature 485, 494–497 (2012).

7. H. Hanninen et al., Experiments are necessary in process-based tree phenology modeling. Trends in Plant Science 24, 199–209 (2019).

8. E. Luedeling, E. H. Girvertz, M. A. Semenov, P. H. Brown, Climate change affects winter chill for temperate fruit and nut trees. PLOS ONE 6, e20155 (2011).

9. A. D. Richardson et al., Ecosystem warming extends vegetation activity but heightens vulnerability to cold temperatures. Nature 560, 368–371 (2018).

10. R. A. Montgomery, K. E. Rice, A. Stefanski, R. L. Rich, P. B. Reich, Phenological responses of temperate and boreal trees to warming depend on ambient spring temperatures, leaf habit, and geographic range. Proc. Natl. Acad. Sci. U.S.A. 117, 10397–10405 (2020).

11. A. K. Ettinger et al., Winter temperatures predominate in spring phenological responses to warming. Nature Climate Change 10, 1137–1142 (2020).

12. L. Meng et al., Urban warming advances spring phenology but reduces the response of phenology to temperature in the conterminous United States. Proc. Nat. Acad. Sci. U.S.A. 117, 4228–4233 (2020).

13. C. Bigler, Y. Vitasse, Daily maximum temperatures induce lagged effects on leaf unfolding in temperate woody species across large elevational gradients. Front. Plant Sci. 10, 398 (2019).

14. T. F. Keenan, A. D. Richardson, K. Hufkens, On quantifying the apparent temperature sensitivity of plant phenology. New Phyt. 225, 1033–1040 (2020).

15. D. N. Laskin et al., Advances in phenology are conserved across scale in present and future climates. Nature Climate Change 9, 419–425 (2019).

16. B. Wenden, M. Mariadassou, F.-M. Chmielewski, Y. Vitasse, Shifts in the temperature-sensitive periods for spring phenology in European beech and pedunculate oak clones across latitudes and over recent decades. Global Change Biol. 26, 1808–1819 (2020).

17. D. F. B. Flynn, E. M. Wolkovich, Temperature and photoperiod drive spring phenology across all species in a temperate forest community. New Phyt. 219, 1353–1362 (2018).

18. A. Lenz, G. Hoch, Y. Vitasse, C. Körner, European deciduous trees exhibit similar safety margins against damage by spring freeze events along elevational gradients. New Phyt. 200, 1166–1175 (2013).

19. J. D. Fridley, Extended leaf phenology and the autumn niche in deciduous forest invasions. Nature 485, 359–362 (2012).

20. M. D. Schwartz, R. Ahas, A. Aasa, Onset of spring starting earlier across the northern hemisphere. Glob. Change Biol. 12, 343–351 (2006).

21. H. Wang et al., Overestimation of the effect of climatic warming on spring phenology due to misrepresentation of chilling. Nature Communications 11, 4945 (2020).

22. A. Vitra, A. Lenz, Y. Vitasse, Frost hardening and dehardening potential in temperate trees from winter to budburst. New Phyt. 216, 113–123 (2017).

23. J. P. Londo, A. P. Kovaleski, Characterization of wild North American grapevine cold hardiness using differential thermal analysis. Am. J. Enol. Vitic. 68, 203–212 (2017)

24. A. P. Kovaleski, B. I. Reisch, J. P. Londo, Deacclimation kinetics as a quantitative phenotype for delineating the dormancy transition and thermal efficiency for budbreak in *Vitis* species. AoB PLANTS 10, ply066 (2018).

25. G. A. Lang, J. D. Early, G. C. Martin, R. L. Darnell, Endo-, para-, and ecodormancy: physiological terminology and classification for dormancy research. HortScience 22, 371–377 (1987).

26. J. Cragin, M. Serpe, M. Keller, K. Shellie, Dormancy and cold hardiness transitions in winegrape cultivars Chardonnay and Cabernet Sauvignon. Am. J. Enol. Vitic. 68, 195–202 (2017).

27. F. Baumgarten, C. M. Zohner, A. Gessler, Y. Vitasse, Chilled to be forced: the best dose to wake up buds from winter dormancy. New Phyt. 230, 1366–1377 (2021).

28. J. Bennet, Temperature and bud rest period: Effect of temperature and exposure on the rest period of deciduous plant leaf buds investigated. California Agriculture 4, 11–16 (1949).

29. E. A. Richardson, S. D. Seeley, D. R. Walker, A model for estimating the completion of rest for ‘Redhaven’ and ‘Elberta’ peach trees. HortScience 9, 331–332 (1974).

30. A. D. Shaltout, C. R. Unrath, Rest completion prediction model for Starkrimson Delicious apples. J. Am. Soc. Hort. Sci. 108, 957–961 (1983).

31. S. Fishman, A. Erez, G. A. Couvillon, The temperature dependence of dormancy breaking in plants: mathematical analysis of a two-step model involving a cooperative transition. J. Theo. Biol. 124, 473–483 (1987).

32. S. Fishman, A. Erez, G. A. Couvillon, The temperature-dependence of dormancy breaking in plants – computer-simulation of processes studied under controlled temperatures. J. Theo. Biol. 126, 309–321 (1987).

33. J. E. K. Cooke, M. E. Eriksson, O. Junttila, The dynamic nature of bud dormancy in trees: environmental control and molecular mechanisms. Plant, Cell & Envir. 35, 1707–1728 (2012)

34. H. Yamane, A. K. Singh, J. E. K. Cooke, Plant dormancy research: from environmental control to molecular regulatory networks. Tree Phys. 41, 523–528 (2021)

35. I. A. Ionescu et al., Transcriptome and metabolite changes during hydrogen cyanamide-induced floral bud break in sweet cherry. Front. Plant Sci. 8, 1233 (2017).

36. Y. E. Miotto et al., Spring is coming: genetic analysis of the bud break date locus reveal candidate genes from the cold perception pathway to dormancy release in apple (*Malus* x *domestica* Borkh.). Front. Plant Sci. 10, 33 (2019).

37. N. Vimont et al., From bud formation to flowering: transcriptomic state defines the cherry developmental phases of sweet cherry bud dormancy. BMC Genomics 20, 974 (2019).

38. D. D. Porto et al., Transcription profiling of the chilling requirement for bud break in apples: a putative role for *FLC*-*like* genes. J. Exp. Bot. 66, 2659–2672 (2015).

39. Y. Vitasse, D. Basler, Is the use of cuttings a good proxy to explore phenological responses of temperate forests in warming and photoperiod experiments? Tree Phys. 34, 174–183 (2014).

40. G. Neuner, K. Monitzer, D. Kaplening, J. Ingruber, Frost survival mechanism of vegetative buds in temperate trees: deep supercooling and extraorgan freezing vs. ice tolerance. Front. Plant Sci. 10, 537 (2019).

41. H. Yu, E. Luedeling, J. Xu, Winter and spring warming result in delayed spring phenology on the Tibetan Plateau. Proc. Natl. Acad. Sci. U.S.A. 51, 22151–22156 (2010)

42. Y. H. Fu et al., Declining global warming effects on the phenology of spring leaf unfolding. Nature 526, 104–107 (2015).

43. A. Viherä-Aarnio, S. Sutinen, J. Partanen, R. Häkkinen, Internal development of vegetative buds of Norway spruce trees in relation to accumulated chilling and forcing temperatures. Tree Phys. 34, 547–556 (2014).

44. A. P. Kovaleski, J. P. Londo, K. D. Finkelstein, X-ray phase contrast imaging of *Vitis* spp. buds shows freezing pattern and correlation between volume and cold hardiness. Sci. Reports 9, 1–12 (2019).

45. K. D. Kobayashi, L. H. Fuchigami, C. J. Weiser, Modeling cold hardiness of red-osier dogwood. J. Amer. Soc. Hort. Sci. 108, 376–381 (1983).

46. S. R. Kalberer, N. Leyva-Estrada, S. L. Krebs, R. Arora, Frost dehardening and rehardening of floral buds of deciduous azaleas are influenced by genotypic biogeography. Env. Exp. Bot. 59, 264–275 (2007).

47. X. Geng et al., Climate warming increases spring phenological differences among temperate trees. Glob. Change Biol. 26, 5979–5987 (2020).

48. A. Lenz, G. Hoch, Y. Vitasse, Fast acclimation of freezing resistance suggests no influence of winter minimum temperature on the range limit of European beech. Tree Phys. 36, 490–501 (2016).

49. J. C. Ferguson, J. M. Tarara, L. J. Mills, G. G. Grove, M. Keller, Dynamic thermal time model of cold hardiness for dormant grapevine buds. Ann. Bot. 107, 389–396 (2011).

50. J. C. Ferguson, M. M. Moyer, L. J. Mills, G. Hoogenboom, M. Keller, Modeling dormant bud cold hardiness and budbreak in twenty three Vitis genotypes reveals variation by region of origin. Am. J. Enol. Vitic. 65, 59–71 (2014).

51. S. Kellomäki et al., A simulation model for the succession of the boreal forest ecosystem. Silva Fennica 26, 1–18 (1992)

52. H. Hänninen, Climate warming and the risk of frost damage to boreal forest trees: identification of critical ecophysiological traits. Tree Phys 26, 889–898 (2006)

53. I. Leinonen, A simulation model for the annual frost hardiness and freeze damage of scots pine. Ann. Bot. 78, 687–693 (1996).

54. I. Leinonen, R. Repo, H. Hänninen, K. E. Burr, A second-order dynamic model for the frost hardiness of trees. Ann. Bot. 76, 89–95 (1995).

55. A. J. P. Quiñones, M. Keller, M. R. S. Gutierrez, L. Khot, G. Hoogenboom, Comparison between grapevine tissue temperature and air temperature. Sci. Hort. 247, 407–420 (2018).

56. J. Grace, The temperature of buds may be higher than you thought. New Phyt. 170, 1–3 (2006).

57. M. Reaumur. Observations du thermometre, faites à Paris pendant l’année 1735, comparées avec celles qui ont été faites sous la Ligne, à l’Isle de France, à Alger, & en quelques-unes de nos Isles de l’Amérique. Memoires de l’Academie Royale des Sciences 1735, 545–576 (1735).

58. E. M. Wolkovich et al., A simple explanation for declining temperature sensitivity with warming. bioRXiv [Preprint] (2021). https://doi.org/10.1101/2021.01.12.426288 (accessed 25 April 2021).

59. S. J. Thackeray et al., Phenological sensitivity to climate across taxa and trophic levels. Nature 535, 241–245 (2016).

60. W. Buermann et al., Widespread seasonal compensation effects of spring warming on northern plant productivity. Nature 562, 110–115 (2018).

61. C. Parmesan, Influences of species, latitudes and methodologis on estimates of phenological response to global warming. Glob. Chang. Biol. 13, 1860–1872 (2007).

62. E. Luedeling, Climate change impacts on winter chill for temperate fruit and nut production: a review. Sci. Hort. 144, 218–229 (2012).

63. J. Zhang, C. Taylor, The dynamic model provides the best description of the chill process on ‘Sirora’ pistachio trees in Australia. HortScience 46, 420–425 (2011).

64. T. A. Knight, XV. Account of some experiments on the ascent of the sap in trees. In a letter from Thomas Andrew Knight, Esq. to the Right Hn. Sir Joseph Banks, Bart. K. B. P. R. S. Philos. Trans. R. Soc. Lond. 91: 333–353 (1801).

65. A. P. Kovaleski, J. P. Londo, Tempo of gene regulation in wild and cultivated *Vitis* species shows coordination between cold deacclimation and budbreak. Plant Sci. 287, 110178 (2019).

66. N. J. Casson et al., Winter weather whiplash: impacts of meteorological events misaligned with natural and human systems in seasonally snow-covered regions. Earth’s Future 7, 1434–1450 (2019).

67. S. Yeaman et al., Convergent local adaptation to climate in distantly related conifers. Science 353, 1431–1433 (2016).

68. S. Tylewicz et al., Photoperiodic control of seasonal growth is mediated by ABA acting on cell-cell communication. Science 360, 212–215 (2018).

69. D. G. Bielenberg et al., A deletion affecting several gene candidates is present in the *Evergrowing* peach mutant. J. Hered. 95, 436–444 (2004).

70. M. S. Dosmann, The history of minimum temperatures at the Arnold Arboretum: variation in time and space. Arnoldia 72, 2–11 (2015).

71. L. J. Mills, J. C. Ferguson, M. Keller, Cold-hardiness evaluation of grapevine buds and cane tissues. Am. J. Enol. Vitic. 57, 194–200 (2006).

72. R Core Team. R: A language and environment for statistical computing. R Foundation for Statistical Computing, Vienna, Austria (2020).

73. RStudio Team. RStudio: Integrated Development Environment for R. RStudio, PBC, Boston, MA (2020).

74. T. L. Pedersen, patchwork: the composer of plots. R package version 1.1.0. https://CRAN.R-project.org/package=patchwork (2020).

75. H. Wickham, ggplot2: elegant graphics for data analysis. Springer-Verlag New York (2016).

76. C. Ritz, F. Baty, J. C. Streibig, D. Gerhard, Dose-Response Analysis Using R. PLOS ONE, 10, e0146021 (2015).

77. E. Luedeling, chillR: Statistical Methods for Phenology Analysis in Temperate Fruit Trees. R package version 0.70.24. https://CRAN.R-project.org/package=chillR (2020).

78. G. Grolemund, H. Wickham, Dates and times made easy with lubridate. J. Stat. Softw. 40, 1–25 (2011).

79. J. Fox, S. Weisberg, An R companion to applied regression. Third Edition. Thousand Oaks CA: Sage (2019).

80. W. N. Venables, B. D. Ripley, Modern applied statistics with S. Fourth Edition. Springer, New York. ISBN 0-387-95457-0 (2002).

81. F. Mendiburu, agricolae: Statistical Procedures for Agricultural Research. R package version 1.3–3 (2020).

82. C. Sievert, Interactive Web-Based Data Visualization with R, plotly, and shiny. Chapman and Hall/CRC Florida (2020).

83. G. Neuner, D. Ambach, O. Buchner, Readiness to frost harden during the dehardening period measured *in situ* in leaves of *Rhododendron ferrugineum* L. at the alpine timberline. Flora 194, 289–296.

