## Supplementary material for "Woody species do not differ in dormancy progression: differences in time to budbreak due to forcing and cold hardiness": SI Appendix Notes

**Supporting Information Notes**

**Integrating cold hardiness and budbreak**

Throughout the work presented, the idea of a connection between spring phenology as budbreak and cold hardiness dynamics is introduced. The two aspects are shown to be mathematically linked in many species, which suggests a new way of studying dormancy that may bring clarity to this elusive aspect of perennial plants. Because of the novelty of the parameters shown, here I present examples of how each of the parameters related to cold hardiness can affect the universally measured time to 50% budbreak (BB_50_). The hypothetical examples presented are cases explored in relation to the following **Eq. 1** and **Eq. 4** presented in the main text:

$$Time to Budbreak=\frac{\left| {CH}_{0}-{CH}_{BB} \right|}{k_{deacc}^{*}}$$

$$k_{{deacc}_{T}}^{*}=\max k_{{deacc}_{T}} \times\Psi_{deacc}$$

where *CH_0_* is the initial cold hardiness (referred to in the main text as the “departure point”), *CH_BB_* is the cold hardiness at budbreak, $k_{deacc}^{*}$ is the effective rate of deacclimation. $k_{{deacc}_{T}}^{*}$ is the effective rate of deacclimation at temperature T, which is a result of the product of the maximum rate of deacclimation at temperature T ($\max k_{{deacc}_{T}}$) and the deacclimation potential ($\Psi_{deacc}$). The cold hardiness at budbreak (*CH_BB_*) is estimated based on the estimated cold hardiness from the deacclimation assay at the time when 50% budbreak is observed (**SI Appendix Fig. S1**).

1. **Differing rates of deacclimation, with constant initial cold hardiness and cold hardiness at budbreak**

When comparing two samples that have the same initial cold hardiness (*CH_0_*) and same cold hardiness at budbreak (*CH_BB_*), this means the path length is the same for these samples (**SI Appendix Fig. S2**). The path length (in °C) is a measurement of how much cold hardiness must be lost before budbreak is observed. In this example, two different effective rates of deacclimation ($k_{deacc}^{*}$) are presented (the slopes of the lines), where the rate is higher in “***A***” than in “***B***”. What occurs, in terms of visible phenotype, is that budbreak in “***A***” is observed earlier than in “***B***”. In addition, the duration of budbreak is shorter, or more concentrated, in “***A***” than in “***B***”. This is a result of the “shallower angle” with which the buds deacclimating in “***B***” reach the cold hardiness at budbreak.

2. **Different cold hardiness at budbreak, with constant rate of deacclimation and initial cold hardiness**

In this example, initial cold hardiness and rates of deacclimation are kept constant, but cold hardiness for budbreak is greater in “***B***” than in “***A***” (**SI Appendix Figs. S3A** and **S3B**). It is important to note that greater cold hardiness means a more negative value, so the greater is in a magnitude sense. This being the case, the greater CH_BB_ leads to a shorter path length to budbreak. Therefore, budbreak is observed later in “***A***” compared to “***B***” (**SI Appendix Fig. S3C**), although the distribution of budbreak is similar.

3. **Different initial cold hardiness, at different levels of chill accumulation, with constant rate of deacclimation and cold hardiness at budbreak**

Here, two samples are presented at two different initial cold hardiness (*CH_0_*; where “***A***” is less cold hardy than “***B***”), while the cold hardiness at budbreak (*CH_BB_*) is kept constant (**SI Appendix Figs. S4A** and **S4B**). In addition the rate of deacclimation ($k_{deacc}^{*}$) is kept constant for the same color curves, where colors represent different levels of chill accumulation. The more cold hardy sample thus takes longer to budbreak (**SI Appendix Figs. S4C**) at any chill accumulation than the less cold hardy sample. By looking at the time to budbreak (**SI Appendix Figs. S4C**), it appears as if “***B***” is decreasing the time to budbreak faster than “***A***” as chill accumulates. The distance in time to budbreak at any given point between “***A***” and “***B***” is the difference in cold hardiness between the samples divided by the rate of deacclimation, which explains the seemingly asymmetrical decrease in time to budbreak. However, when the rates of deacclimation are observed in relation to chill accumulation, no difference is observed between the two samples (**SI Appendix Figs. S4D**). This suggests that the physiological state in response to chilling is the same for both, but presumably lower temperatures experienced by “***B***” led to greater gain of cold hardiness in the field prior to hypothetical deacclimation assays compared to “***A***”.

4. **Different rates of deacclimation, at different levels of chill accumulation, with constant initial cold hardiness and cold hardiness at budbreak**

In this example, the initial cold hardiness and cold hardiness at budbreak are kept constant, but the effective rates of deacclimation ($k_{deacc}^{*}$) at the same levels of chill accumulation (same colors) are greater in “***A***” compared to “***B***” (**SI Appendix** **Figs. S5A** and **S5B**). Because of the lower rates, “***B***” shows a longer time to budbreak at same levels of chill accumulation (same colors) compared to “***A***” (**SI Appendix Fig. S5C**). If we look at the effective rates of deacclimation in response to chilling (**SI Appendix Fig. S5D**), “**b**” has lower rates of deacclimation at the same chill accumulations compared to “***A***”. However, if the rates are standardized based on the highest rate measured at high chill accumulation (shown in red in **SI Appendix Fig. S5**), we obtain the deacclimation potential at any point in chill accumulation (**SI Appendix Fig. S5E**). This suggests that again the physiological state of the two samples at these points in chill accumulation are the same, but one species (or genotype) is inherently faster than the other.

5. **Considerations**

Here the examples in 1-4 describe changes in single parameters at a time to simplify the effects observed in time to budbreak. However, most parameters will differ for different species (*CH_0_*, *CH_BB_*, and $\max k_{{deacc}_{T}}$), and possibly for genotypes within species, though it appears that $\Psi_{deacc}$ is very similar across distantly related species. This suggests that the regulation of dormancy in all plants is very conserved, and perhaps a simple process such that any changes in regulation lead to severe maladaptation. Also, while *CH_BB_* and $\max k_{{deacc}_{T}}$ are species dependent, *CH_0_* will vary depending on the lowest temperatures experienced in any given year or region, or the time of collection within a dormant season (see **SI Appendix Fig. S6**).
