## Supplementary material for "Woody species do not differ in dormancy progression: differences in time to budbreak due to forcing and cold hardiness": Fig. S

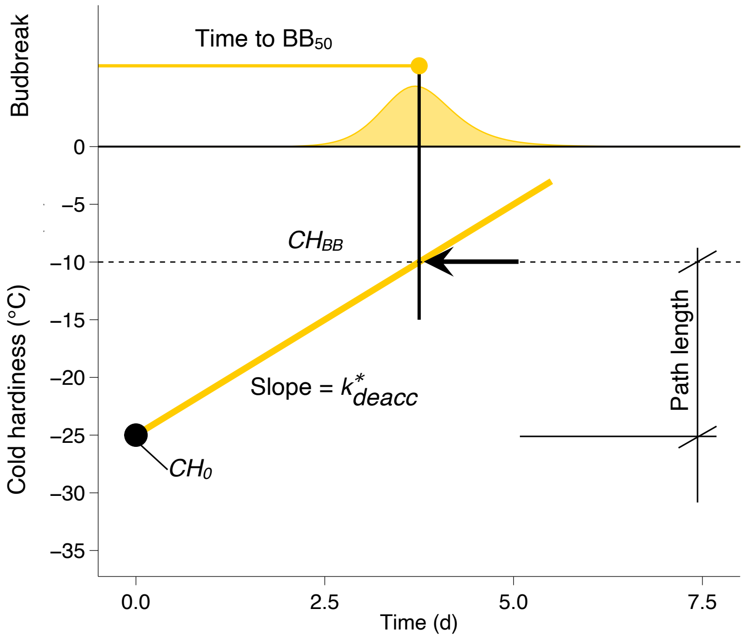


**Supporting Information Figure S1. Schematic view of variables used in this study.** The initial cold hardiness ($CH_0$) is the cold hardiness of buds collected from the field. The effective rate of deacclimation $k^*_{deacc}$ is the slope of the deacclimation of buds in controlled environment. The time to 50% budbreak ($BB_{50}$) is measured in the same controlled environment as deacclimation. Cold hardiness at budbreak ($CH_{BB}$) is estimated based on the cold hardiness estimate when $BB_{50}$ occurs. The path length is the distance in °C from $CH_0$ to $CH_{BB}$.

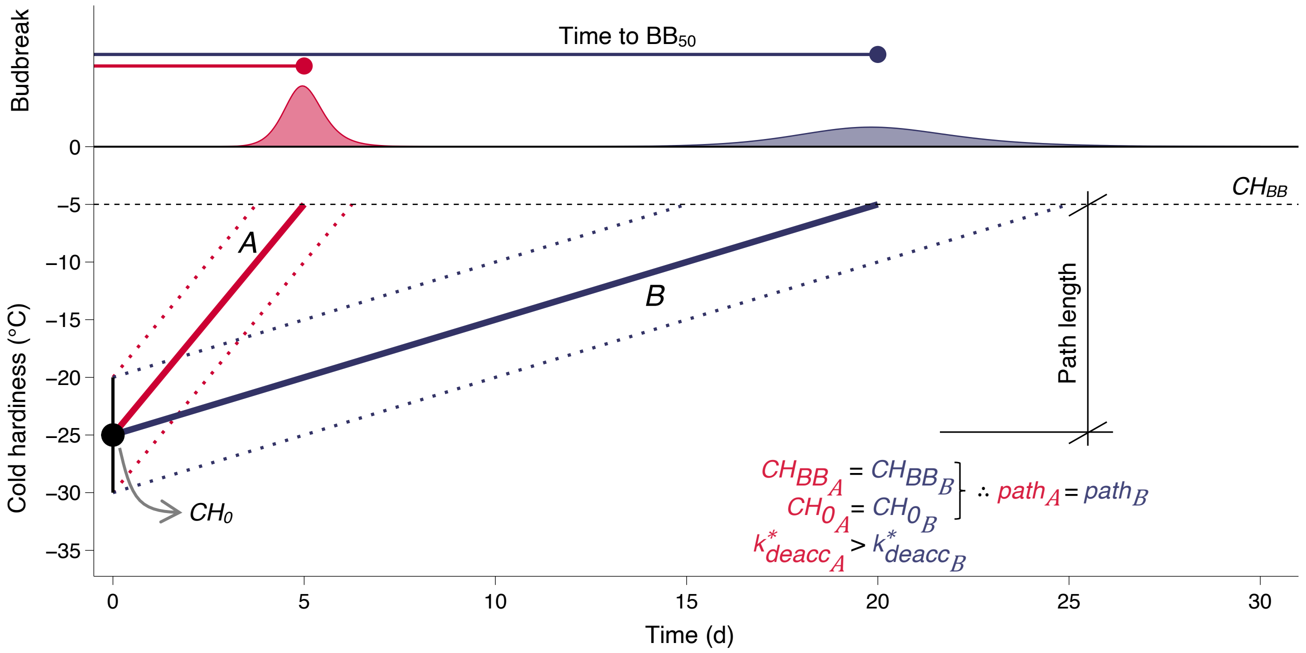


**Supporting Information Figure S2. Different rates of deacclimation.** Without varying initial cold hardiness ($CH_{0_A} = CH_{0_B}$) or cold hardiness at budbreak ($CH_{BB_A} = CH_{BB_B}$), the path length remains the same ($path_A = path_B$). However, as rate of deacclimation in *A* ($k^*_{deacc_A}$) is greater than in *B* ($k^*_{deacc_B}$), 50% budbreak ($BB_{50}$) is earlier and more concentrated in time.

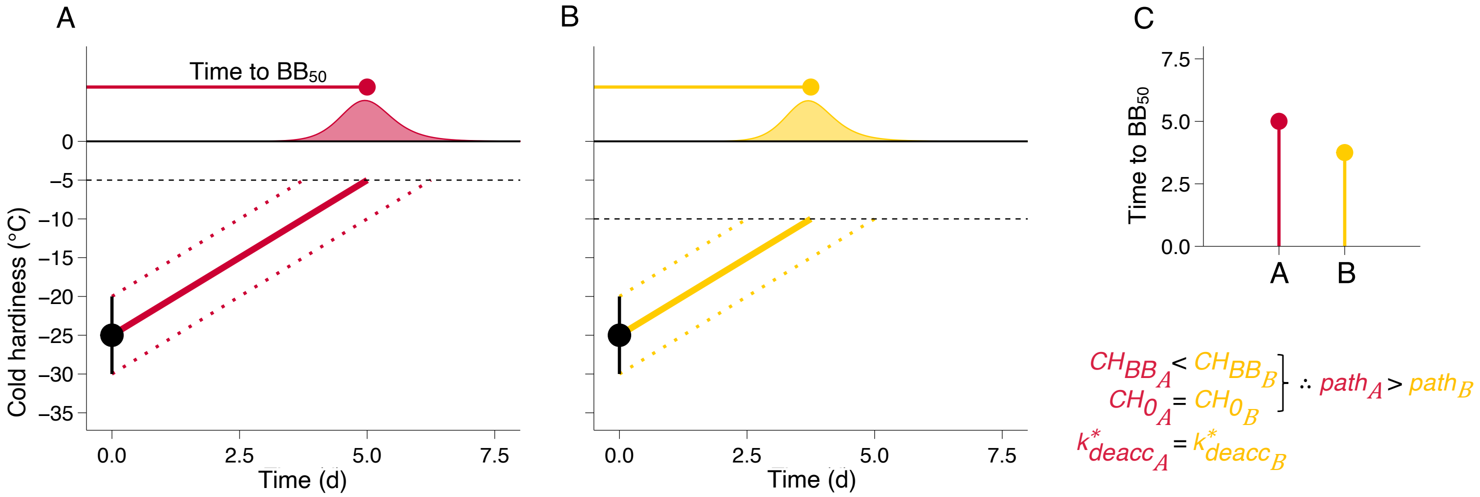


**Supporting Information Figure S3. Different cold hardiness at budbreak.** Without varying initial cold hardiness ($CH_{0_A} = CH_{0_B}$) but having *A* be less cold hardy at budbreak than *B* ($CH_{BB_A} < CH_{BB_B}$), the path length is greater in *A* ($path_A > path_B$). Therefore, even as the rates of deacclimation are the same ($k^*_{deacc_A} = k^*_{deacc_B}$), there is a perceived difference in time to 50% budbreak between the two *(C).*

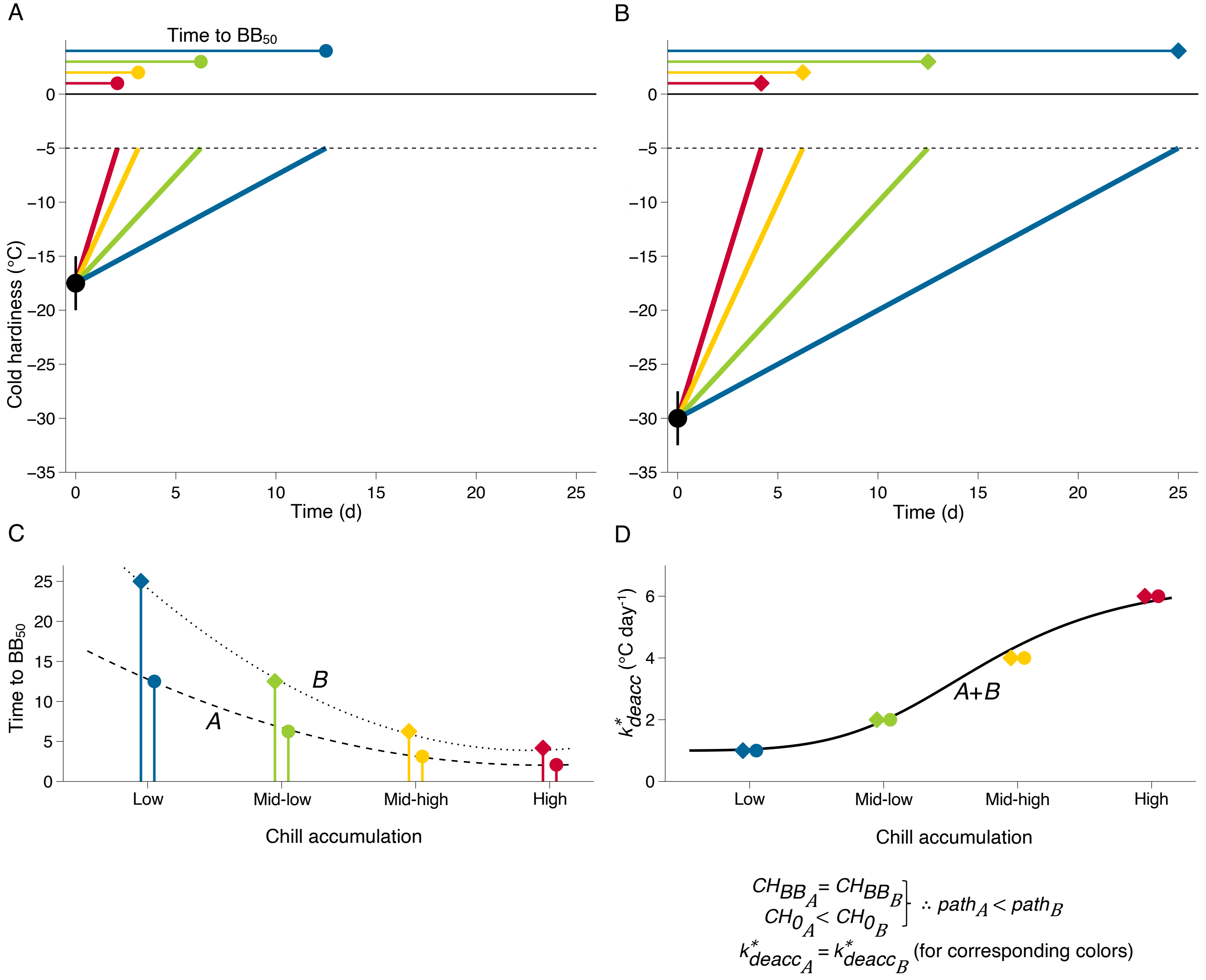


**Supporting Information Figure S4. Different initial cold hardiness, at different levels of chill accumulation.** As *A* has a lesser initial cold hardiness than *B* ($CH_{0_A} < CH_{0_B}$), while having the same cold hardiness at budbreak ($CH_{BB_A} = CH_{BB_B}$), the path length in *A* is shorter than in *B* ($path_A < path_B$). The time to 50% budbreak (*C*) therefore differs for same levels of chill accumulation (corresponding colors). However, the effective rates of deacclimation ($k^*_{deacc}$; *D*) are the same at the same levels of chill accumulation.

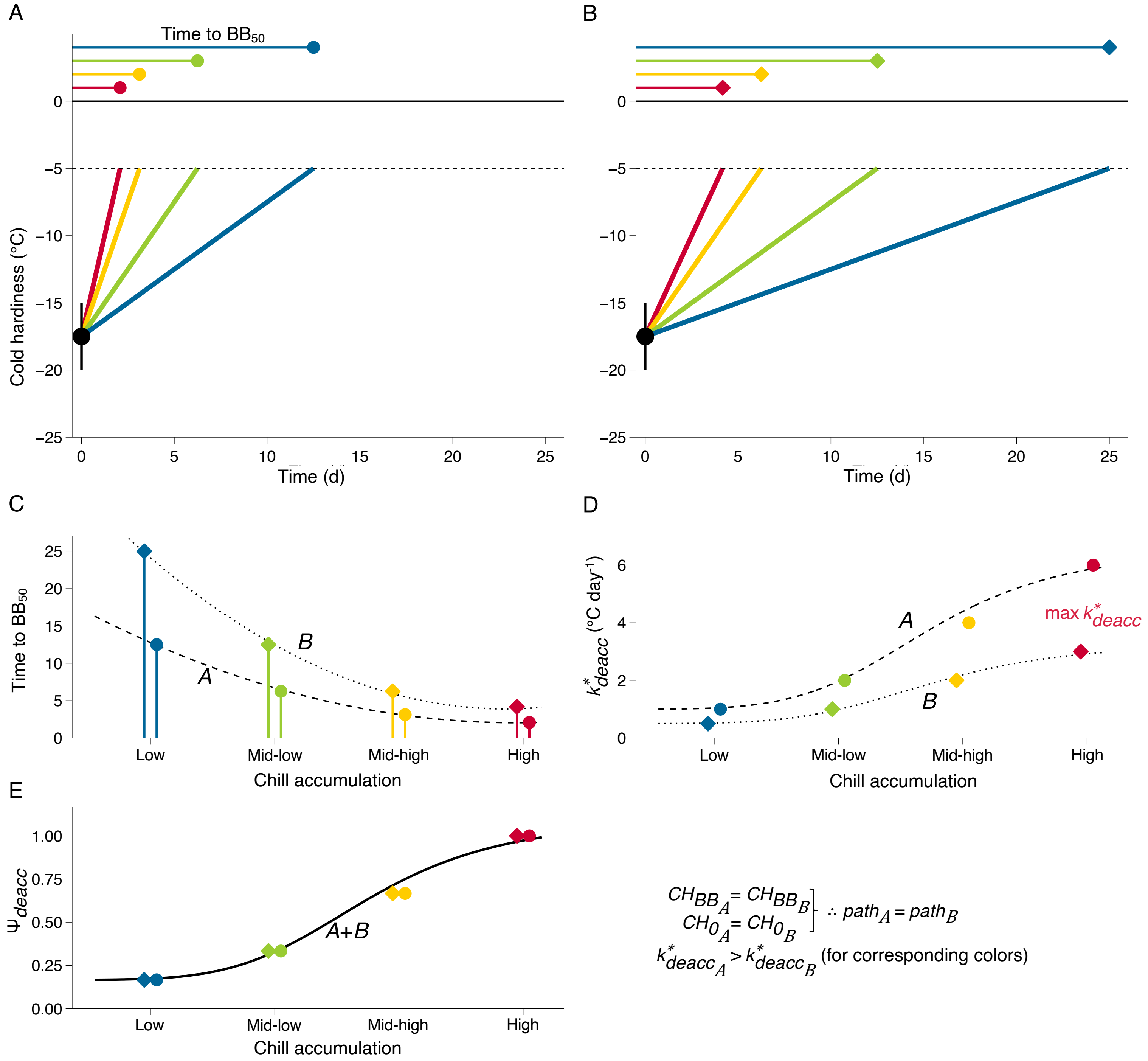


**Supporting Information Figure S5. Different rates of deacclimation, at different levels of chill accumulation.** *A* and *B* have the same initial cold hardiness and same cold hardiness at budbreak, and therefore the same path length ($path_A = path_B$). The time to 50% budbreak (*C*) differs for same levels of chill accumulation (corresponding colors). This occurs as the effective rates of deacclimation ($k^*_{deacc}$; *D*) are greater in *A* than in *B* at the same levels of chill accumulation. However, if the rates are standardized based on the maximum rate observed (max $k^*_{deacc}$), thus generating the deacclimation potential ($\Psi_{deacc}$; *E*), no difference is observed between *A* and *B*.

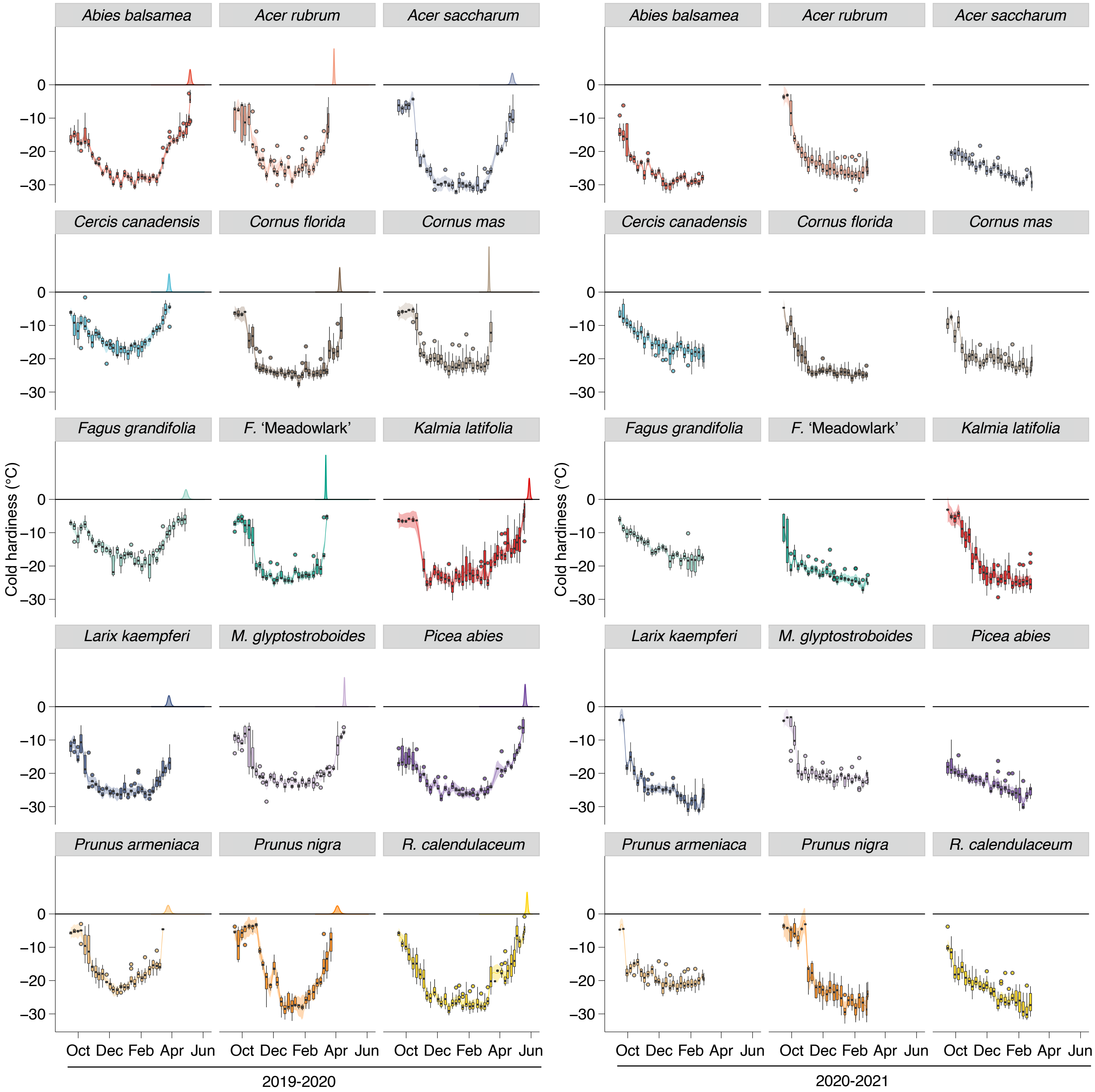


**Supporting Information Figure S6. Field cold hardiness.** Cold hardiness of buds was measured weekly throughout two dormant seasons for 15 species. Density curves above 0 °C line in 2019-2020 represent observed budbreak in the field. Main figure equivalent is **Fig. 2**.

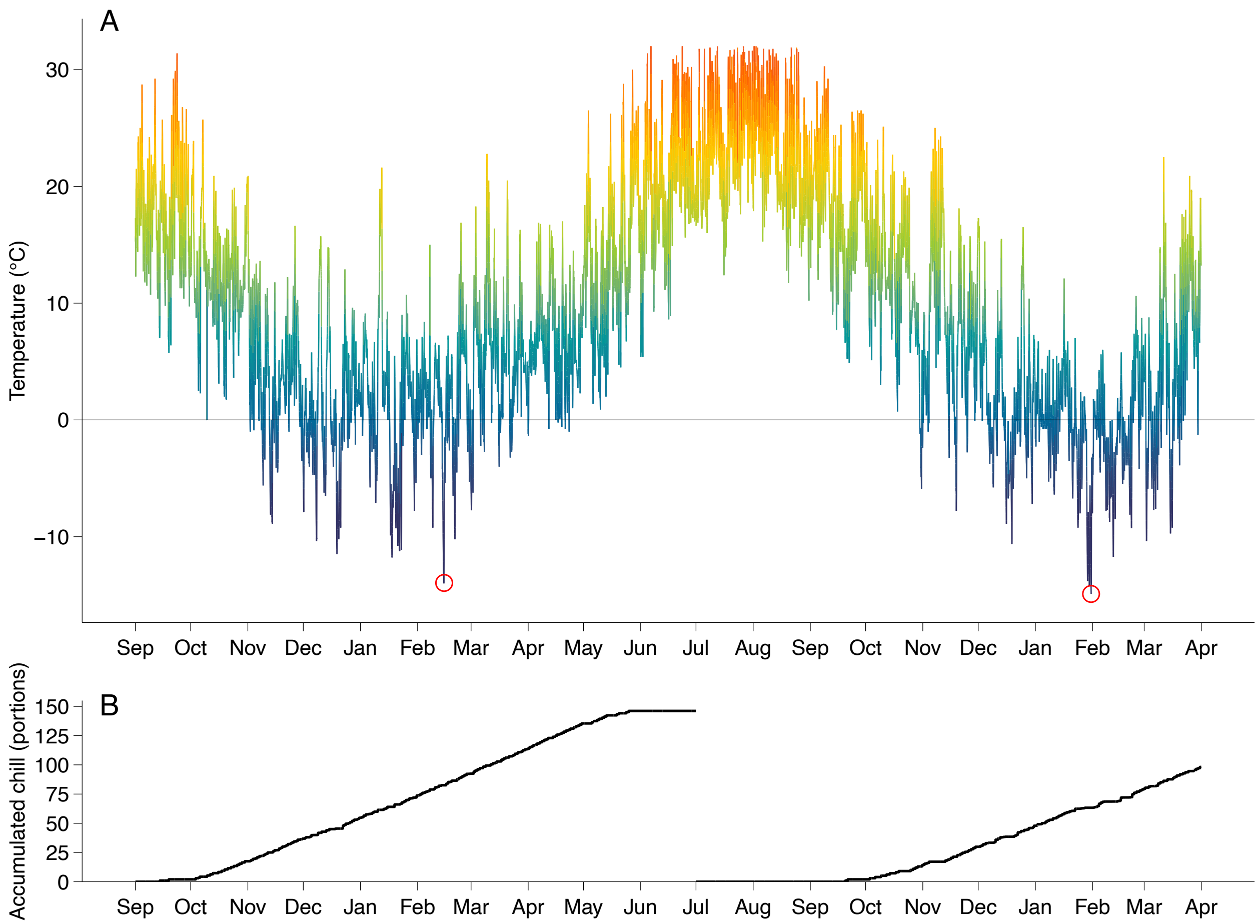


**Supporting Information Figure S7. Temperature conditions in the field.** Observed hourly temperatures (*A*) and chill accumulation (*B*) between 1 September 2019 and 31 March 2021 at the Arnold Arboretum of Harvard University, in Boston, MA, USA. Red circles (*A*) show minimum temperature observed in each season (–14 °C in 2020, –15.7 °C in 2021).

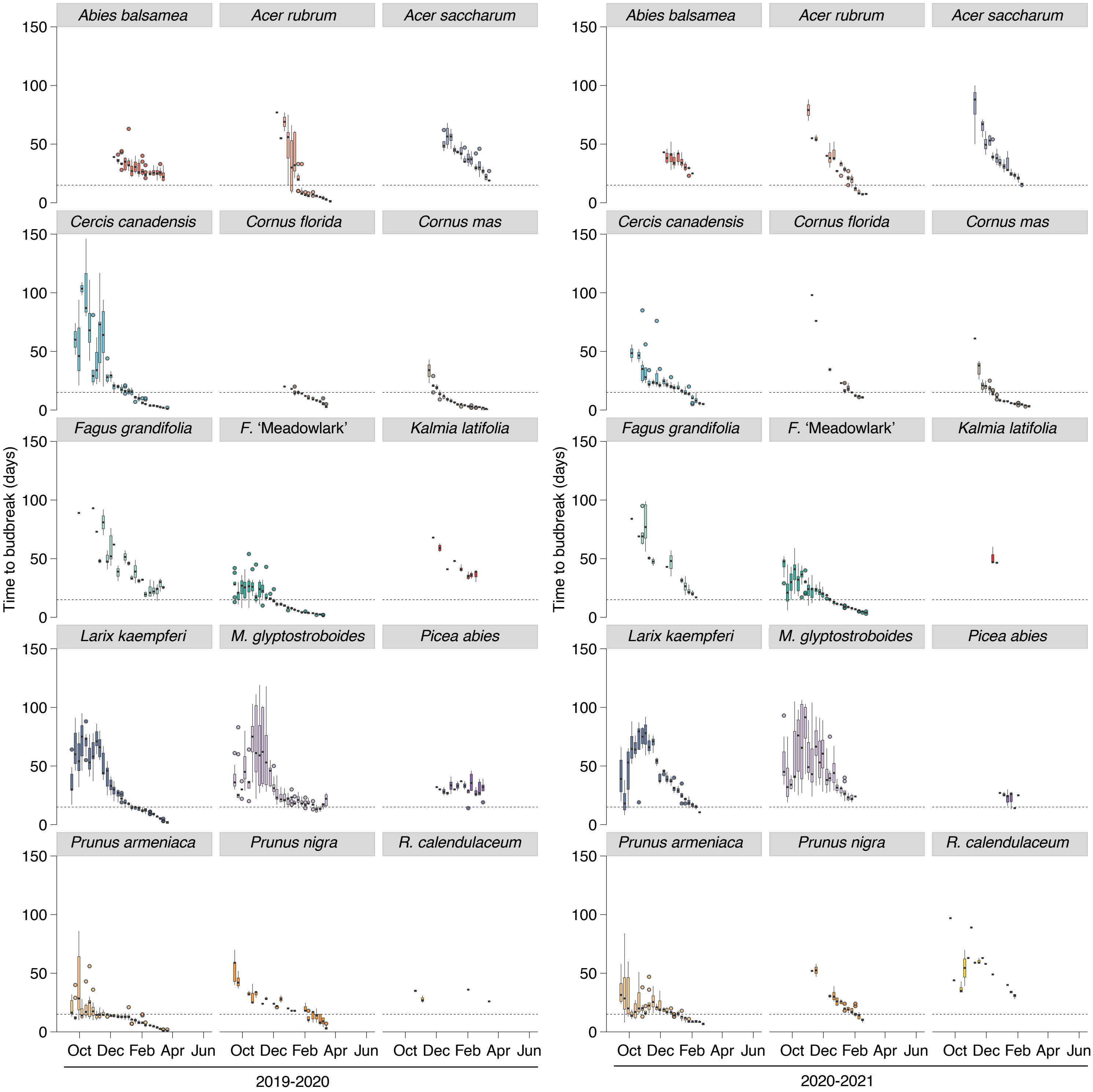


**Supporting Information Figure S8. Budbreak in controlled environment.** Time to budbreak in forcing conditions in controlled environment of buds collected weekly throughout two dormant seasons for 15 species. Main figure equivalent is **Fig. 2**.

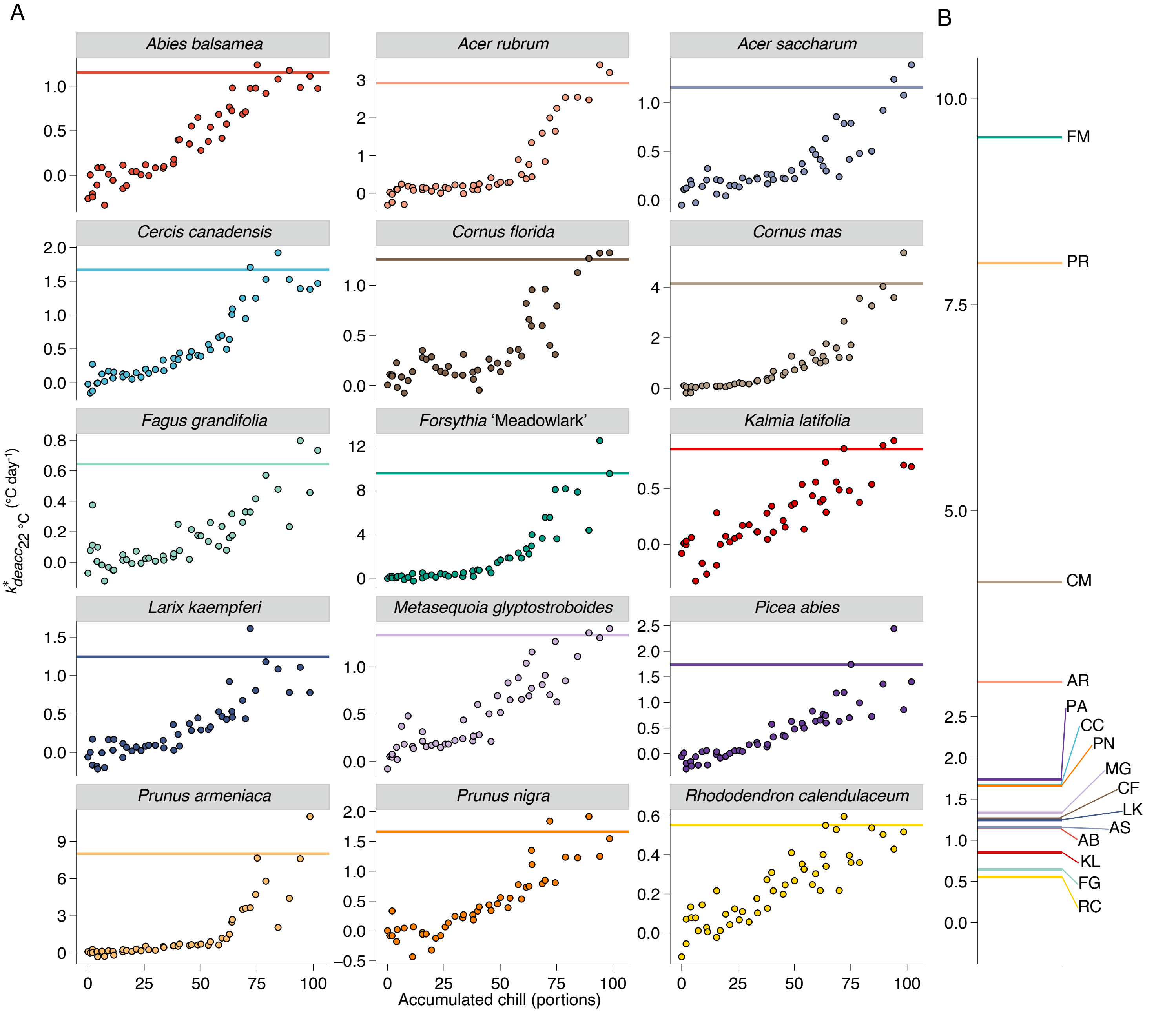


**Supporting Information Figure S9. Deacclimation rates response to chill accumulation.** Deacclimation rates obtained from weekly deacclimation assays at 22 °C for each species during two dormant seasons (*A*). Horizontal line shows maximum rate of deacclimation at 22 °C as the average of the four highest deacclimation rates in each panel. Note different y axes for each species. In the right side *(B)*, maximum rates of all species are plotted together for magnitude comparisons. AB – *Abies balsamea*; AR – *Acer rubrum*; AS – *Acer saccharum*; CC – *Cercis canadensis*; CF – *Cornus florida*; CM – *Cornus mas*; FG – *Fagus grandifolia*; FM – *Forsythia* 'Meadowlark'; KL – *Kalmia latifolia*; LK – *Larix kaempferi*; MG – *Metasequoia glyptostroboides*; PA – *Picea abies*; PR – *Prunus armeniaca*; PN – *Prunus nigra*; RC – *Rhododendron calendulaceum.* Main figure equivalent is **Fig. 4A**.

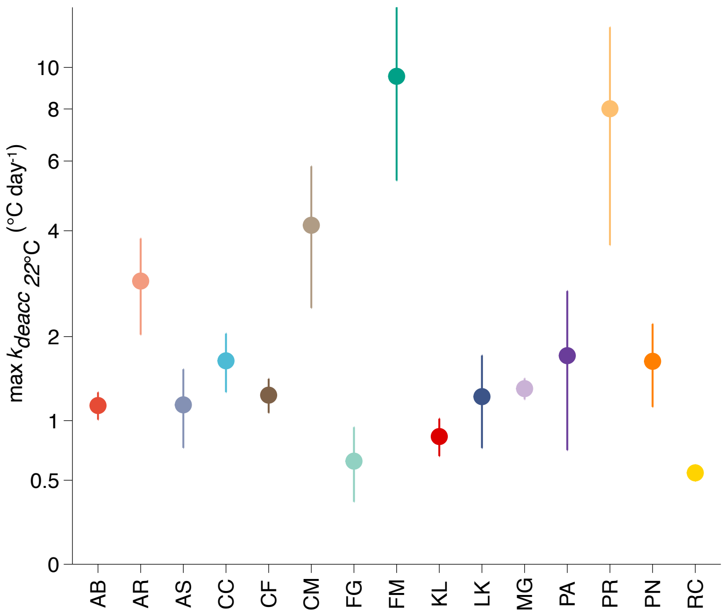


**Supporting Information Figure S10. Maximum deacclimation rates for each species.** Maximum deacclimation rates measured from weekly deacclimation assays at 22 °C for each species during two dormant seasons (log scale). Error bars indicate standard error. AB – *Abies balsamea*; AR – *Acer rubrum*; AS – *Acer saccharum*; CC – *Cercis canadensis*; CF – *Cornus florida*; CM – *Cornus mas*; FG – *Fagus grandifolia*; FM – *Forsythia* 'Meadowlark'; KL – *Kalmia latifolia*; LK – *Larix kaempferi*; MG – *Metasequoia glyptostroboides*; PA – *Picea abies*; PR – *Prunus armeniaca*; PN – *Prunus nigra*; RC – *Rhododendron calendulaceum.*

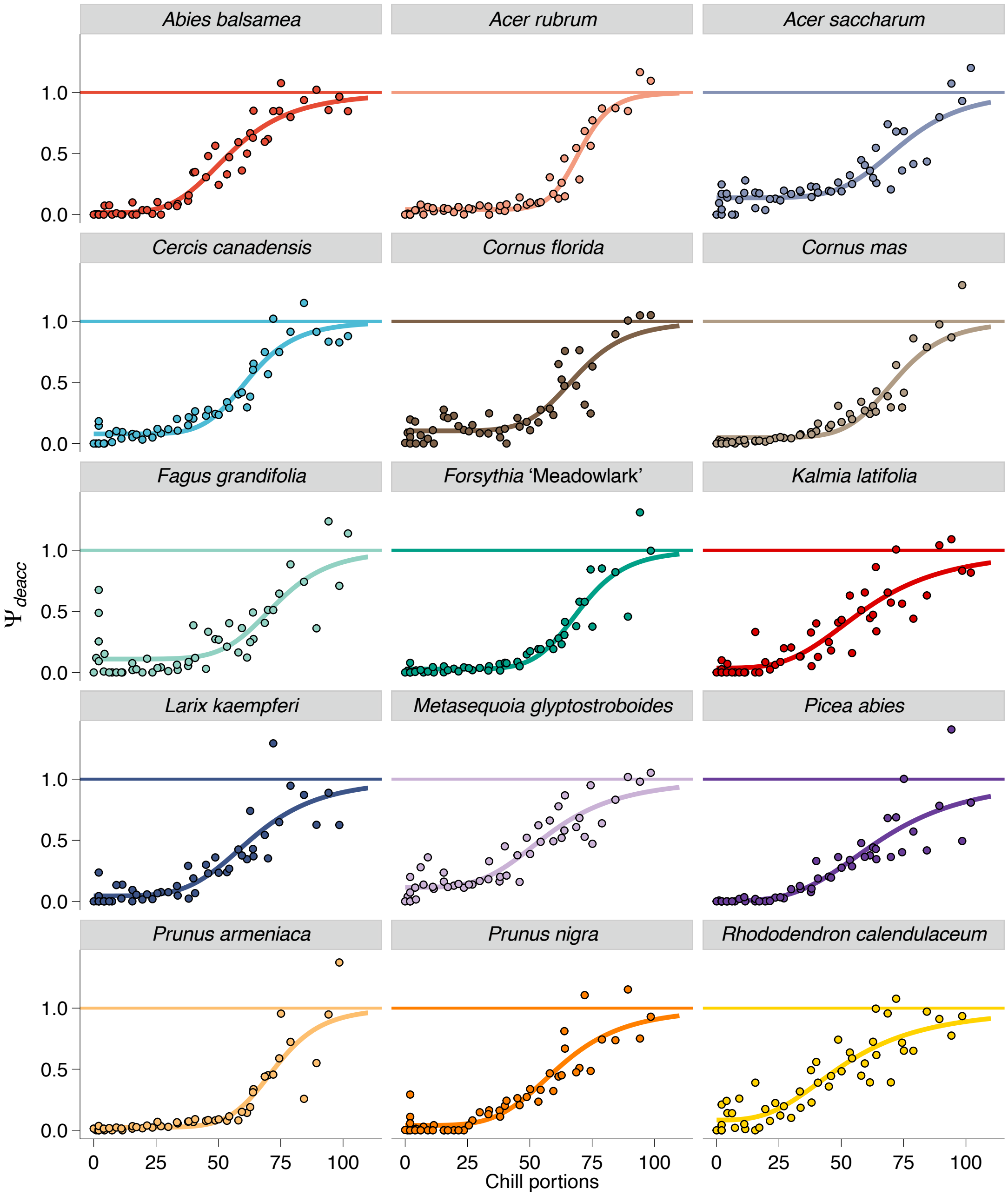


**Supporting Information Figure S11. Normalized deacclimation rates response to chill accumulation.** Deacclimation rates standardized to maximum deacclimation rate at 22 °C for each species produce values of deacclimation potential varying between 0 and 1. Main figure equivalent is **Fig. 4B**.

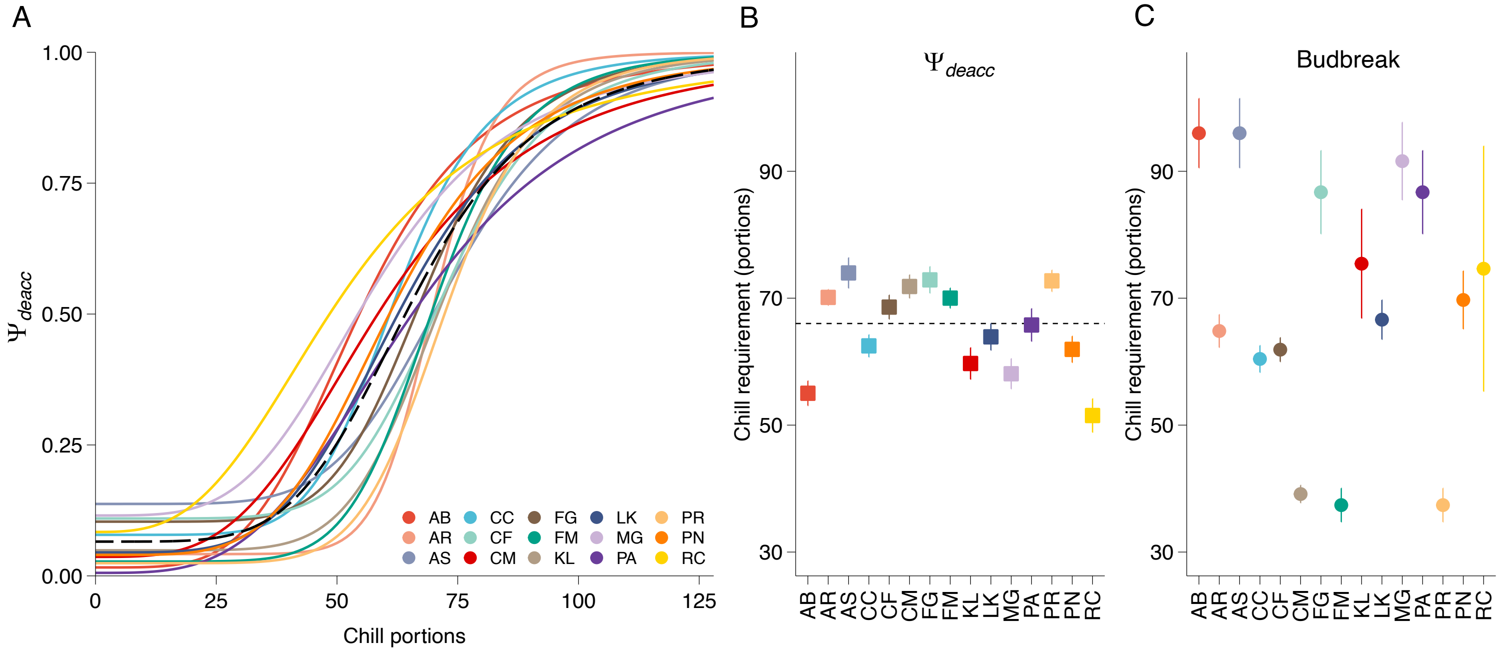


**Supporting Information Figure S12. Dormancy progression based on deacclimation rates and time to budbreak.** Normalized rates of deacclimation produce the deacclimation potential for all species (*A*; $51 \geq n \geq 49$). Inflection points of the deacclimation potential (*B*) are much more similar across species than chill requirement based on budbreak (*C, n* = 4). Dashed lines (*A* and *B*) show the average based on deacclimation potential of all 15 species studied. Error bars, when visible, represent standard deviation. AB – *Abies balsamea*; AR – *Acer rubrum*; AS – *Acer saccharum*; CC – *Cercis canadensis*; CF – *Cornus florida*; CM – *Cornus mas*; FG – *Fagus grandifolia*; FM – *Forsythia* 'Meadowlark'; KL – *Kalmia latifolia*; LK – *Larix kaempferi*; MG – *Metasequoia glyptostroboides*; PA – *Picea abies*; PR – *Prunus armeniaca*; PN – *Prunus nigra*; RC – *Rhododendron calendulaceum.* Main figure equivalent is **Fig. 5.**

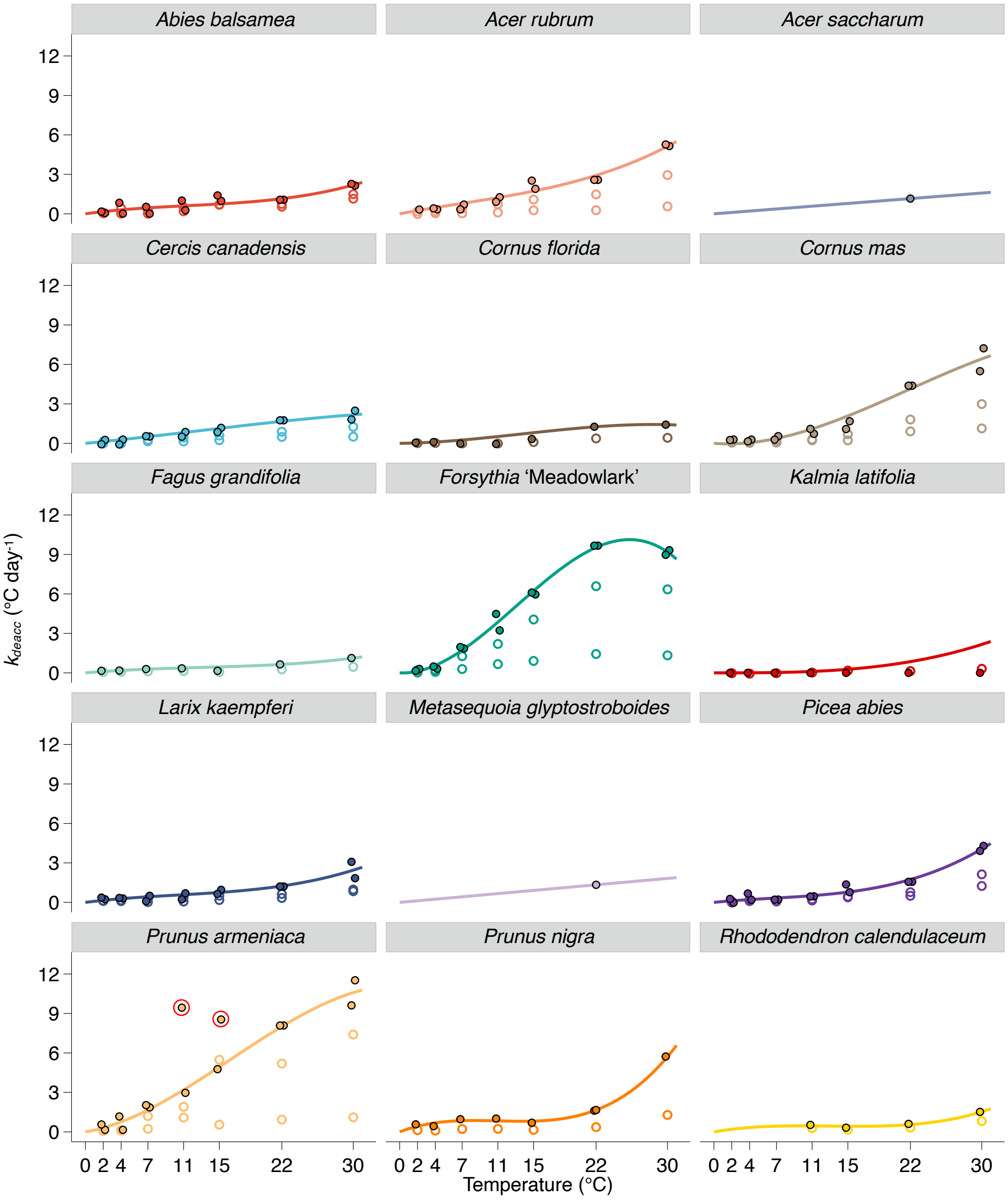


**Supporting Information Fig S13. Deacclimation response to temperature.** Temperature responses for individual species [open circles show measured rates, closed circles shows corrected rates based on $\max k_{deacc_{22\,°C}}$]. Line shows predicted model for $k_{deacc}$ using a third-degree polynomial with intercept = 0. For species which did not have deacclimation measured at temperatures other than 22 °C *(A. saccharum* and *M. glyptostroboides*), temperature response was modeled as a linear regression with intercept = 0 using $\max k_{deacc_{22\,°C}}$. Red circles in *P. armeniaca* show rates removed from model as outliers. Main figure equivalent is **Fig. 6**.

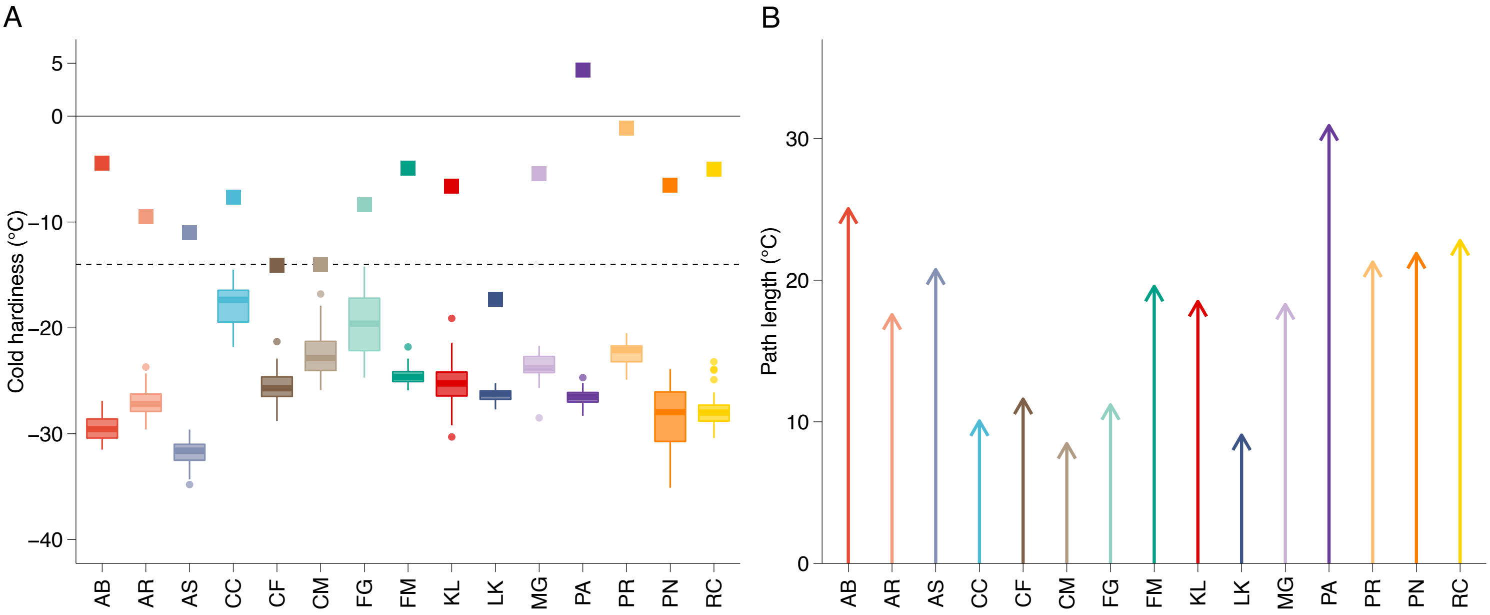


**Supporting Information Figure S14. Cold hardiness parameters for each species.** Maximum cold hardiness ($CH_{max}$) observed in the 2019-2020 season (boxplots) and cold hardiness at budbreak ($CH_{BB}$; squares) for each species (dashed line shows minimum temperature observed in the season for an idea of safety margins; *A*). Path length ( $|CH_{max} – CH_{BB}|$ ) to budbreak for each species (*B* AB – *Abies balsamea*; AR – *Acer rubrum*; AS – *Acer saccharum*; CC – *Cercis canadensis*; CF – *Cornus florida*; CM – *Cornus mas*; FG – *Fagus grandifolia*; FM – *Forsythia* 'Meadowlark'; KL – *Kalmia latifolia*; LK – *Larix kaempferi*; MG – *Metasequoia glyptostroboides*; PA – *Picea abies*; PR – *Prunus armeniaca*; PN – *Prunus nigra*; RC – *Rhododendron calendulaceum.* Main figure equivalent is **Fig. 7**.

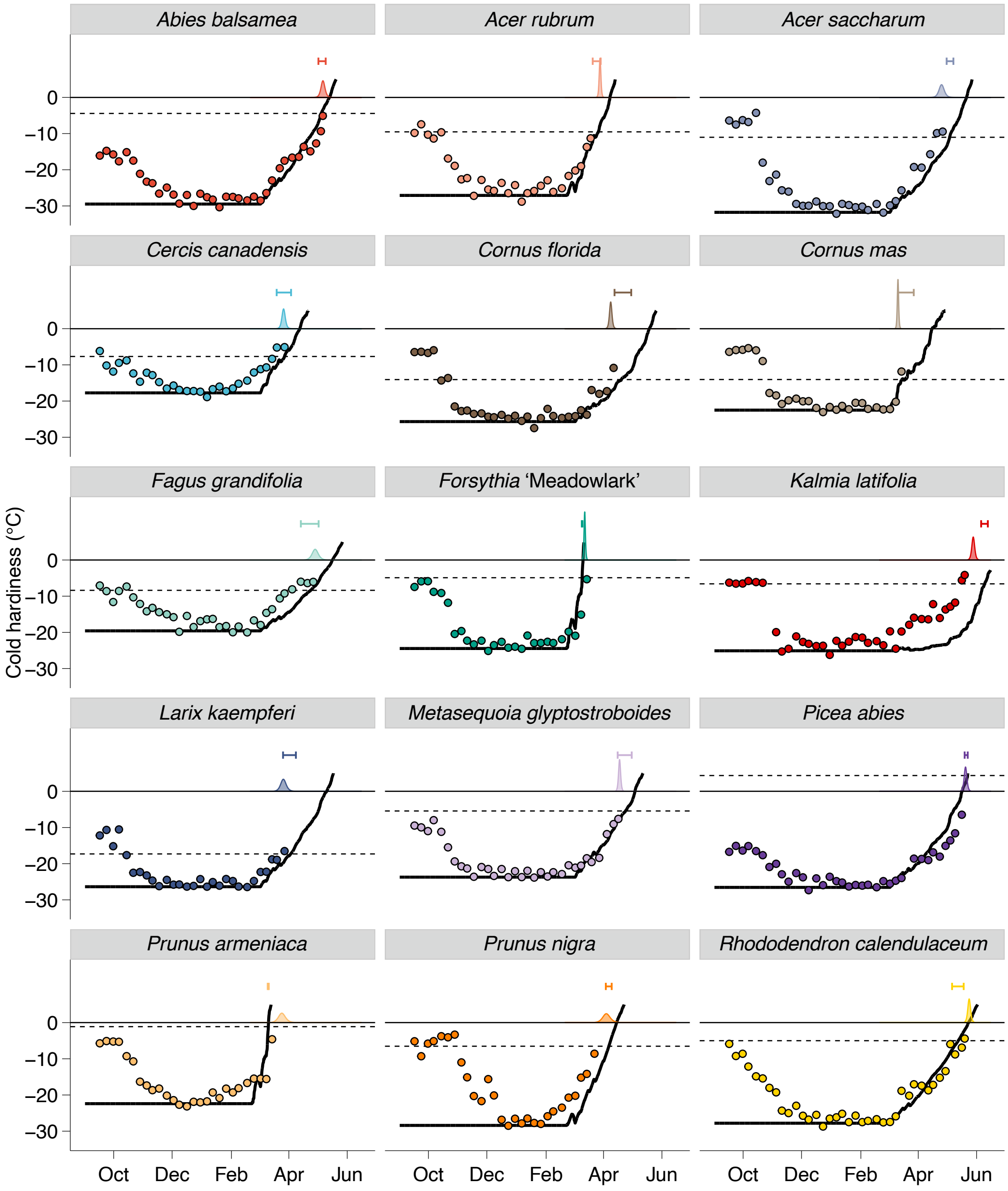


**Supporting Information Figure S15. Predicting budbreak based on cold hardiness dynamics.** Average cold hardiness [circles, no error for simplicity (see **SI Appendix Fig. S6**)] during the entire season are shown, as well as field budbreak (density plots at 0 °C). Predicted cold hardiness started at maximum cold hardiness ($CH_{max}$) and runs until +5 °C based on species-specific deacclimation parameters (solid line). Budbreak is predicted to occur when prediction line crosses the cold hardiness at budbreak ($CH_{BB}$; dashed lines) for each species. The interval is produced by adding error of ± 2.5 °C to $CH_{max}$. Main figure equivalent is **Fig. 8**.

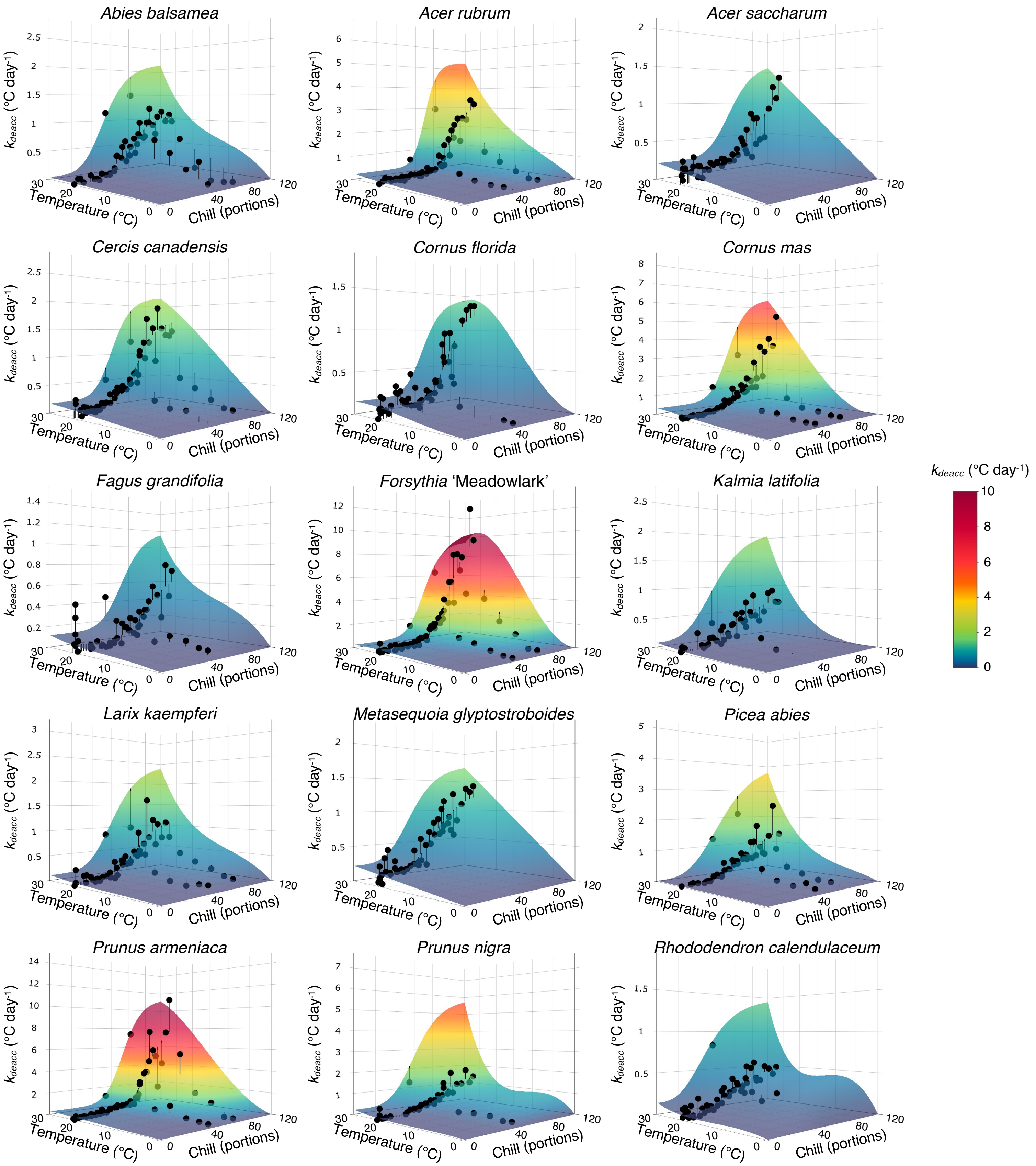


**Supporting Information Figure S16. Plasticity in temperature and chill accumulation response.** Predicted rates of deacclimation (surface plot) based on individual species measured responses (circles) to chill and temperature. Lines connect points to surface plot to demonstrate residuals. Different z axes are used such that surface curvature is visible, but color scale used is the same for all species.

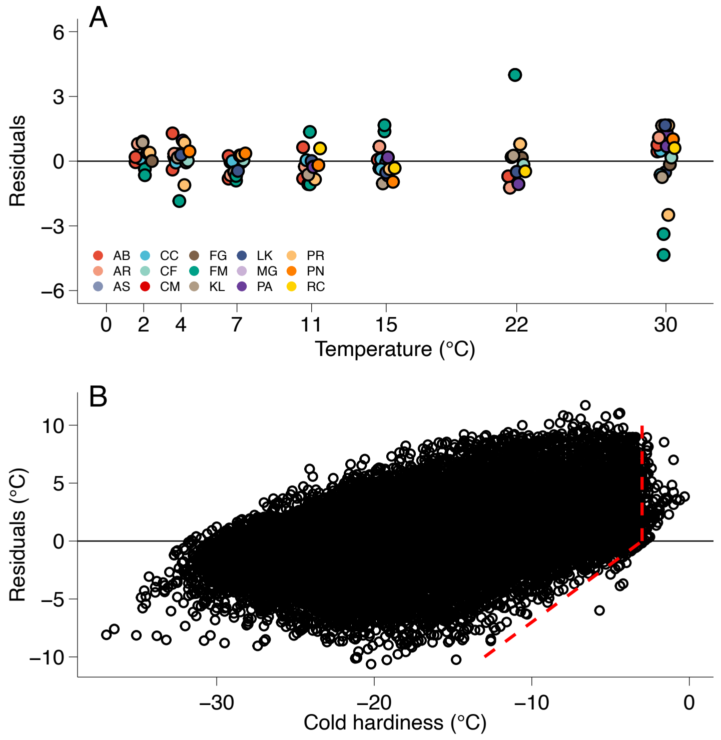


**Supporting Information Figure S17. Residuals for rates of deacclimation and cold hardiness measurements.** Studentized residuals for rates of deacclimation for 9 species based on a linear model response (*A*) and residuals based on linear deacclimation rates at 22 °C at weekly collections for all species (*B*). Red dashed line shows approximate limit of detection at –3 °C. AB – *Abies balsamea*; AR – *Acer rubrum*; AS – *Acer saccharum*; CC – *Cercis canadensis*; CF – *Cornus florida*; CM – *Cornus mas*; FG – *Fagus grandifolia*; FM – *Forsythia* 'Meadowlark'; KL – *Kalmia latifolia*; LK – *Larix kaempferi*; MG – *Metasequoia glyptostroboides*; PA – *Picea abies*; PR – *Prunus armeniaca*; PN – *Prunus nigra*; RC – *Rhododendron calendulaceum.*

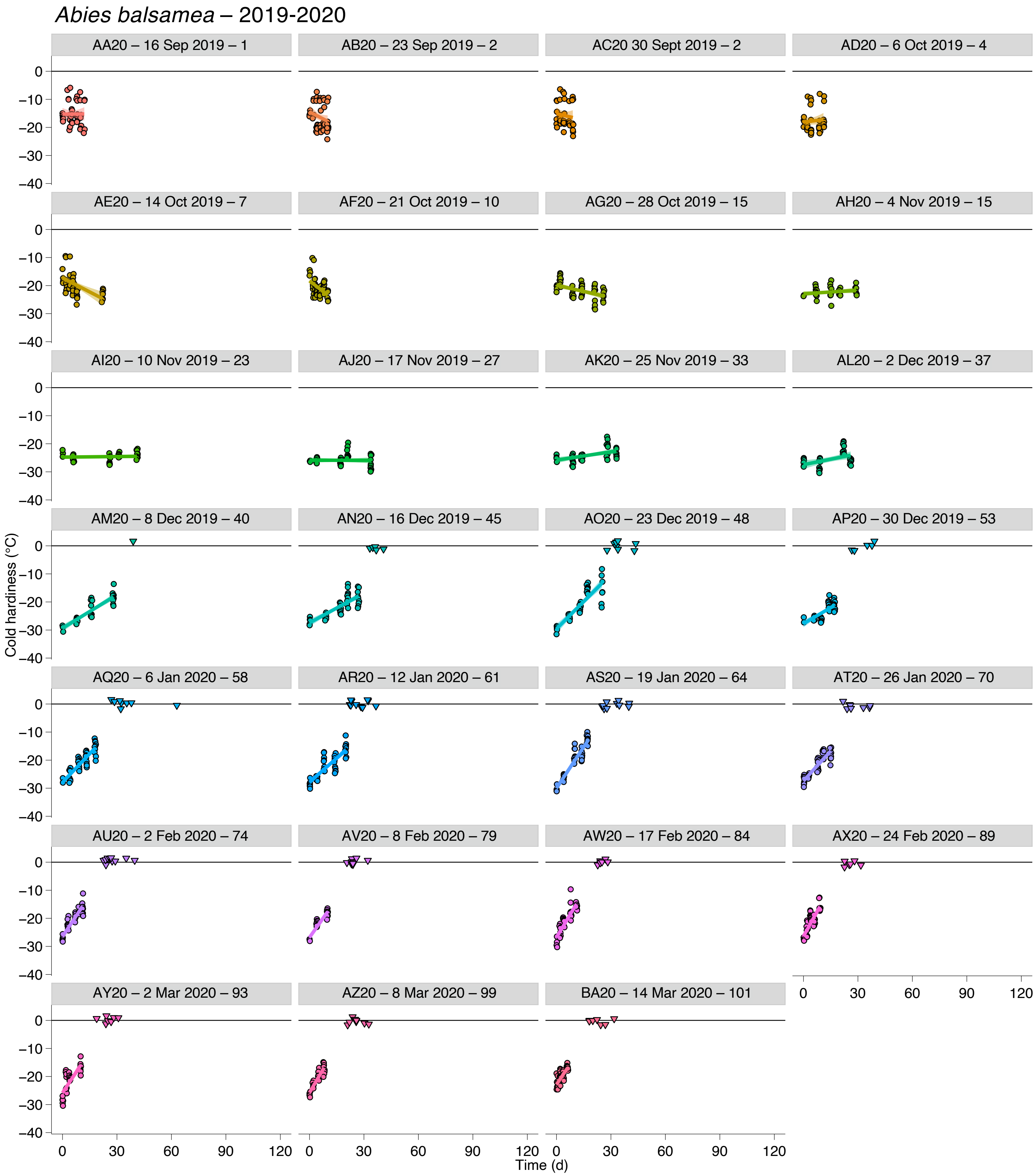


**Supporting Information Figure S18. Deacclimation and budbreak of *Abies balsamea* buds collected during the 2019-2020 dormant season.** Loss of cold hardiness (circles) and budbreak (upside down triangles at 0 °C line) for *Abies balsamea*. Each graph is identified by collection name, date of collection, and chill accumulated in portions.

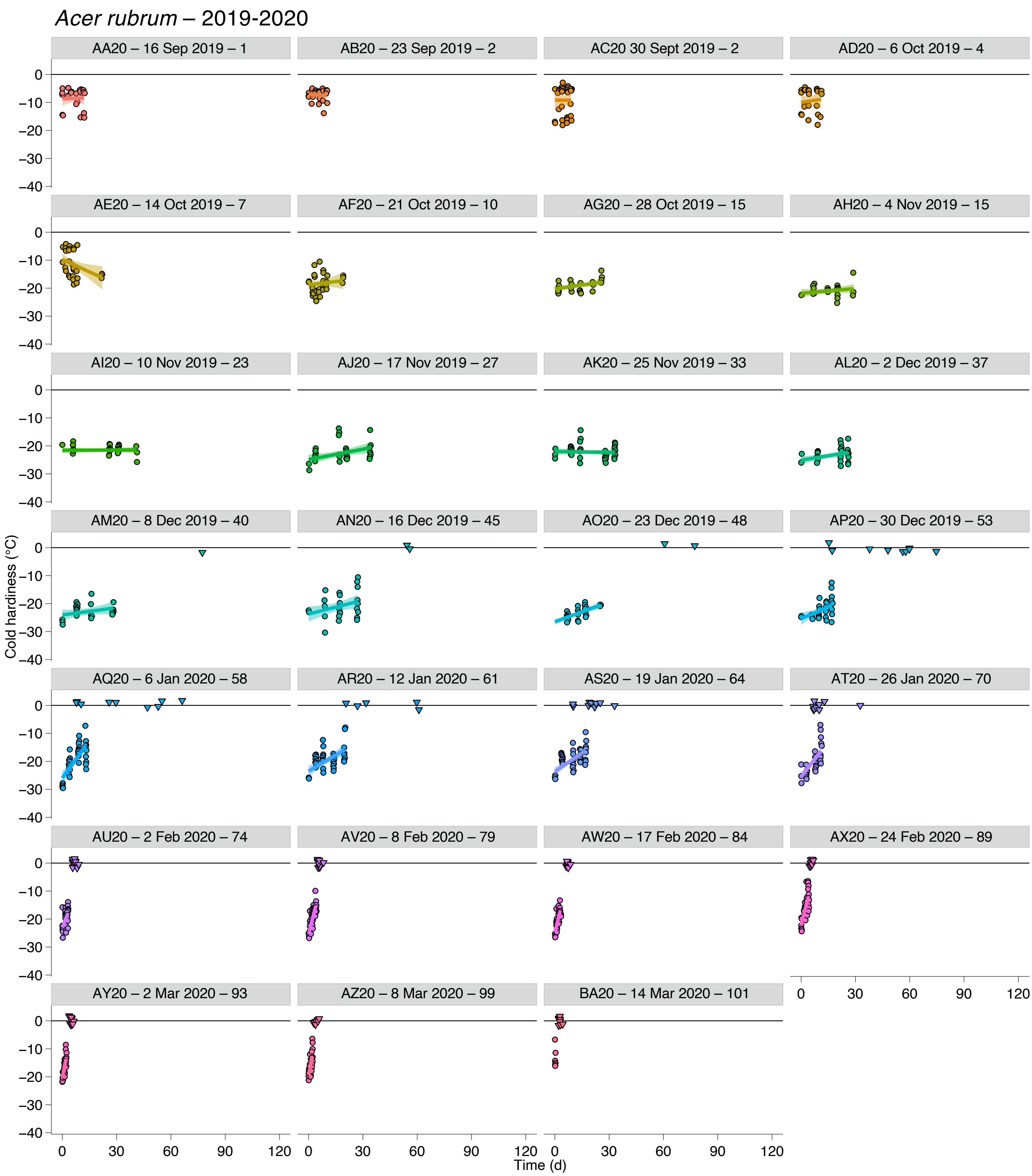


**Supporting Information Figure S19. Deacclimation and budbreak of *Acer rubrum* buds collected during the 2019-2020 dormant season.** Loss of cold hardiness (circles) and budbreak (upside down triangles at 0 °C line) for *Acer rubrum*. Each graph is identified by collection name, date of collection, and chill accumulated in portions.

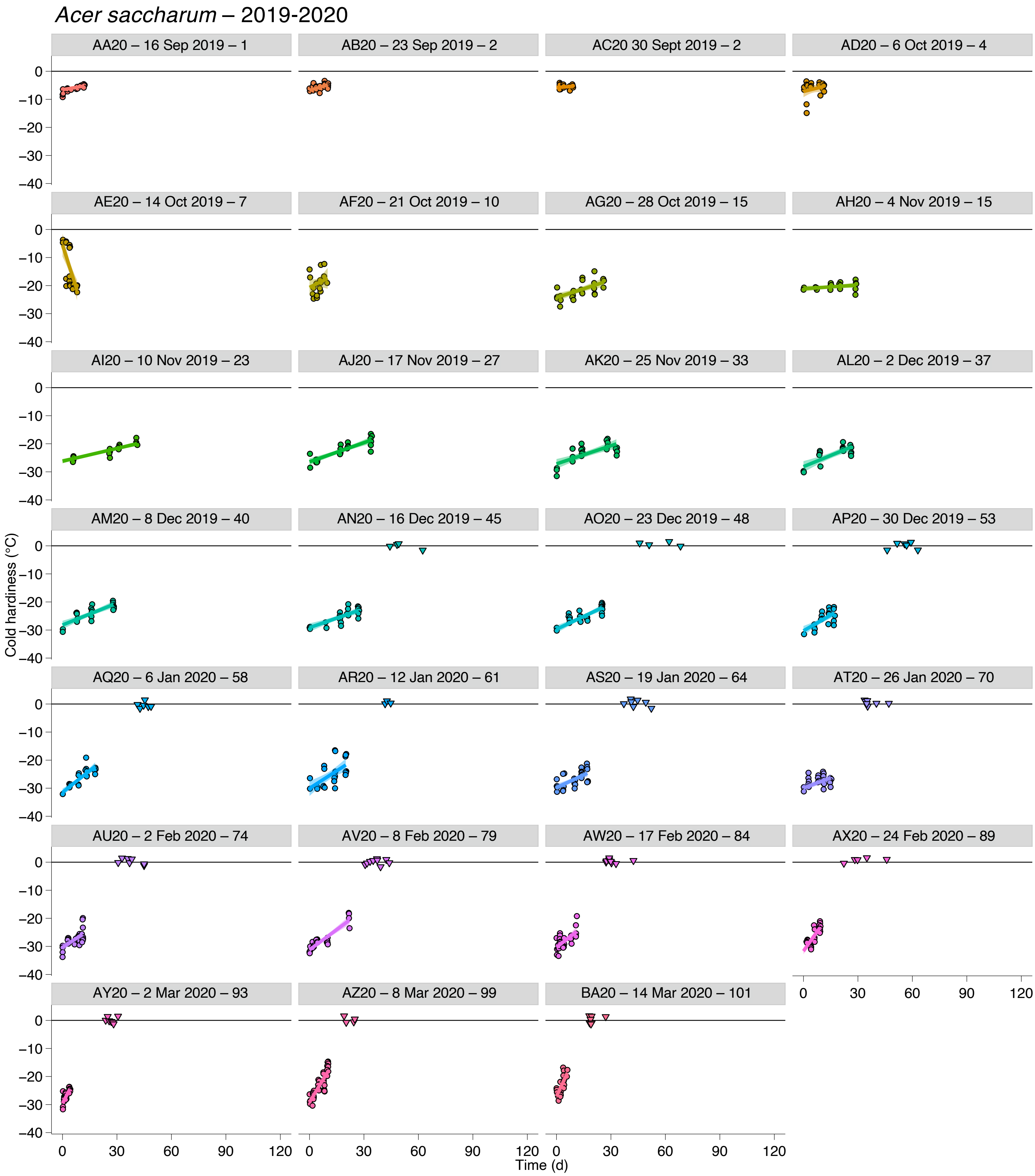


**Supporting Information Figure S20. Deacclimation and budbreak of *Acer saccharum* buds collected during the 2019-2020 dormant season.** Loss of cold hardiness (circles) and budbreak (upside down triangles at 0 °C line) for *Acer saccharum*. Each graph is identified by collection name, date of collection, and chill accumulated in portions.

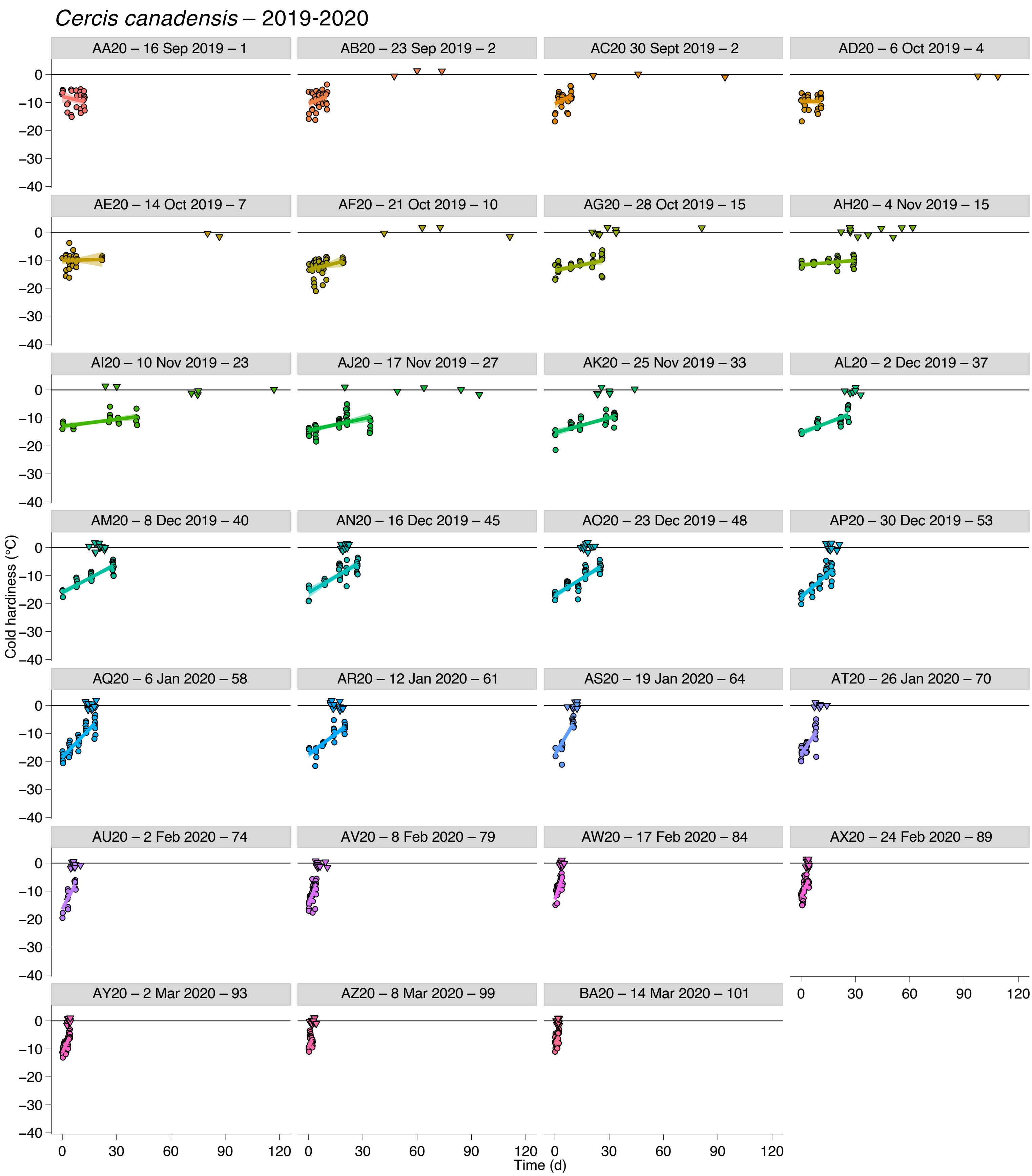


**Supporting Information Figure S21. Deacclimation and budbreak of *Cercis canadensis* buds collected during the 2019-2020 dormant season.** Loss of cold hardiness (circles) and budbreak (upside down triangles at 0 °C line) for *Cercis canadensis*. Each graph is identified by collection name, date of collection, and chill accumulated in portions.

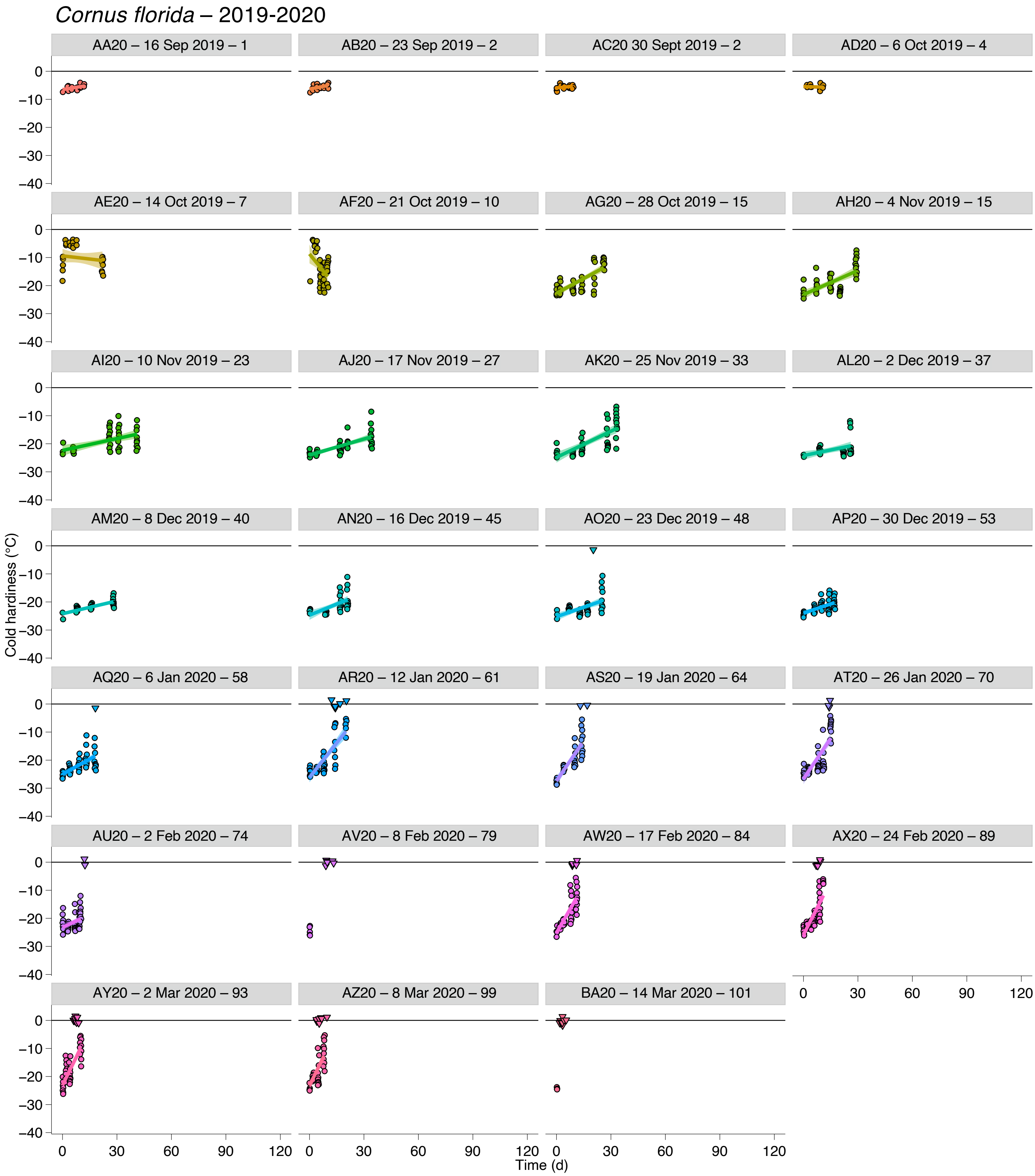


**Supporting Information Figure S22. Deacclimation and budbreak of *Cornus florida* buds collected during the 2019-2020 dormant season.** Loss of cold hardiness (circles) and budbreak (upside down triangles at 0 °C line) for *Cornus florida*. Each graph is identified by collection name, date of collection, and chill accumulated in portions.

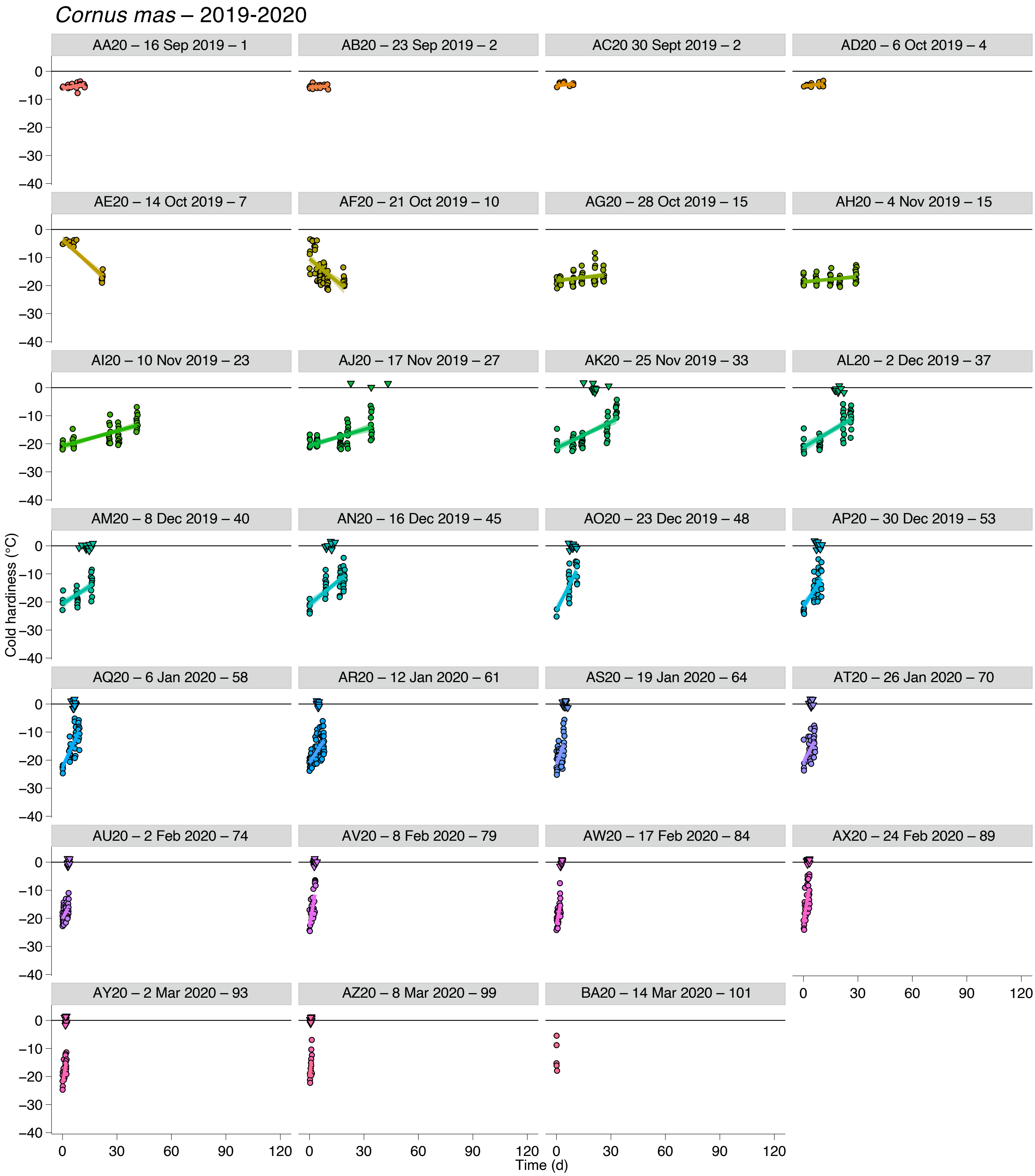


**Supporting Information Figure S23. Deacclimation and budbreak of *Cornus mas* buds collected during the 2019-2020 dormant season.** Loss of cold hardiness (circles) and budbreak (upside down triangles at 0 °C line) for *Cornus mas*. Each graph is identified by collection name, date of collection, and chill accumulated in portions.

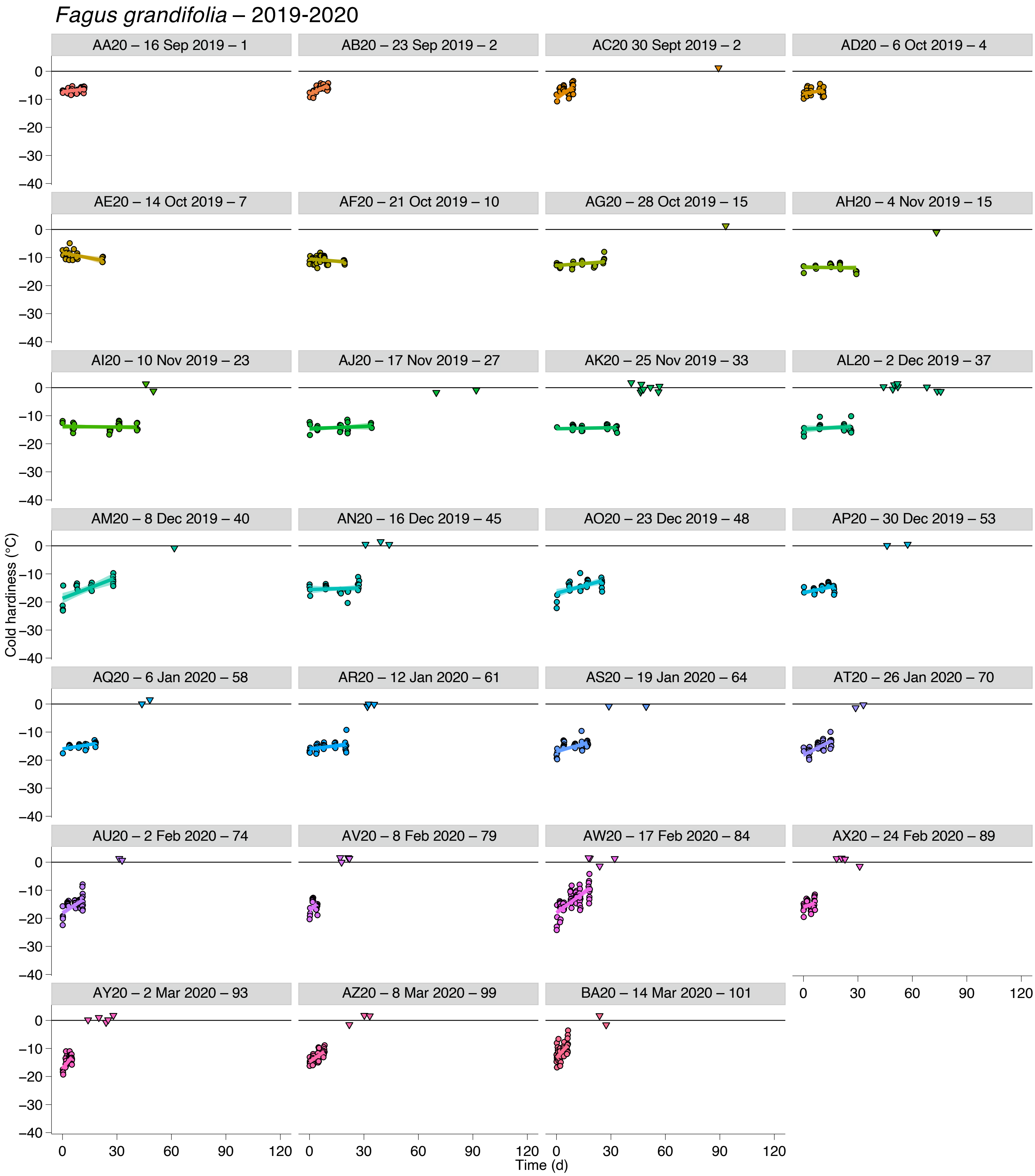


**Supporting Information Figure S24. Deacclimation and budbreak of *Fagus grandifolia* buds collected during the 2019-2020 dormant season.** Loss of cold hardiness (circles) and budbreak (upside down triangles at 0 °C line) for *Fagus grandifolia*. Each graph is identified by collection name, date of collection, and chill accumulated in portions.

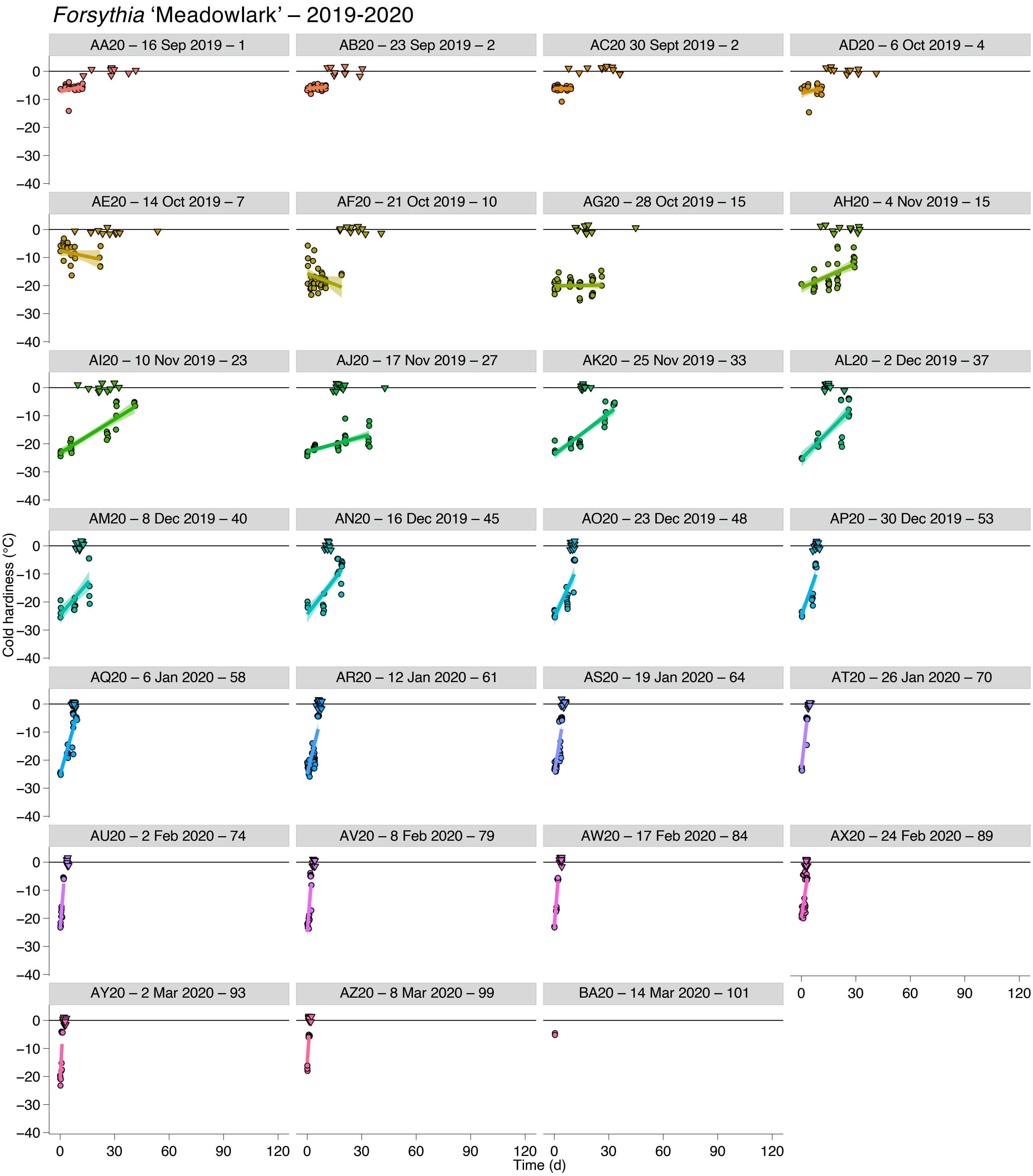


**Supporting Information Figure S25. Deacclimation and budbreak of *Forsythia* 'Meadowlark' buds collected during the 2019-2020 dormant season.** Loss of cold hardiness (circles) and budbreak (upside down triangles at 0 °C line) for *Forsythia* 'Meadowlark'. Each graph is identified by collection name, date of collection, and chill accumulated in portions.

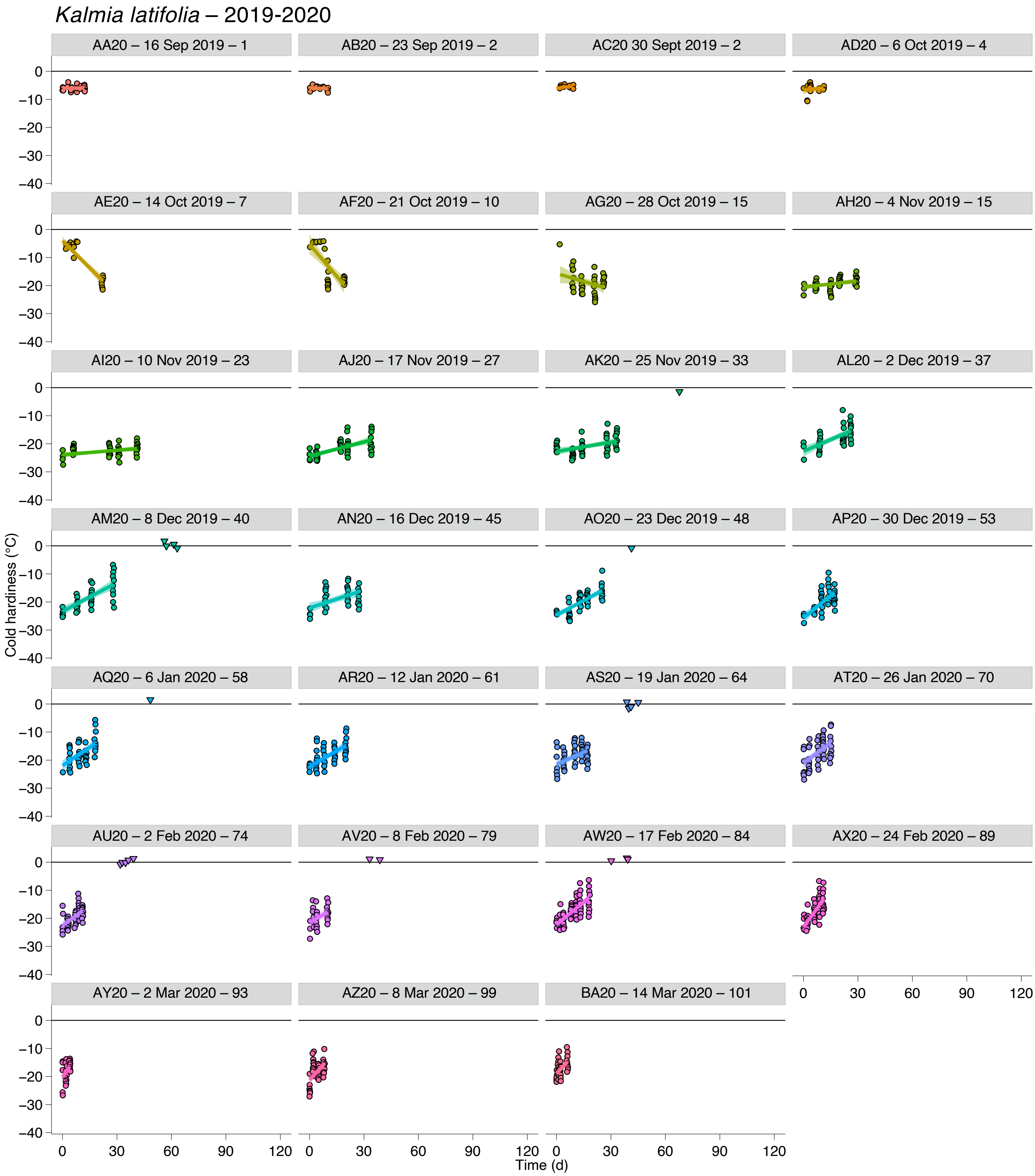


**Supporting Information Figure S26. Deacclimation and budbreak of *Kalmia latifolia* buds collected during the 2019-2020 dormant season.** Loss of cold hardiness (circles) and budbreak (upside down triangles at 0 °C line) for *Kalmia latifolia*. Each graph is identified by collection name, date of collection, and chill accumulated in portions.

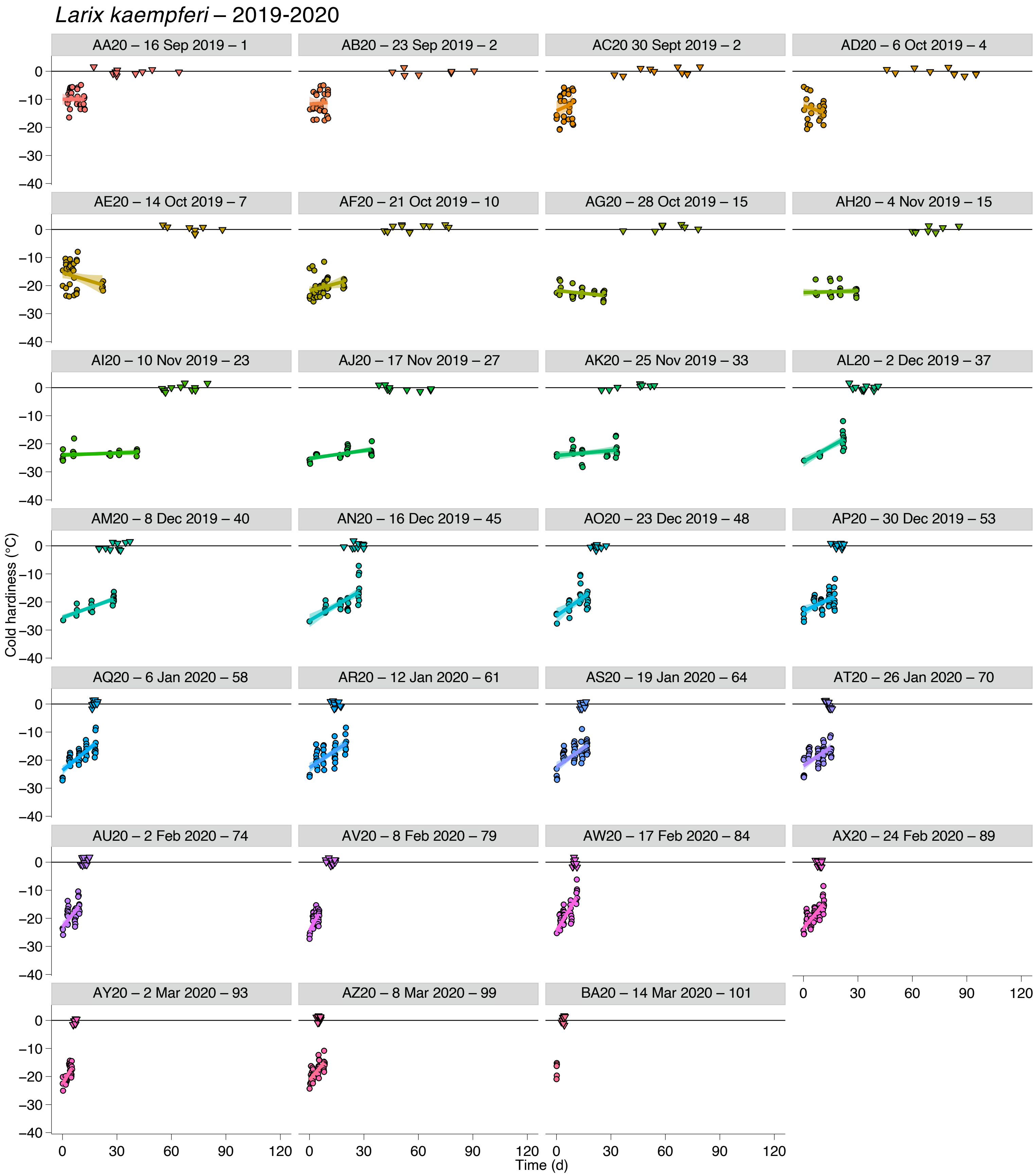


**Supporting Information Figure S27. Deacclimation and budbreak of *Larix kaempferi* buds collected during the 2019-2020 dormant season.** Loss of cold hardiness (circles) and budbreak (upside down triangles at 0 °C line) for *Larix kaempferi*. Each graph is identified by collection name, date of collection, and chill accumulated in portions.

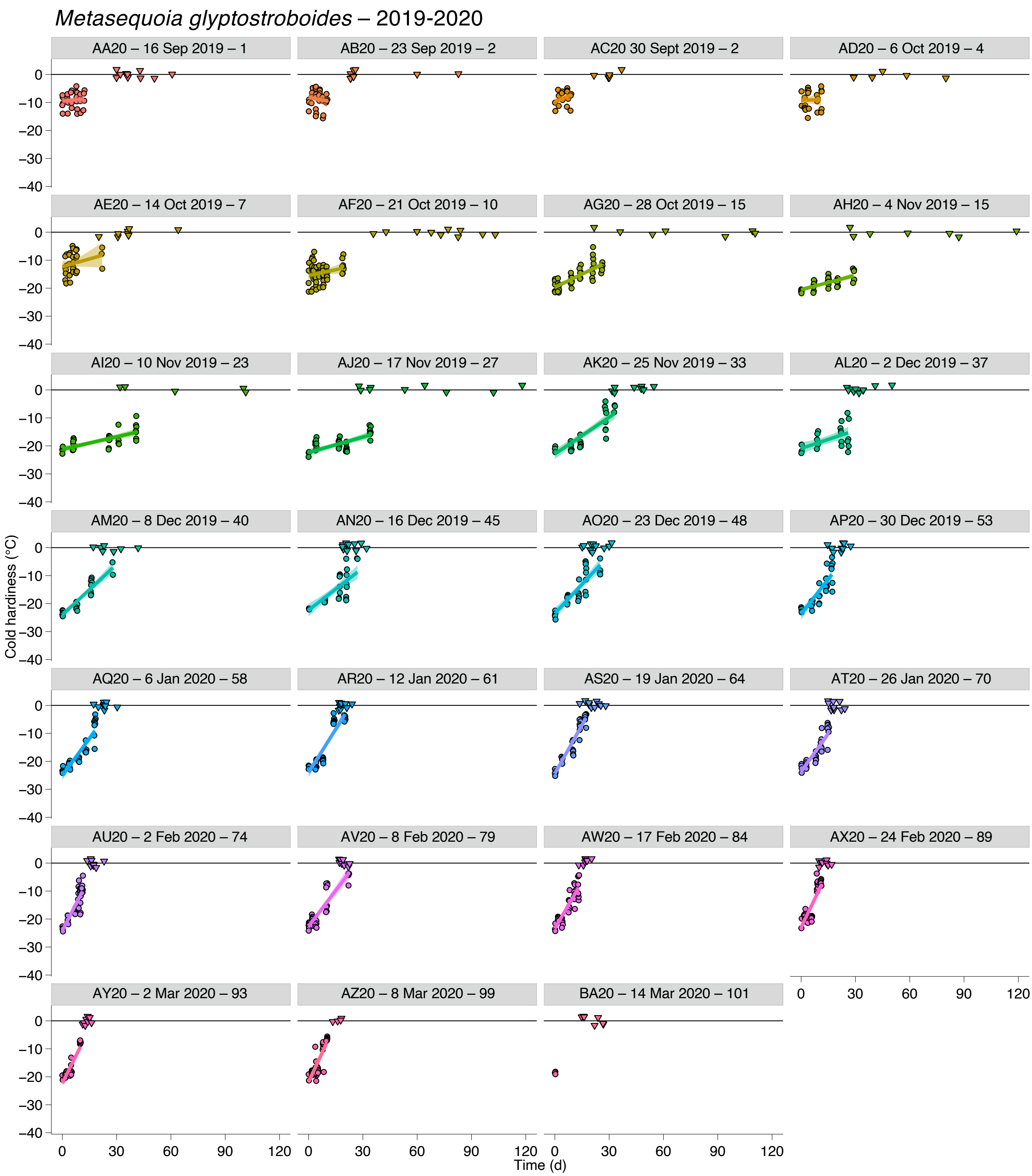


**Supporting Information Figure S28. Deacclimation and budbreak of *Metasequoia glyptostroboides* buds collected during the 2019-2020 dormant season.** Loss of cold hardiness (circles) and budbreak (upside down triangles at 0 °C line) for *Metasequoia glyptostroboides*. Each graph is identified by collection name, date of collection, and chill accumulated in portions.

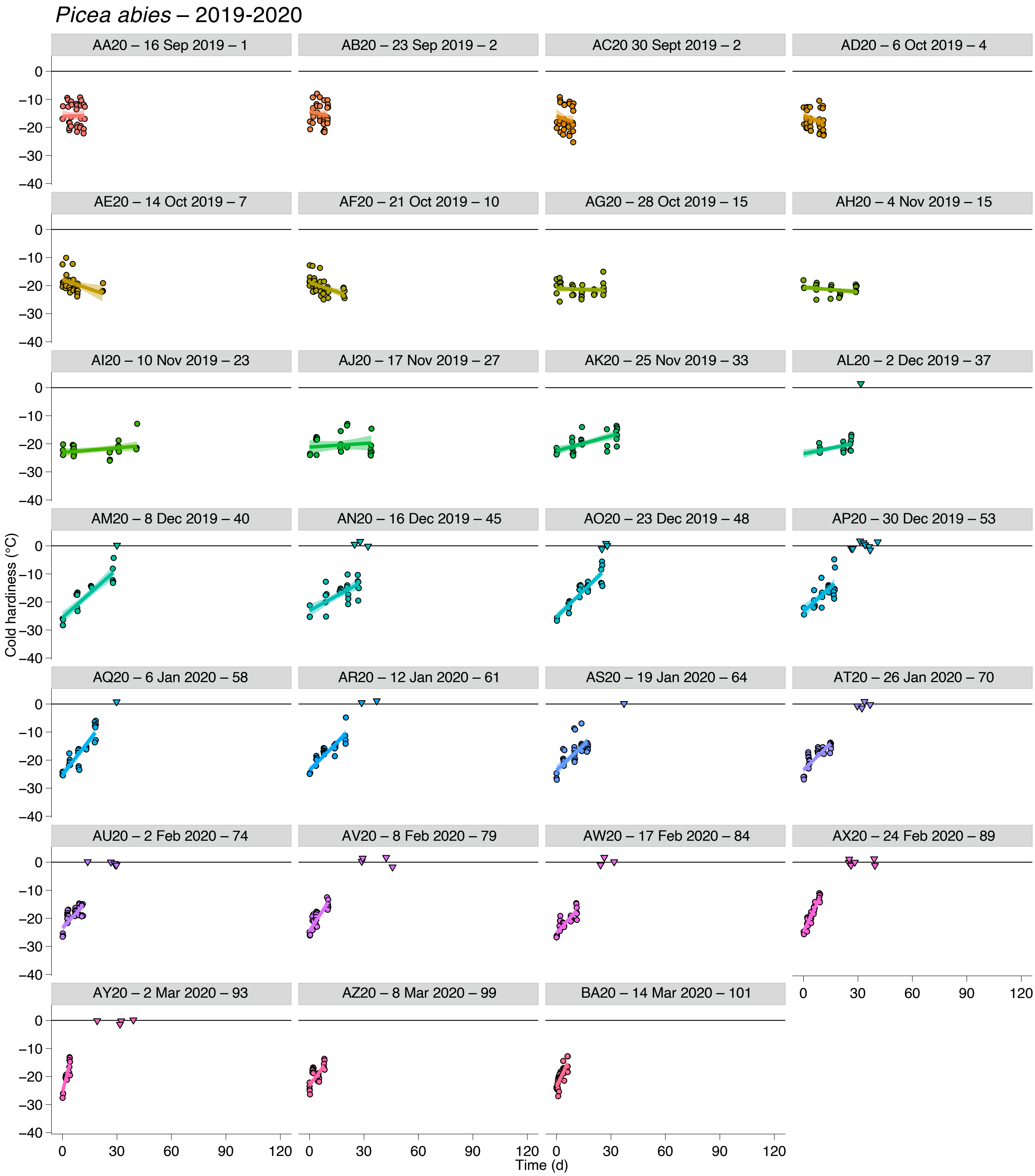


**Supporting Information Figure S29. Deacclimation and budbreak of *Picea abies* buds collected during the 2019-2020 dormant season.** Loss of cold hardiness (circles) and budbreak (upside down triangles at 0 °C line) for *Picea abies*. Each graph is identified by collection name, date of collection, and chill accumulated in portions.

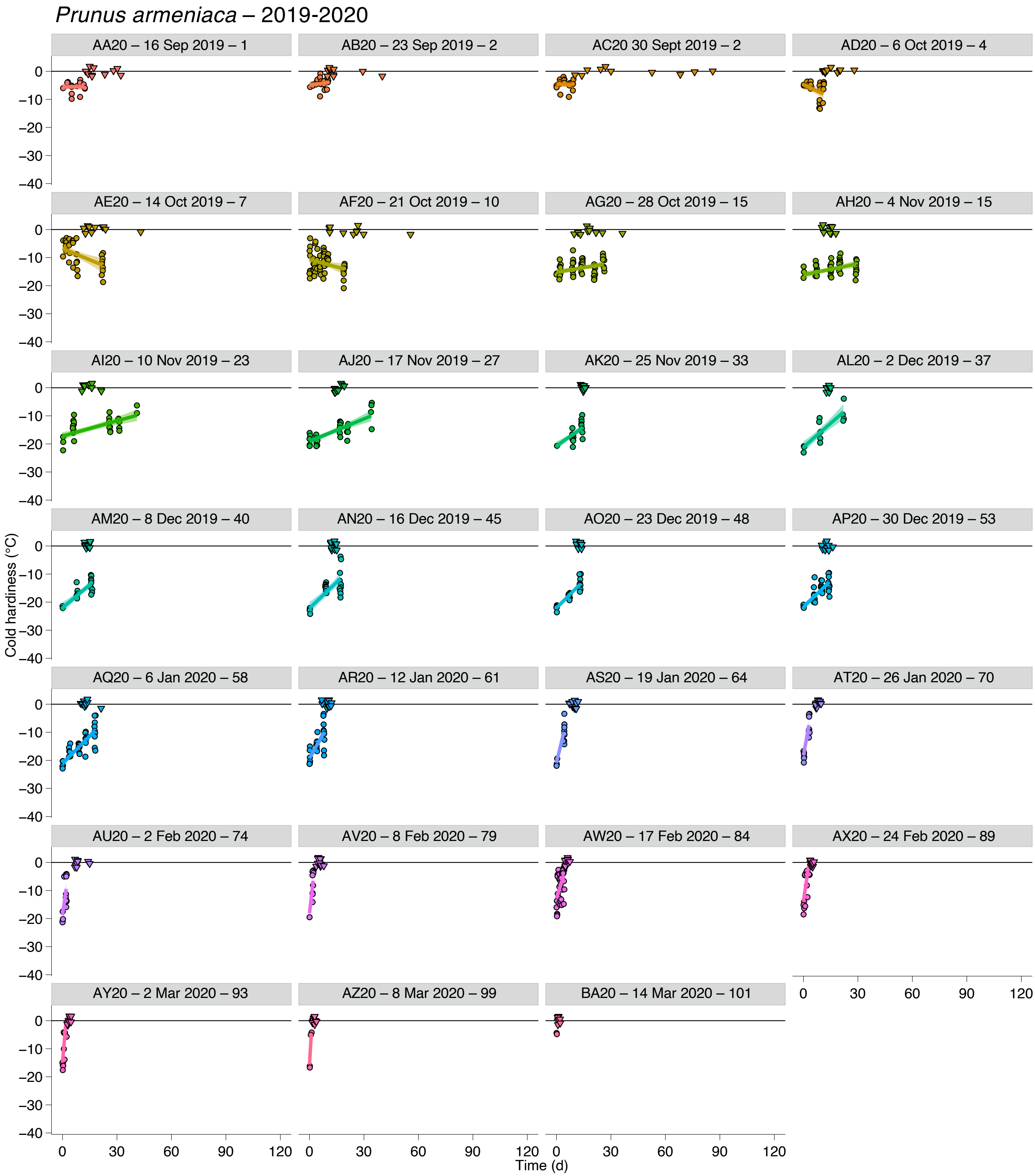


**Supporting Information Figure S30. Deacclimation and budbreak of *Prunus armeniaca* buds collected during the 2019-2020 dormant season.** Loss of cold hardiness (circles) and budbreak (upside down triangles at 0 °C line) for *Prunus armeniaca*. Each graph is identified by collection name, date of collection, and chill accumulated in portions.

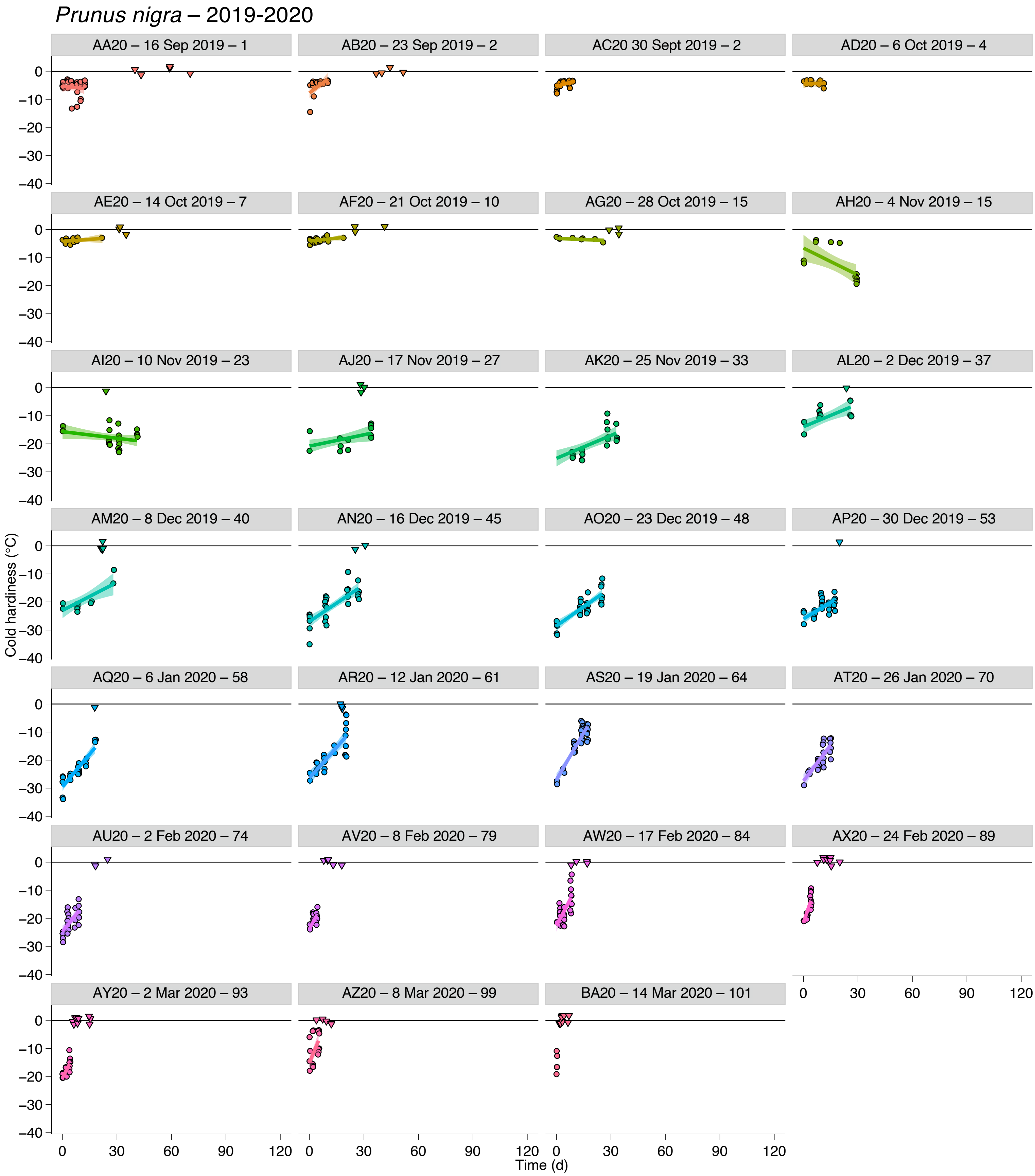


**Supporting Information Figure S31. Deacclimation and budbreak of *Prunus nigra* buds collected during the 2019-2020 dormant season.** Loss of cold hardiness (circles) and budbreak (upside down triangles at 0 °C line) for *Prunus nigra*. Each graph is identified by collection name, date of collection, and chill accumulated in portions.

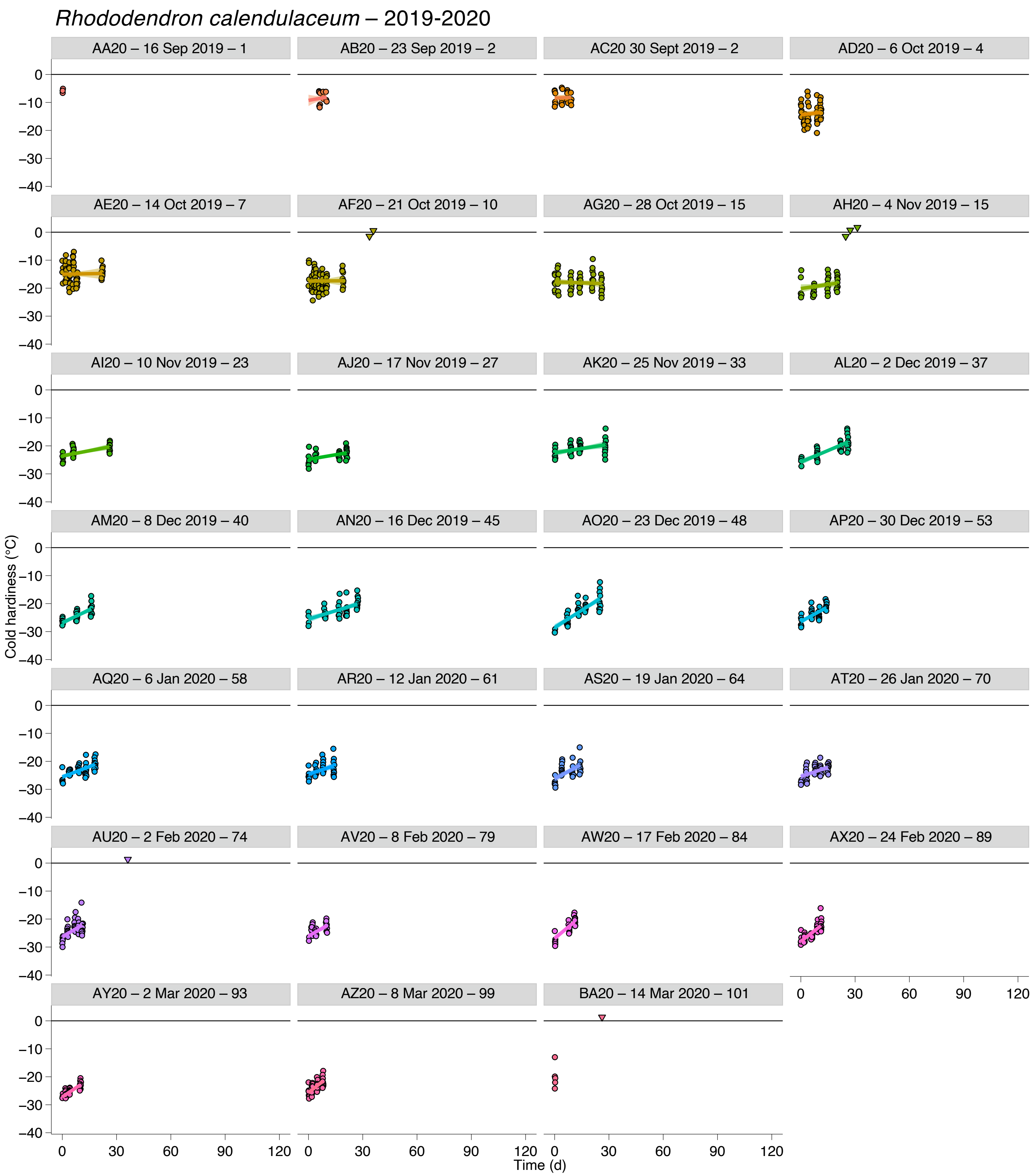


**Supporting Information Figure S32. Deacclimation and budbreak of *Rhododendron calendulaceum* buds collected during the 2019-2020 dormant season.** Loss of cold hardiness (circles) and budbreak (upside down triangles at 0 °C line) for *Rhododendron calendulaceum*. Each graph is identified by collection name, date of collection, and chill accumulated in portions.

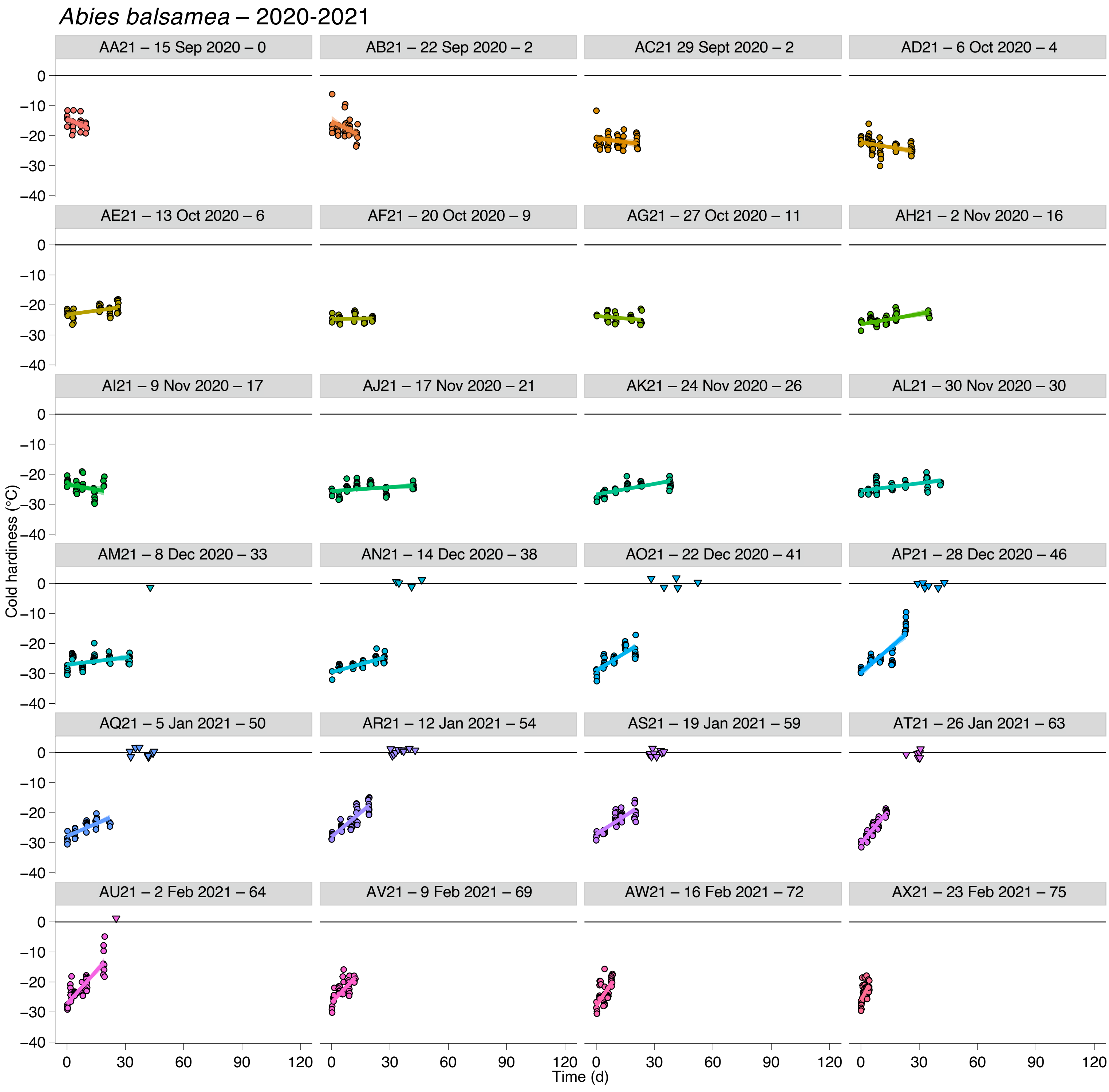


**Supporting Information Figure S33. Deacclimation and budbreak of *Abies balsamea* buds collected during the 2020-2021 dormant season.** Loss of cold hardiness (circles) and budbreak (upside down triangles at 0 °C line) for *Abies balsamea*. Each graph is identified by collection name, date of collection, and chill accumulated in portions.

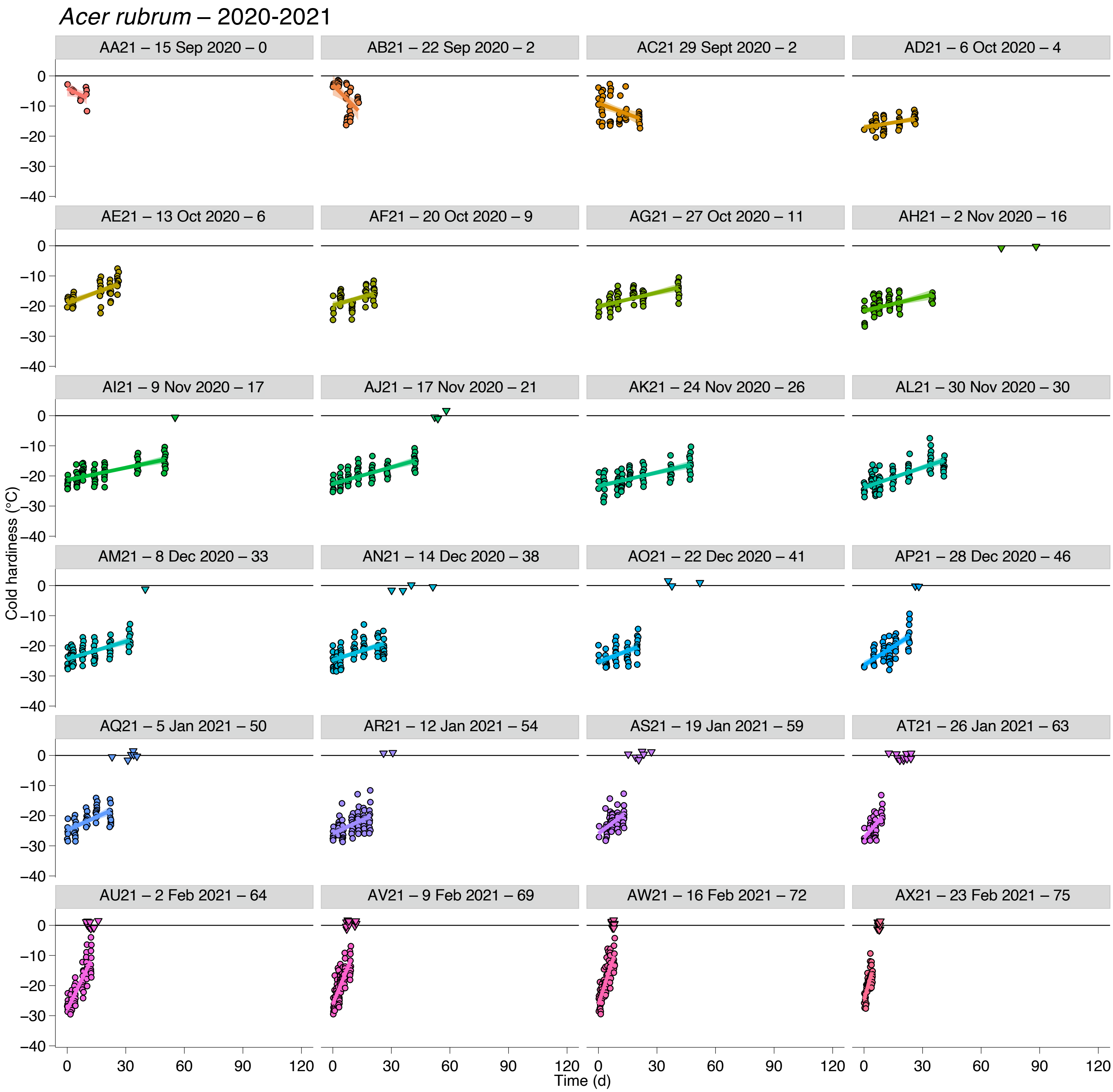


**Supporting Information Figure S34. Deacclimation and budbreak of *Acer rubrum* buds collected during the 2020-2021 dormant season.** Loss of cold hardiness (circles) and budbreak (upside down triangles at 0 °C line) for *Acer rubrum*. Each graph is identified by collection name, date of collection, and chill accumulated in portions.

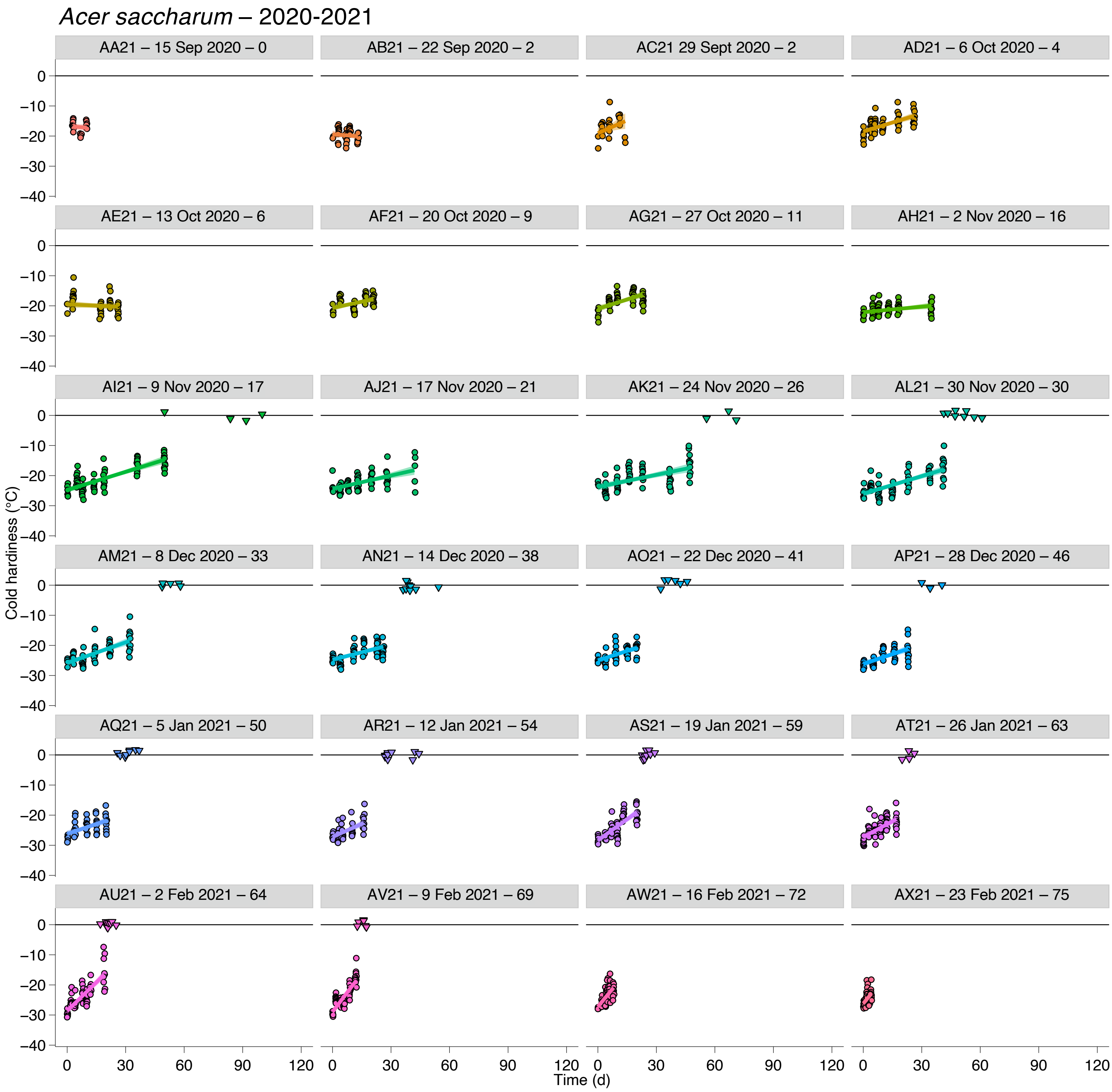


**Supporting Information Figure S35. Deacclimation and budbreak of *Acer saccharum* buds collected during the 2020-2021 dormant season.** Loss of cold hardiness (circles) and budbreak (upside down triangles at 0 °C line) for *Acer saccharum*. Each graph is identified by collection name, date of collection, and chill accumulated in portions.

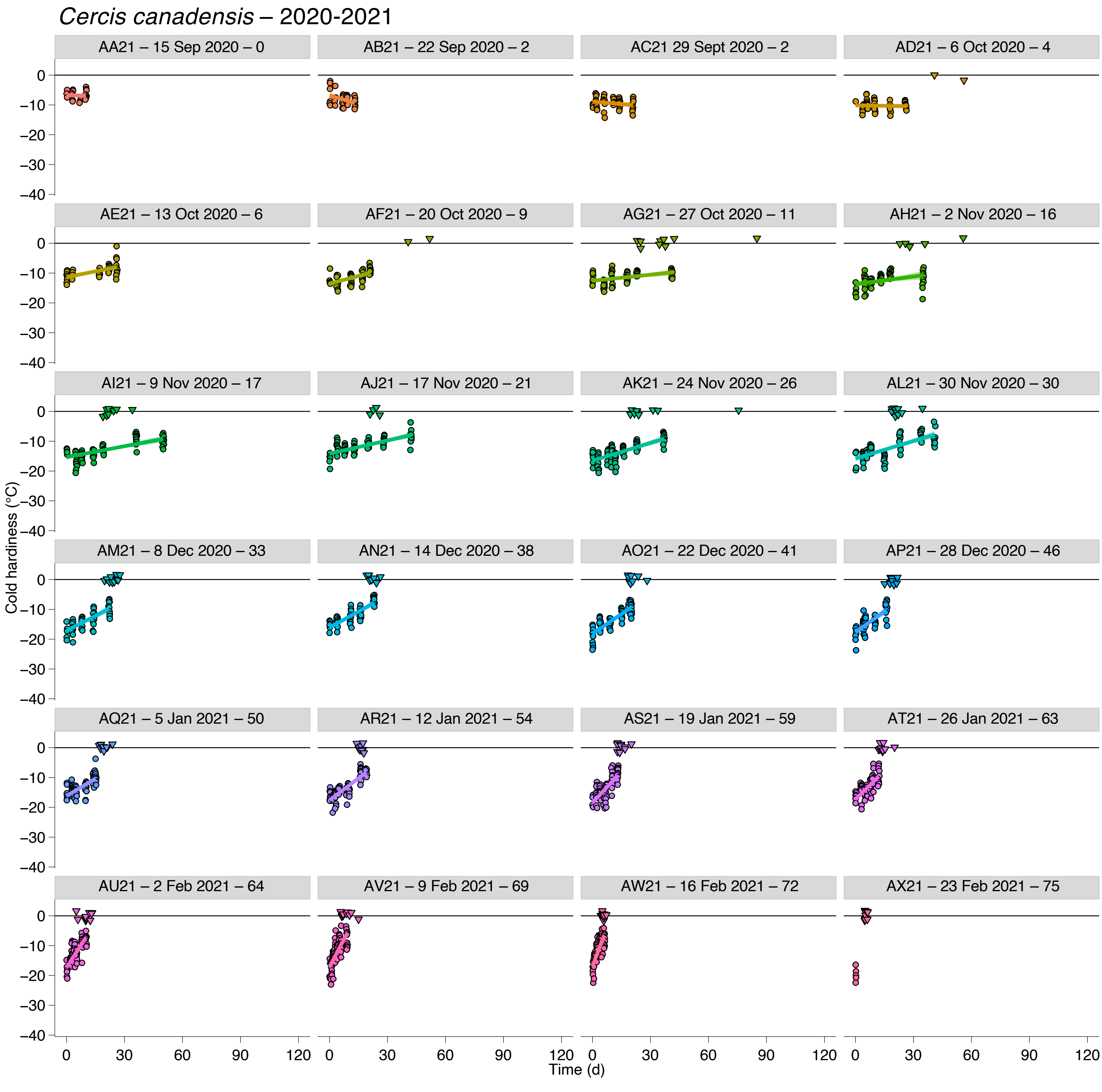


**Supporting Information Figure S36. Deacclimation and budbreak of *Cercis canadensis* buds collected during the 2020-2021 dormant season.** Loss of cold hardiness (circles) and budbreak (upside down triangles at 0 °C line) for *Cercis canadensis*. Each graph is identified by collection name, date of collection, and chill accumulated in portions.

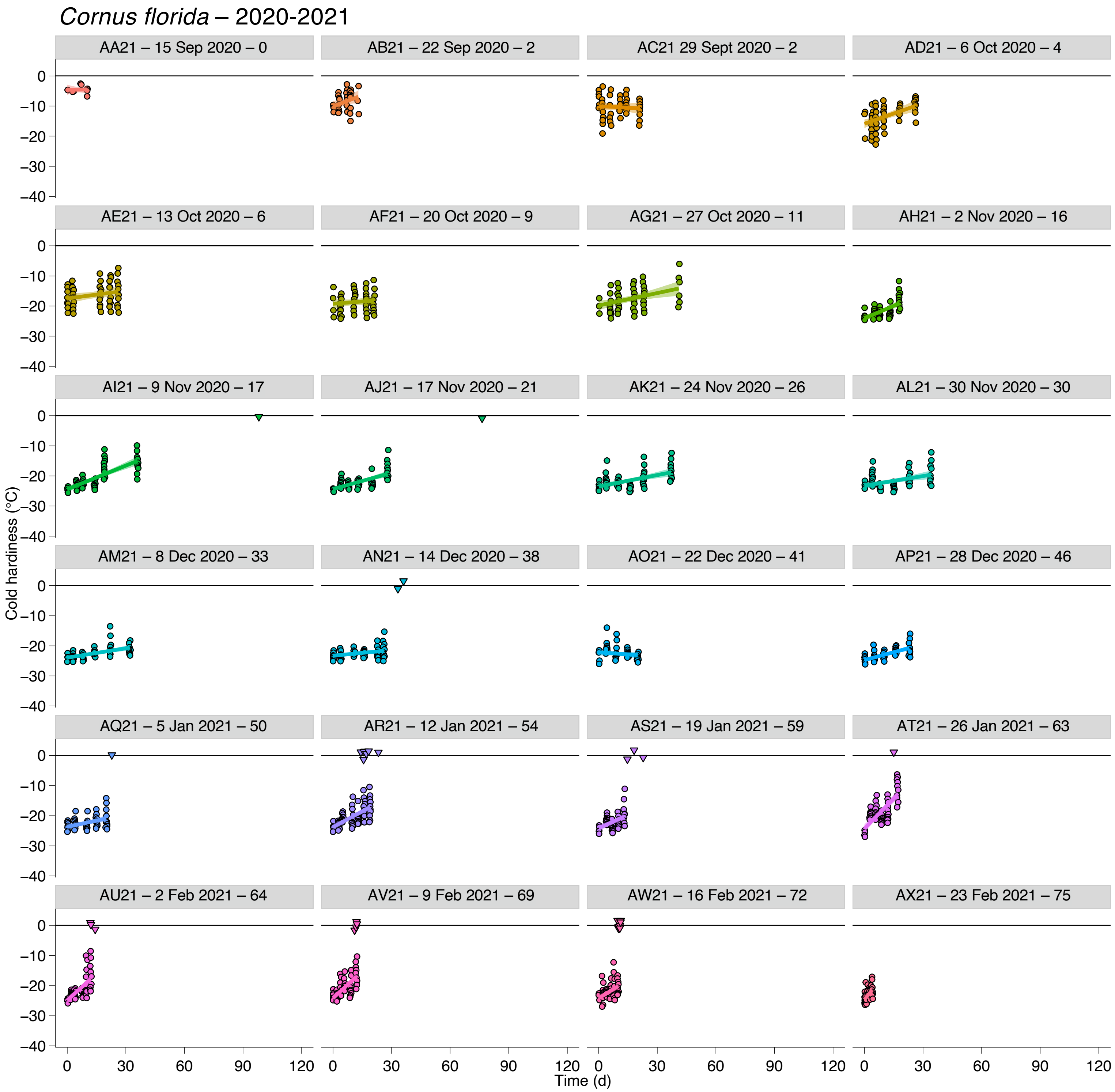


**Supporting Information Figure S37. Deacclimation and budbreak of *Cornus florida* buds collected during the 2020-2021 dormant season.** Loss of cold hardiness (circles) and budbreak (upside down triangles at 0 °C line) for *Cornus florida*. Each graph is identified by collection name, date of collection, and chill accumulated in portions.

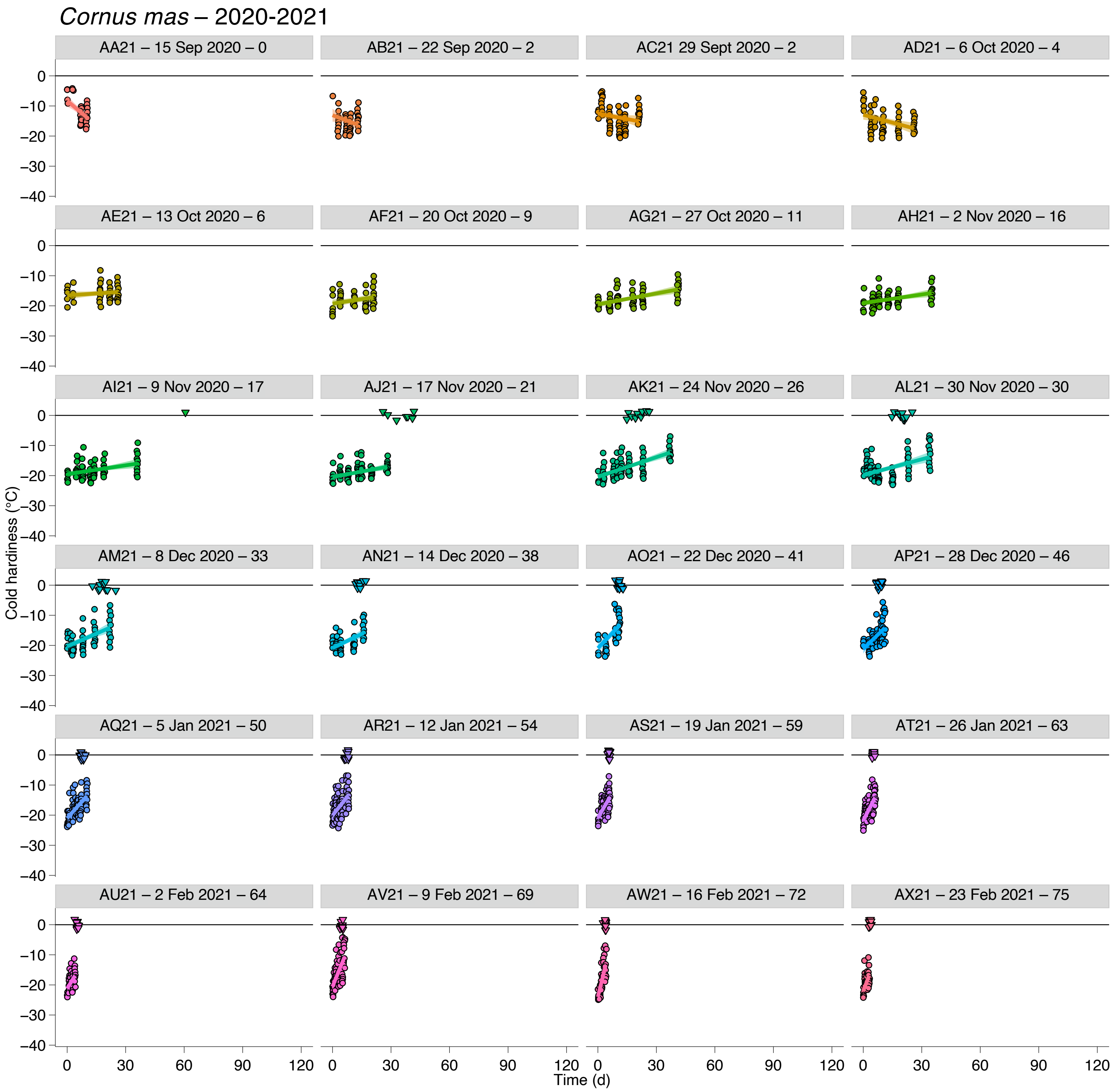


**Supporting Information Figure S38. Deacclimation and budbreak of *Cornus mas* buds collected during the 2020-2021 dormant season.** Loss of cold hardiness (circles) and budbreak (upside down triangles at 0 °C line) for *Cornus mas*. Each graph is identified by collection name, date of collection, and chill accumulated in portions.

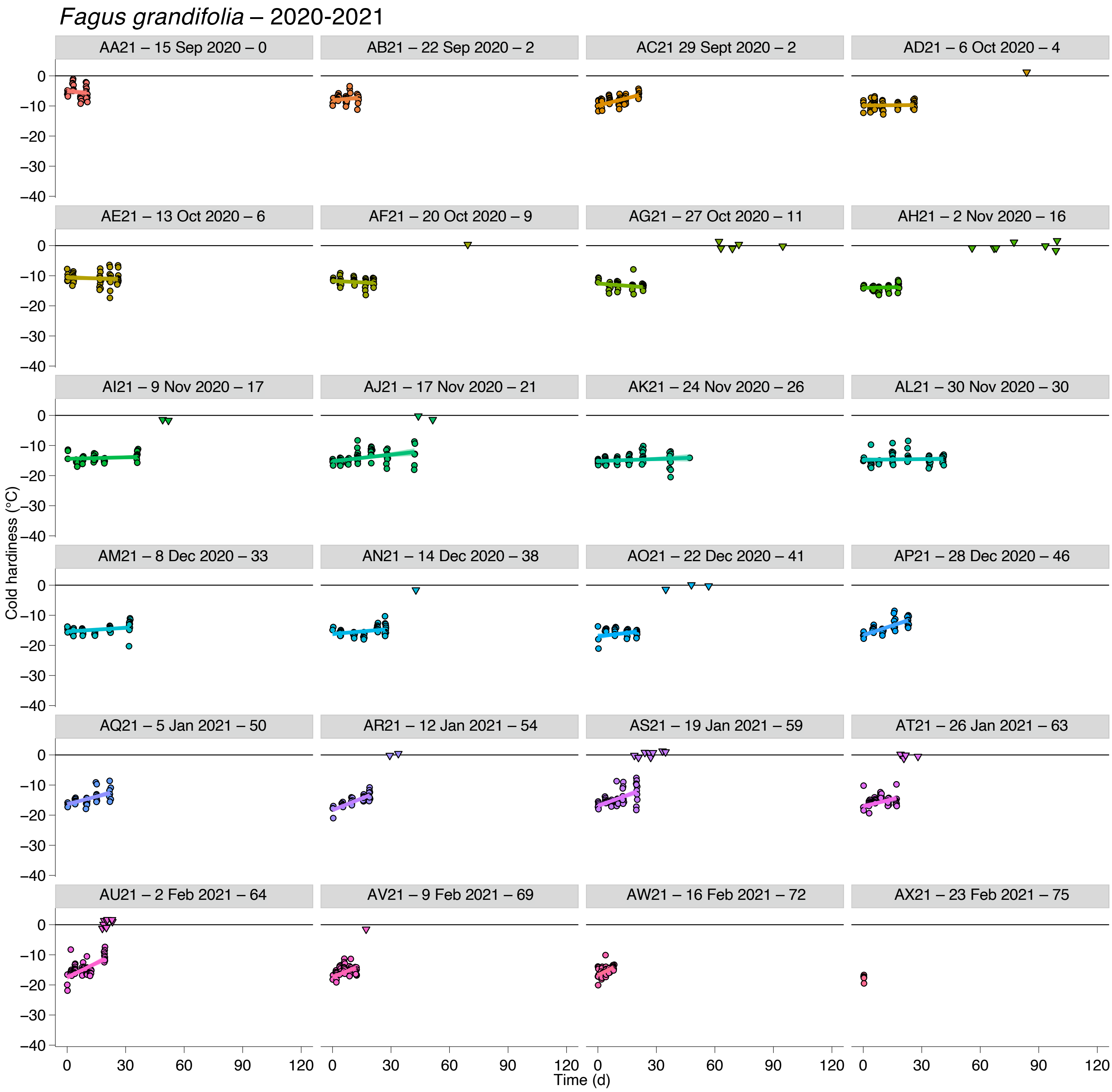


**Supporting Information Figure S39. Deacclimation and budbreak of *Fagus grandifolia* buds collected during the 2020-2021 dormant season.** Loss of cold hardiness (circles) and budbreak (upside down triangles at 0 °C line) for *Fagus grandifolia*. Each graph is identified by collection name, date of collection, and chill accumulated in portions.

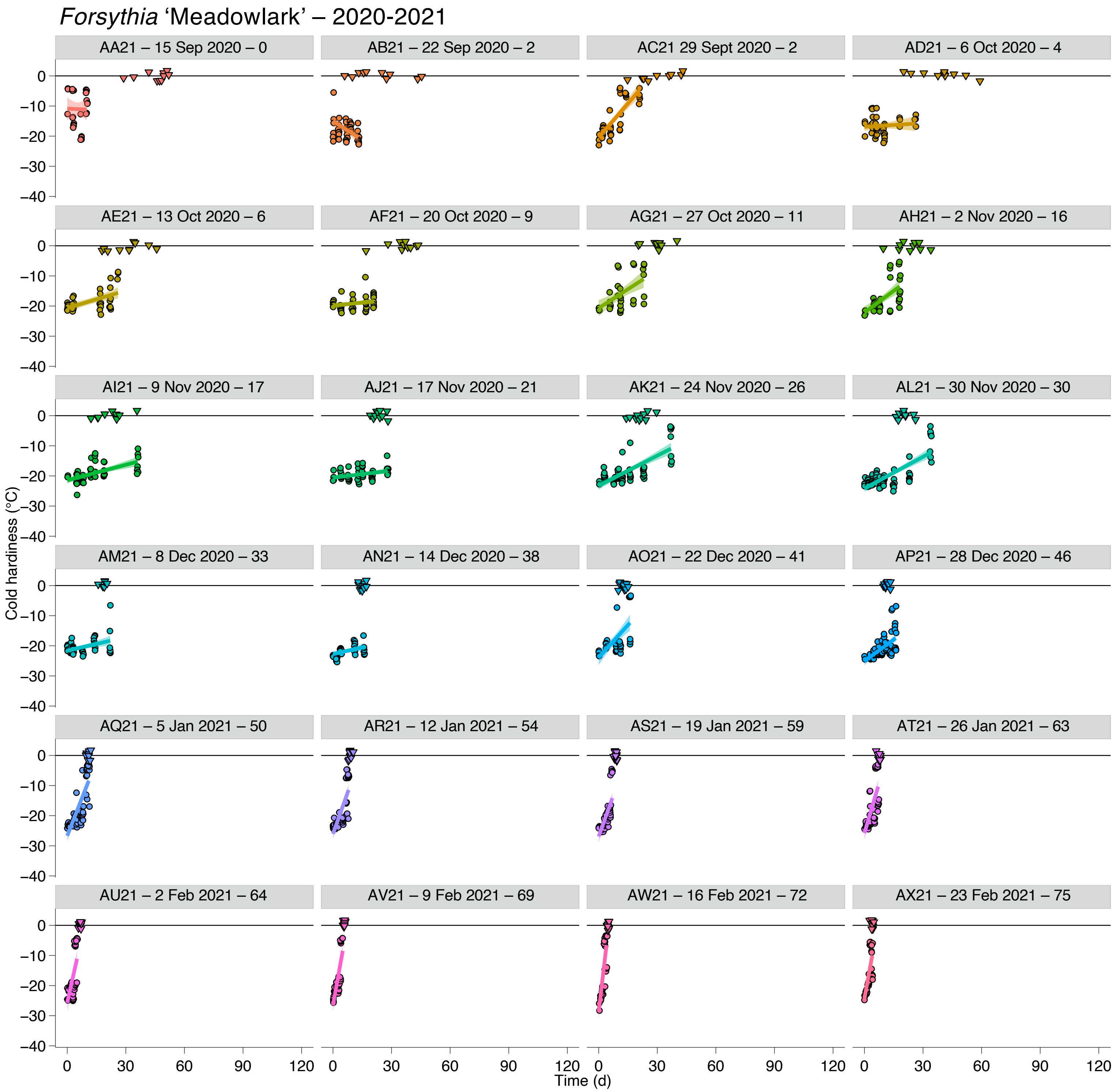


**Supporting Information Figure S40. Deacclimation and budbreak of *Forsythia* 'Meadowlark' buds collected during the 2020-2021 dormant season.** Loss of cold hardiness (circles) and budbreak (upside down triangles at 0 °C line) for *Forsythia* 'Meadowlark'. Each graph is identified by collection name, date of collection, and chill accumulated in portions.

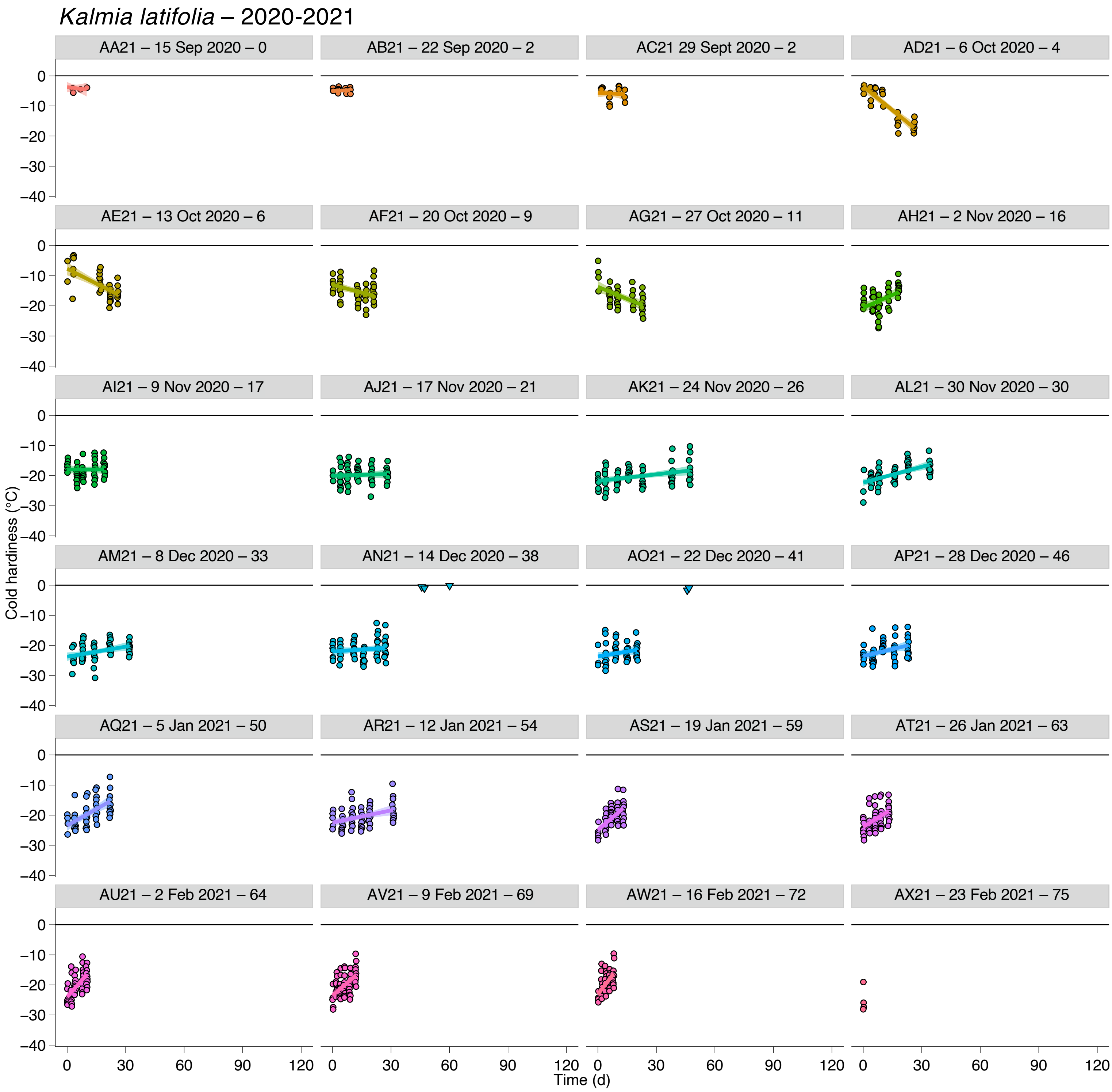


**Supporting Information Figure S41. Deacclimation and budbreak of *Kalmia latifolia* buds collected during the 2020-2021 dormant season.** Loss of cold hardiness (circles) and budbreak (upside down triangles at 0 °C line) for *Kalmia latifolia*. Each graph is identified by collection name, date of collection, and chill accumulated in portions.

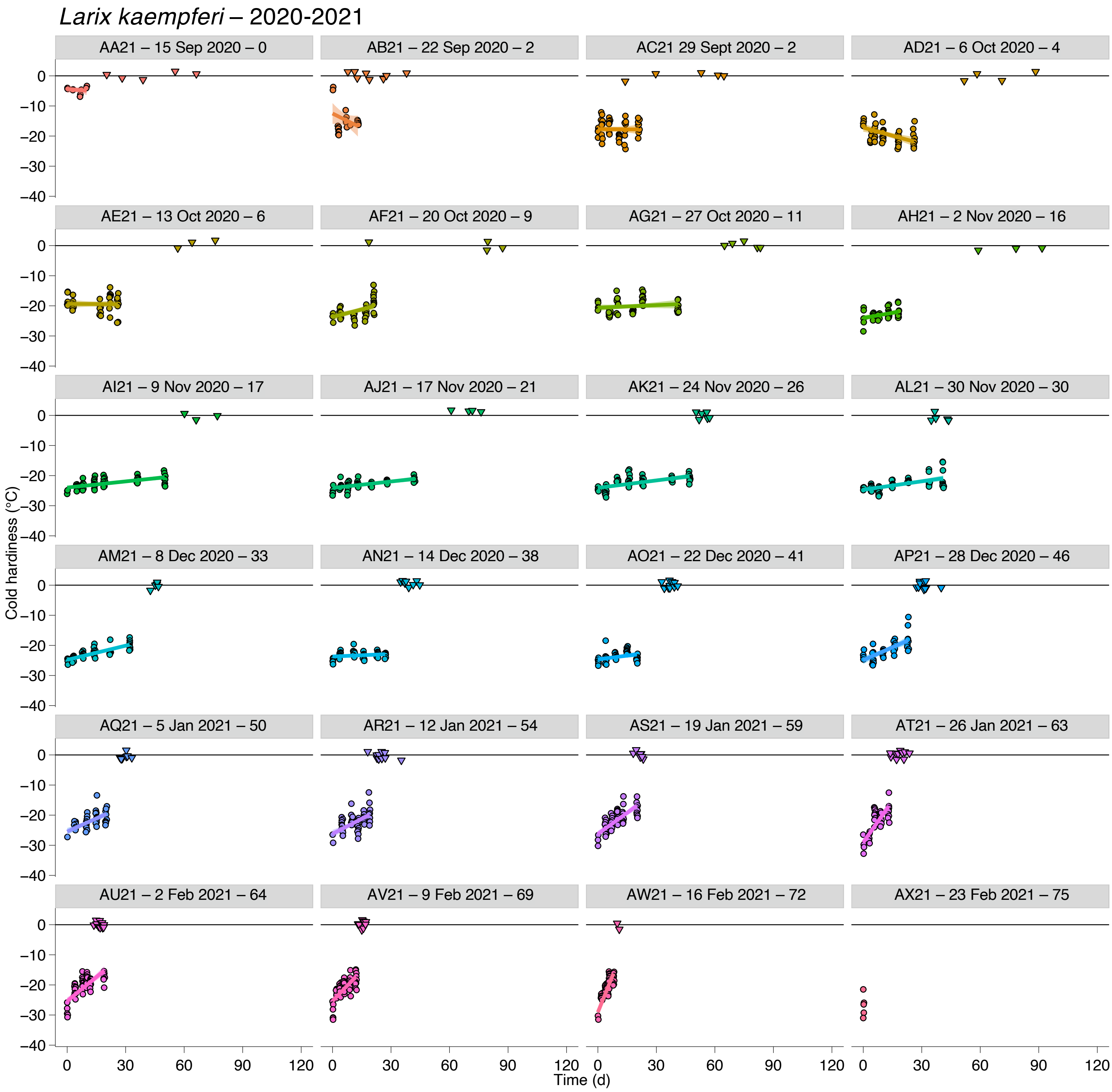


**Supporting Information Figure S42. Deacclimation and budbreak of *Larix kaempferi* buds collected during the 2020-2021 dormant season.** Loss of cold hardiness (circles) and budbreak (upside down triangles at 0 °C line) for *Larix kaempferi*. Each graph is identified by collection name, date of collection, and chill accumulated in portions.

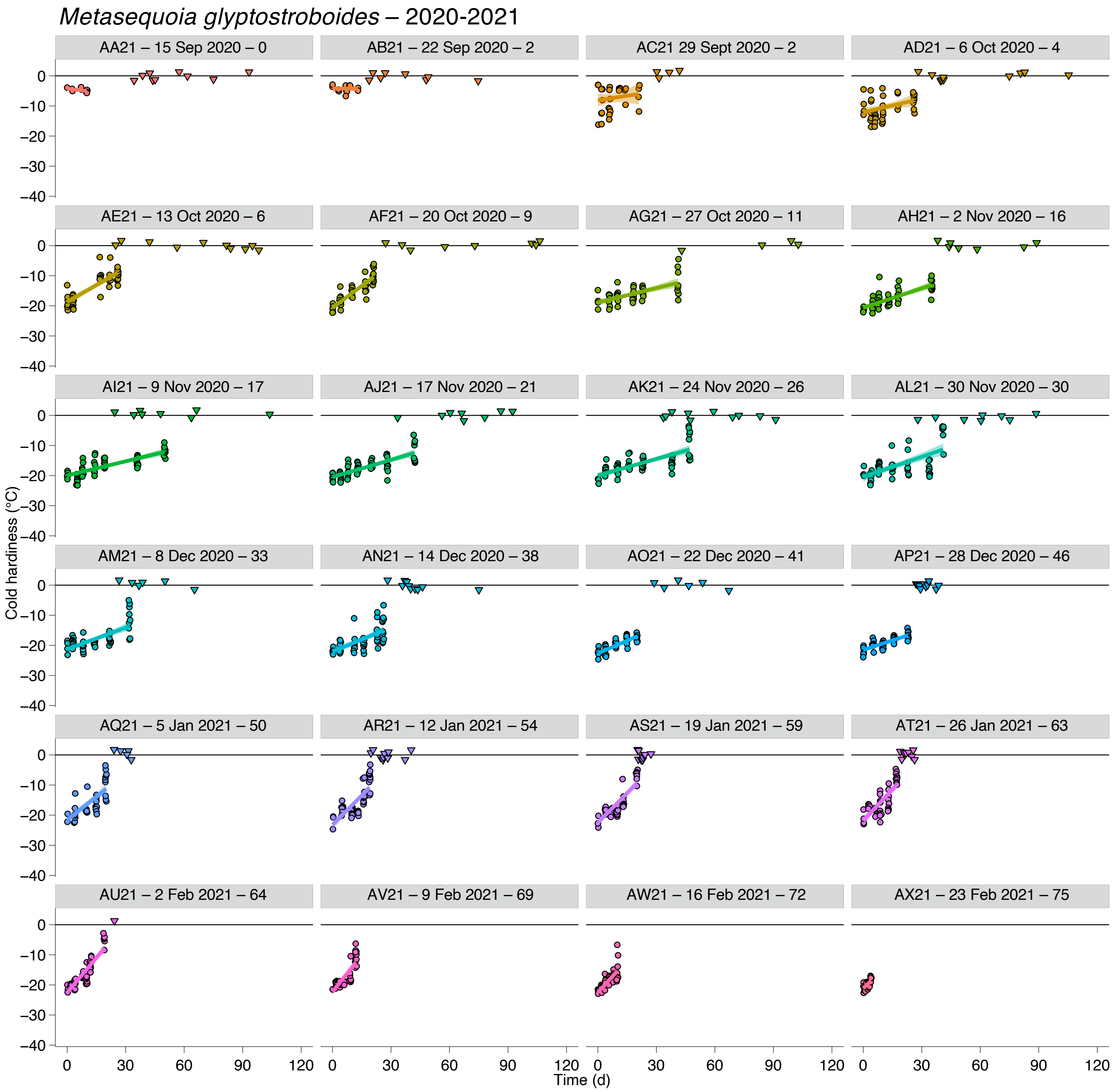


**Supporting Information Figure S43. Deacclimation and budbreak of *Metasequoia glyptostroboides* buds collected during the 2020-2021 dormant season.** Loss of cold hardiness (circles) and budbreak (upside down triangles at 0 °C line) for *Metasequoia glyptostroboides*. Each graph is identified by collection name, date of collection, and chill accumulated in portions.

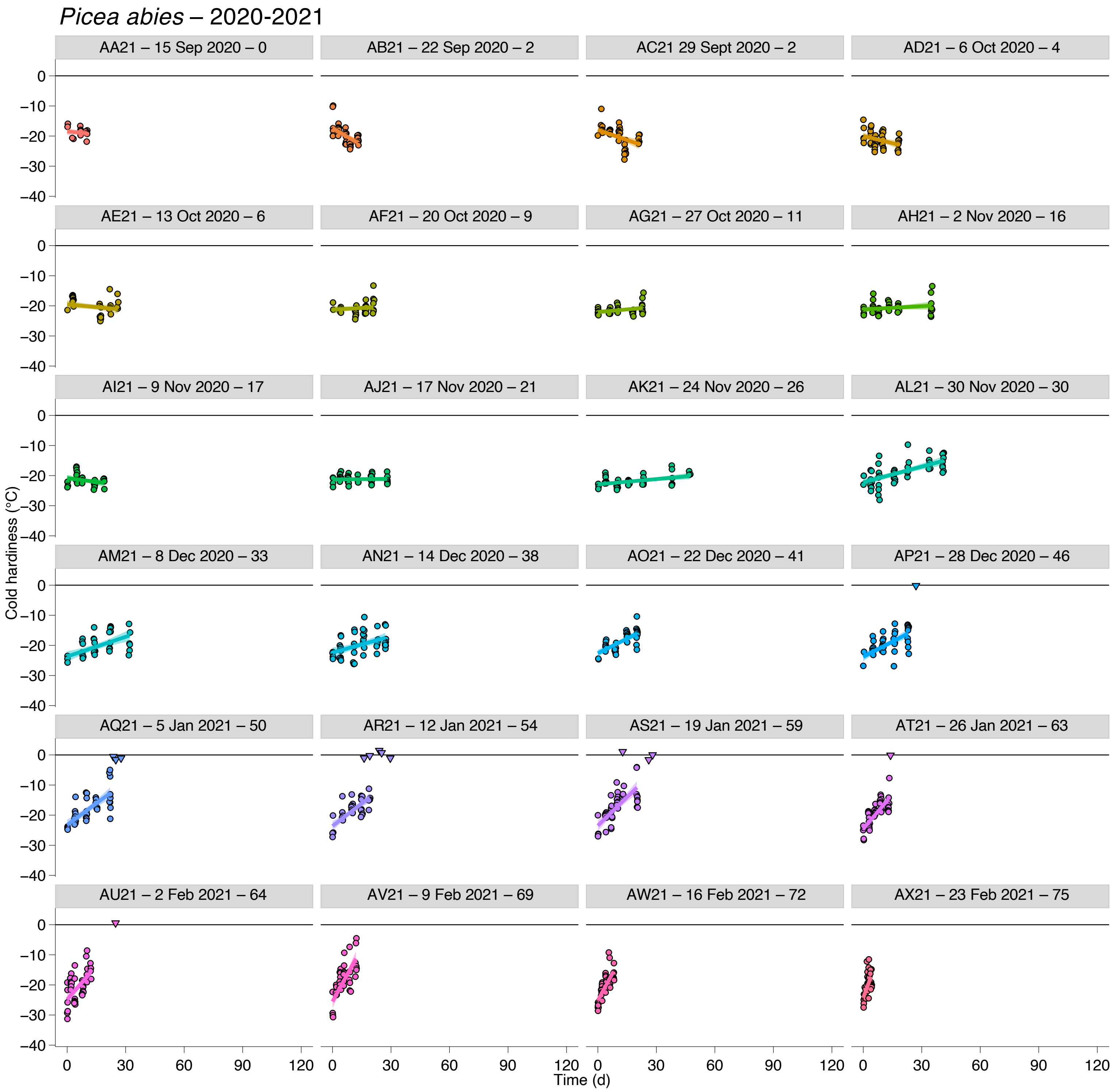


**Supporting Information Figure S44. Deacclimation and budbreak of *Picea abies* buds collected during the 2020-2021 dormant season.** Loss of cold hardiness (circles) and budbreak (upside down triangles at 0 °C line) for *Picea abies*. Each graph is identified by collection name, date of collection, and chill accumulated in portions.

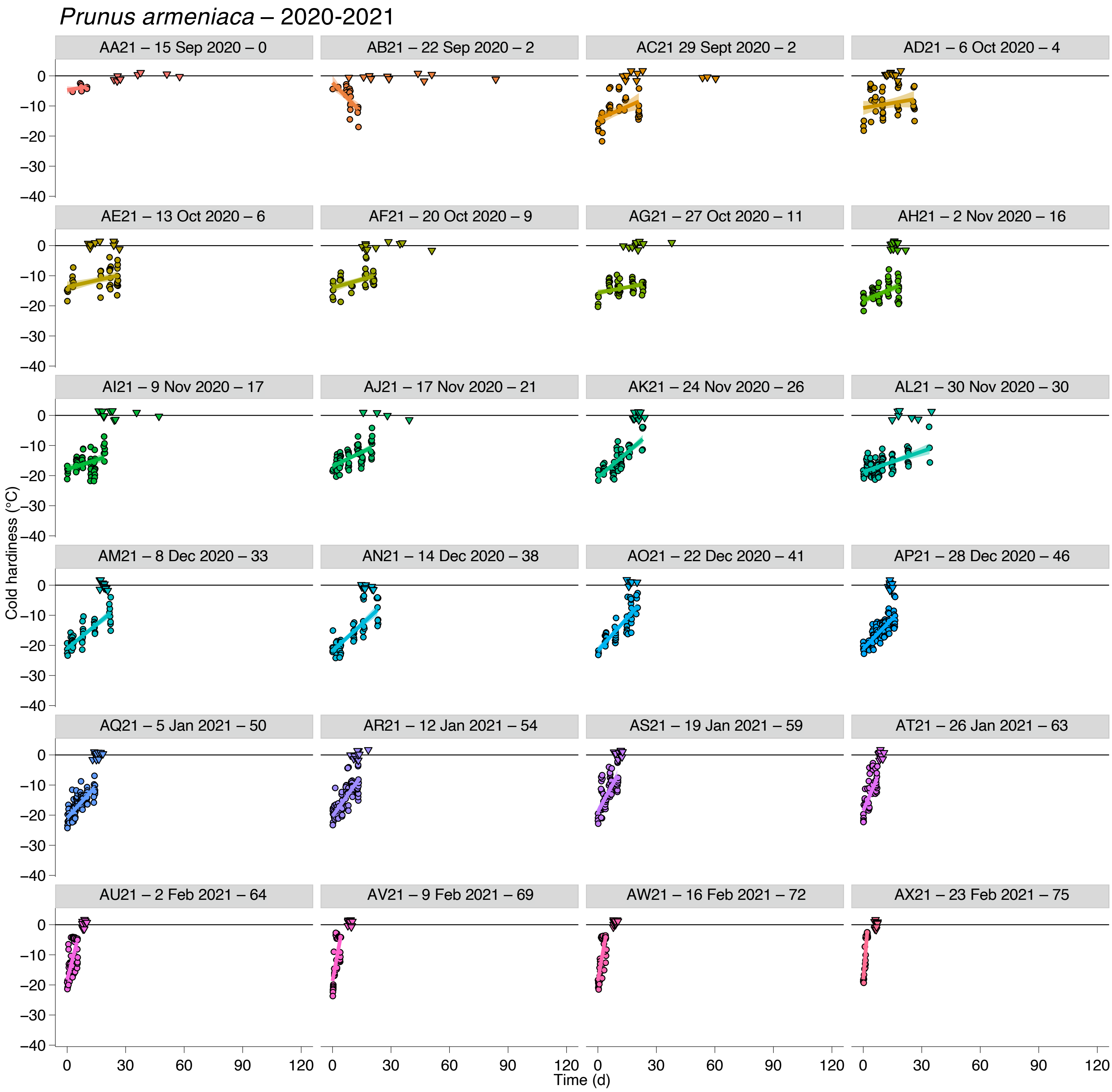


**Supporting Information Figure S45. Deacclimation and budbreak of *Prunus armeniaca* buds collected during the 2020-2021 dormant season.** Loss of cold hardiness (circles) and budbreak (upside down triangles at 0 °C line) for *Prunus armeniaca*. Each graph is identified by collection name, date of collection, and chill accumulated in portions.

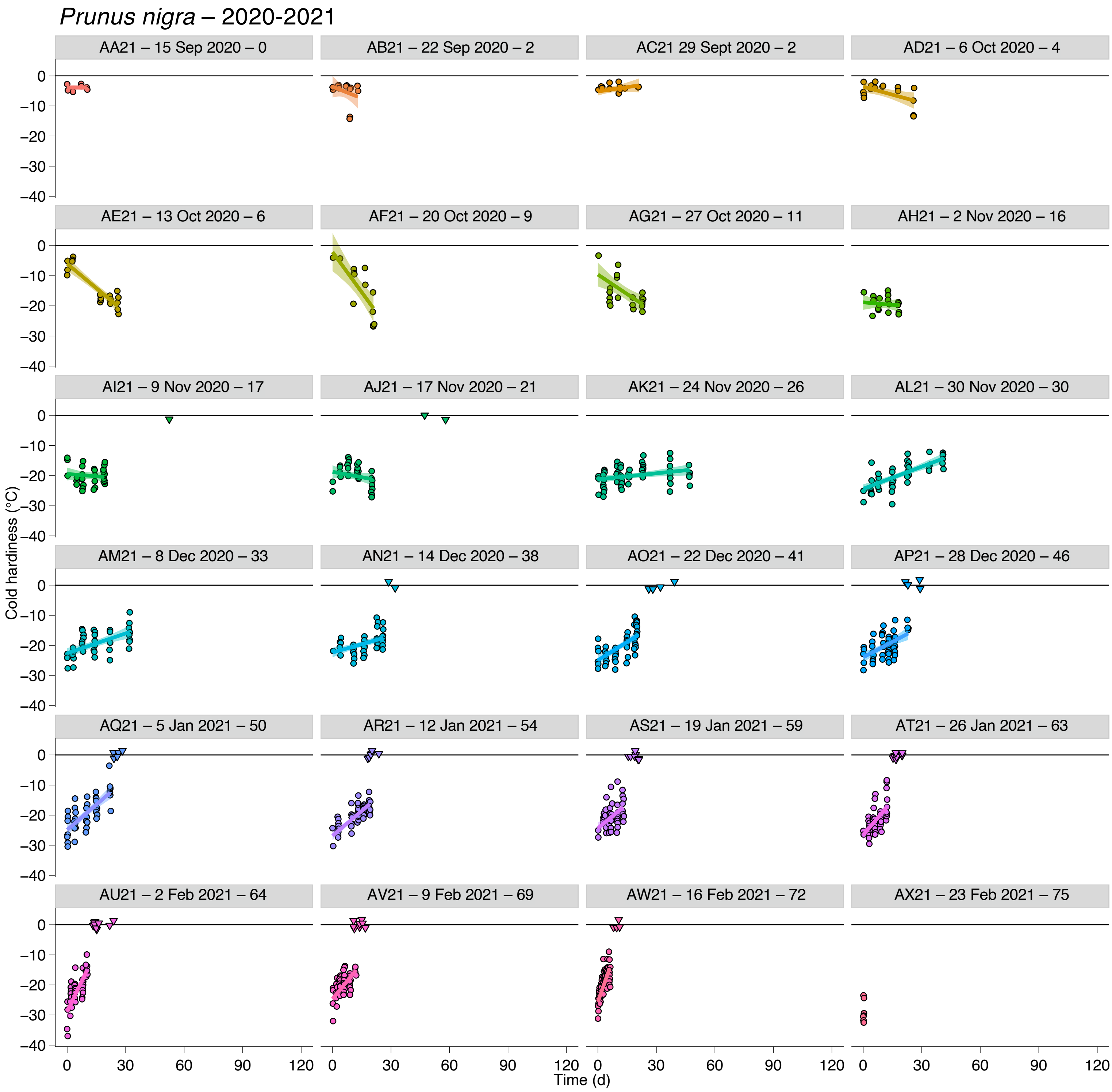


**Supporting Information Figure S46. Deacclimation and budbreak of *Prunus nigra* buds collected during the 2020-2021 dormant season.** Loss of cold hardiness (circles) and budbreak (upside down triangles at 0 °C line) for *Prunus nigra*. Each graph is identified by collection name, date of collection, and chill accumulated in portions.

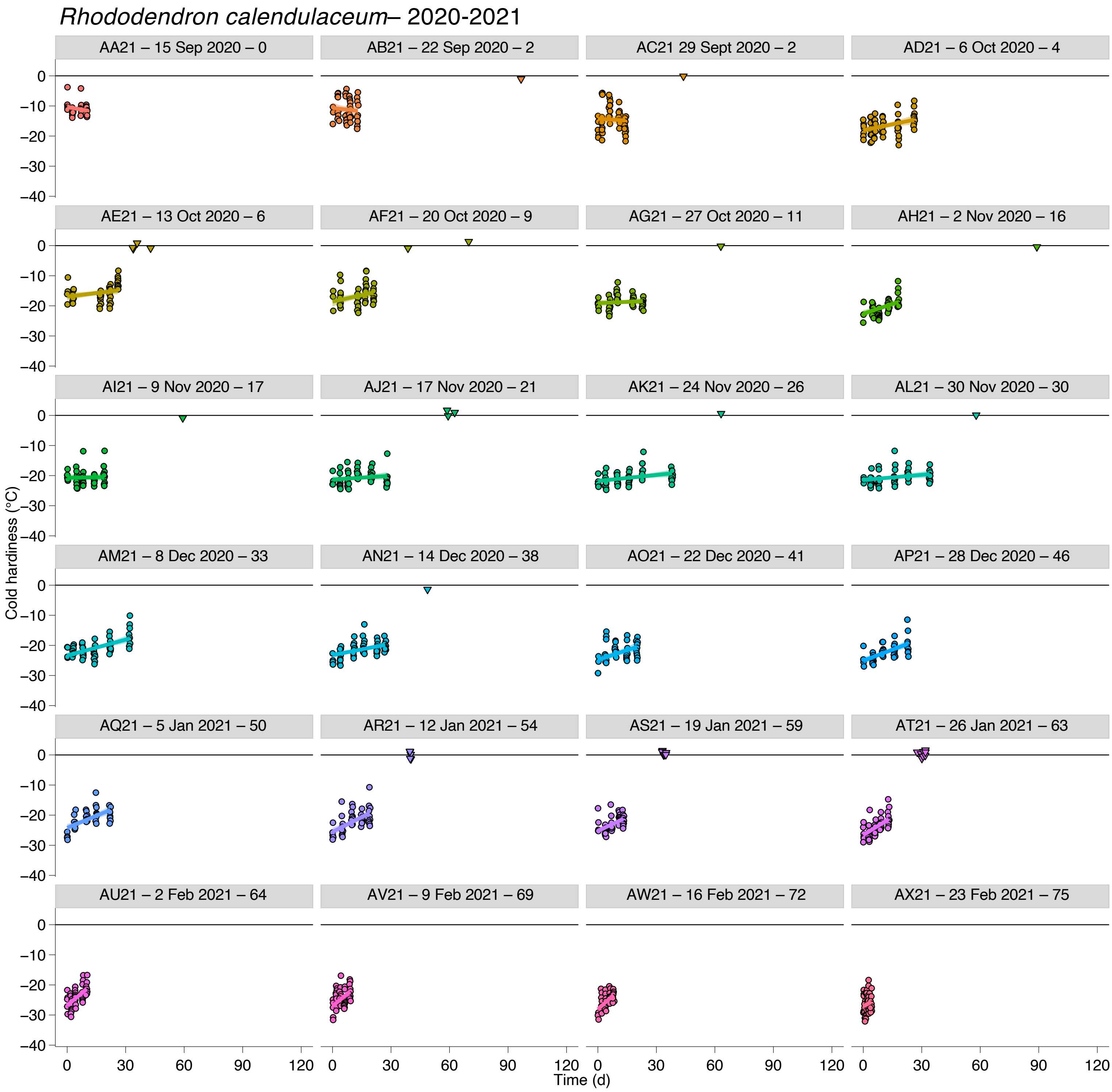


**Supporting Information Figure S47. Deacclimation and budbreak of *Rhododendron calendulaceum* buds collected during the 2020-2021 dormant season.** Loss of cold hardiness (circles) and budbreak (upside down triangles at 0 °C line) for *Rhododendron calendulaceum*. Each graph is identified by collection name, date of collection, and chill accumulated in portions.

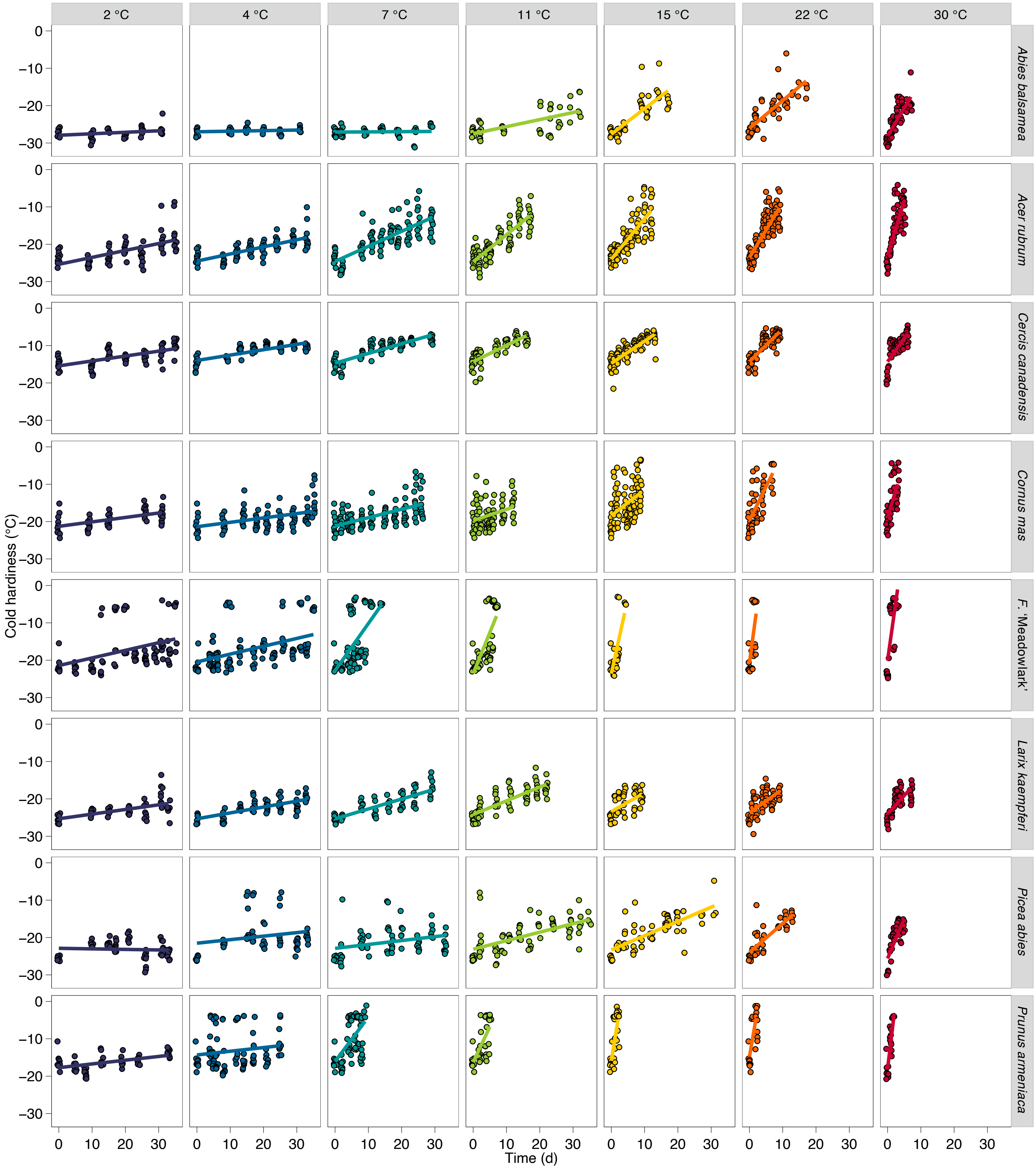


**Supporting Information Figure S48. Deacclimation at different temperatures.** Loss of cold hardiness over time of buds from eight species (rows) at seven different temperatures (columns), collected on 12 Feb 2020 (82 chill portions accumulated).

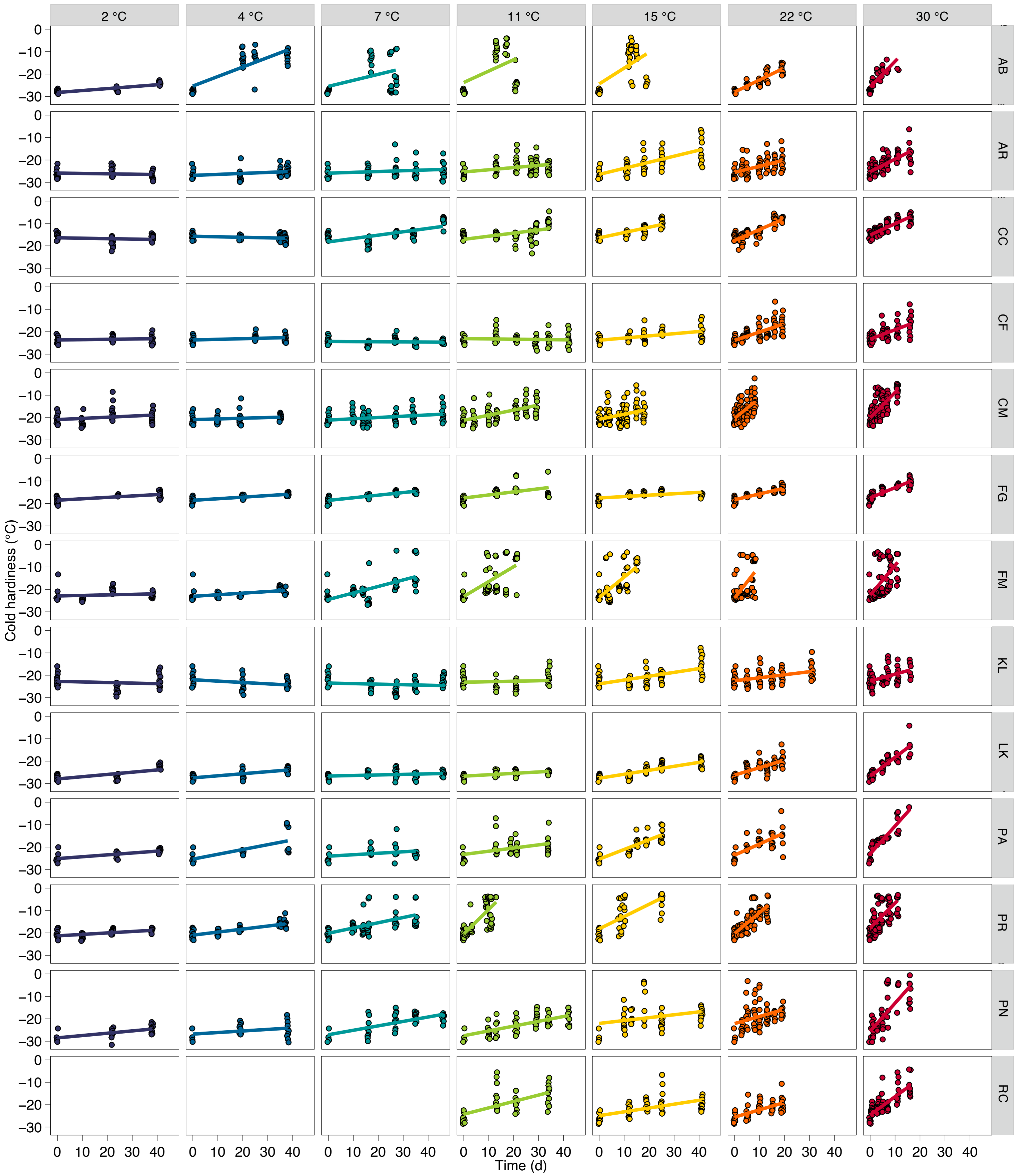


**Supporting Information Figure S49. Deacclimation at different temperatures.** Loss of cold hardiness over time of buds from nine species (rows) at seven different temperatures (columns), collected on 12 Jan 2021 (54 chill portions accumulated).

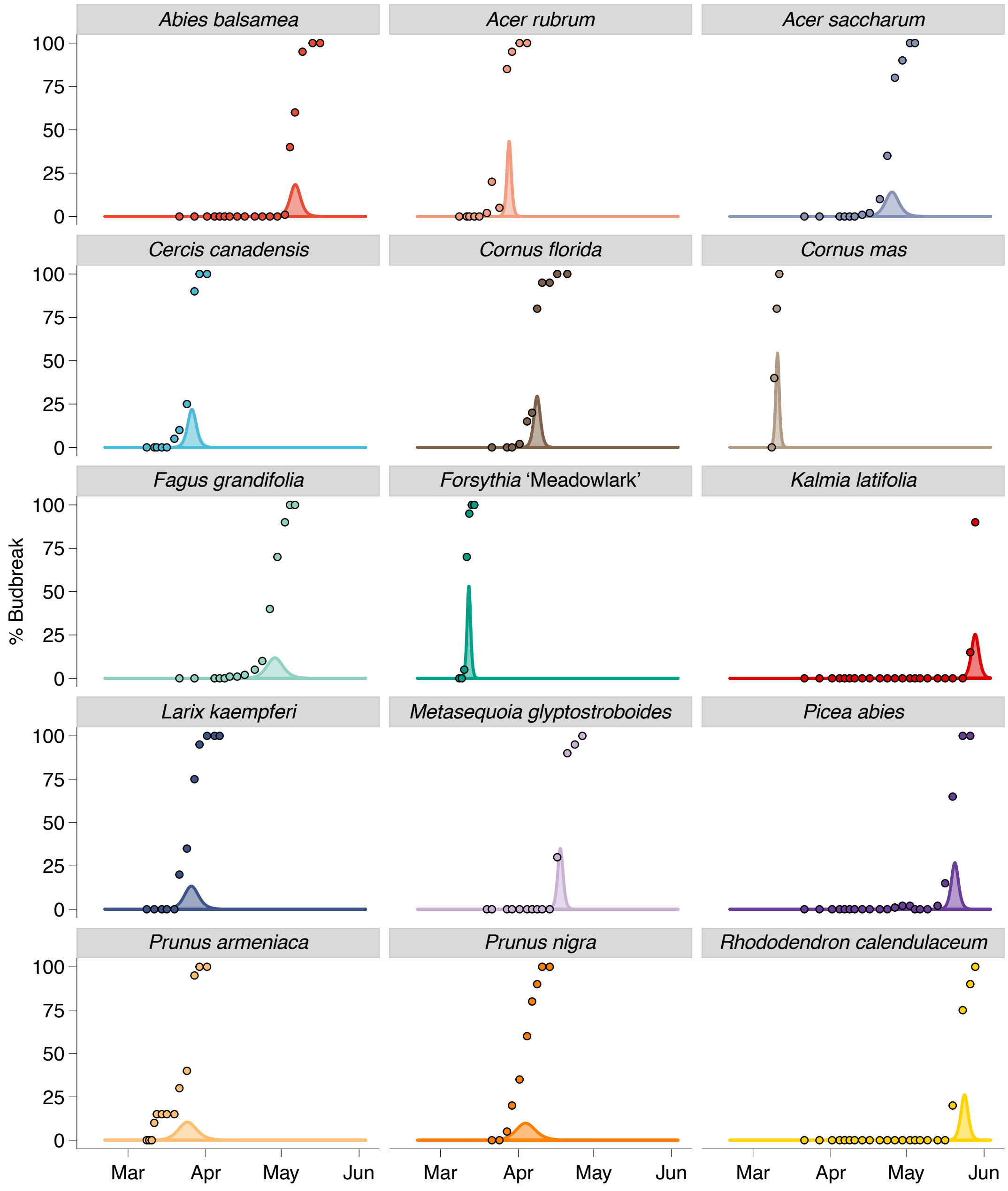


**Supporting Information Figure S50. Field budbreak.** Measured percent budbreak (circles) and density plots based on probability density function of as sigmoid response to time of different species at the Arnold Arboretum of Harvard University in the spring of 2020.
