## Supplementary material for "Woody species do not differ in dormancy progression: differences in time to budbreak due to forcing and cold hardiness": Table S1

**Supporting Information Table S1. Description of species in this study.**

| Species | Abbreviation | Common name | Accession | Family | Habit | Bud observed^†^ | Description of budbreak | Total number of cold hardiness measurements |
| --- | --- | --- | --- | --- | --- | --- | --- | --- |
| *Abies balsamea* | AB | Balsam  Fir | 610-93*C | Pinaceae | Evergreen | V | Bud scales separate and needle tips show | 3,003 |
| *Acer rubrum* | AR | Red  Maple | 688-39*H | Sapindaceae | Deciduous | R | Flower bud opens | 3,286 |
| *Acer saccharum* | AS | Sugar  Maple | 655-93*G | Sapindaceae | Deciduous | V | Multiple bud scales separate and leaf tips are visible | 2,054 |
| *Cercis canadensis* | CC | Eastern  Redbud | 403-95*B | Fabaceae | Deciduous | R | Multiple bud scales separate and flower tips are visible | 3,174 |
| *Cornus florida* | CF | Flowering  Dogwood | 216-2014*A | Cornaceae | Deciduous | R | Two pairs of bud scales separate and flowers are visible | 2,683 |
| *Cornus mas* | CM | Cornelian-Cherry | 645-79*A | Cornaceae | Deciduous | R | Two pairs of bud scales separate and flowers are visible | 3,386 |
| *Fagus grandifolia* | FG | American  Beech | 6XX-2008*A | Fagaceae | Deciduous | V | Bud swells and scales fall to show leaves/flowers | 2,323 |
| *Forsythia* × *‘Meadowlark’* | FM | Forsythia | 198-2007*MASS-A | Oleaceae | Deciduous | V/R | Flower bud elongates, points downwards and petals are visible and yellow. For vegetative, vegetative growth starts below a floral bud. | 2,904 |
| *Kalmia latifolia* | KL | Mountain  Laurel | 440-66*A;  150-58*A;  728-54*B;  728-54*D | Ericaceae | Evergreen | R | Inflorescence elongates, at least one flower expands into “balloon” stage | 2,521 |
| *Larix kaempferi* | LK | Japanese  Larch | 11276*F | Pinaceae | Deciduous | V | Multiple bud scales separate and leaf tips are visible | 2,730 |
| *Metasequoia glyptostroboides* | MG | Dawn  Redwood | 524-48*Z | Cupressaceae | Deciduous | V | Multiple bud scales separate and leaf tips are visible | 2,541 |
| *Picea abies* | PA | Norway  Spruce | 23026*M | Pinaceae | Evergreen | V | Multiple bud scales separate and leaf tips are visible | 2,652 |
| *Prunus armeniaca* | PR | Apricot | 382-92*B | Rosaceae | Deciduous | V/R | Multiple bud scales separate; for reproductive, sepals open and petals are visible; for vegetative, leaf tips are visible. | 2,829 |
| *Prunus nigra* | PN | Canadian  Plum | 107-78*A | Rosaceae | Deciduous | V/R | Multiple bud scales separate; for reproductive, sepals open and petals are visible; for vegetative, leaf tips are visible. | 1840 |
| *Rhododendron calendulaceum* | RC | Flame  Azalea | 17759*A; 656-70*MASS; 19483*MASS | Ericaceae | Deciduous | R | Multiple bud scales separate and flower tips are visible | 2630 |

^†^V – Vegetative; R – Reproductive.

**Supporting Information Table S2. Details on field collections.**

| **2019-2020** | | | |  | **2020-2021** | | | |
| --- | --- | --- | --- | --- | --- | --- | --- | --- |
| **Collection** | **Date of Collection** | **Chill portions accumulated** | **Deacclimation and budbreak assay** |  | **Collection** | **Date of Collection** | **Chill portions accumulated** | **Deacclimation and budbreak assay** |
| AA20 | 16 Sep 2019 | 1 | × |  | AA21 | 15 Sep 2020 | 0 | × |
| AB20 | 23 Sep 2019 | 2 | × |  | AB21 | 22 Sep 2020 | 2 | × |
| AC20 | 30 Sep 2019 | 2 | × |  | AC21 | 29 Sep 2020 | 2 | × |
| AD20 | 6 Oct 2019 | 4 | × |  | AD21 | 6 Oct 2020 | 4 | × |
| AE20 | 14 Oct 2019 | 7 | × |  | AE21 | 13 Oct 2020 | 6 | × |
| AF20 | 21 Oct 2019 | 10 | × |  | AF21 | 20 Oct 2020 | 9 | × |
| AG20 | 28 Oct 2019 | 15 | × |  | AG21 | 27 Oct 2020 | 11 | × |
| AH20 | 4 Nov 2019 | 15 | × |  | AH21 | 2 Nov 2020 | 16 | × |
| AI20 | 10 Nov 2019 | 23 | × |  | AI21 | 9 Nov 2020 | 17 | × |
| AJ20 | 17 Nov 2019 | 27 | × |  | AJ21 | 17 Nov 2020 | 21 | × |
| AK20 | 25 Nov 2019 | 33 | × |  | AK21 | 24 Nov 2020 | 26 | × |
| AL20 | 2 Dec 2019 | 37 | × |  | AL21 | 30 Nov 2020 | 30 | × |
| AM20 | 8 Dec 2019 | 40 | × |  | AM21 | 8 Dec 2020 | 33 | × |
| AN20 | 16 Dec 2019 | 45 | × |  | AN21 | 14 Dec 2020 | 38 | × |
| AO20 | 23 Dec 2019 | 48 | × |  | AO21 | 22 Dec 2020 | 41 | × |
| AP20 | 30 Dec 2019 | 53 | × |  | AP21 | 28 Dec 2020 | 46 | × |
| AQ20 | 6 Jan 2020 | 58 | × |  | AQ21 | 5 Jan 2021 | 50 | × |
| AR20 | 12 Jan 2020 | 61 | × |  | AR21 | 12 Jan 2021 | 54 | × |
| AS20 | 19 Jan 2020 | 64 | × |  | AS21 | 19 Jan 2021 | 59 | × |
| AT20 | 26 Jan 2020 | 70 | × |  | AT21 | 26 Jan 2021 | 63 | × |
| AU20 | 2 Feb 2020 | 74 | × |  | AU21 | 2 Feb 2021 | 64 | × |
| AV20 | 8 Feb 2020 | 79 | × |  | AV21 | 9 Feb 2021 | 69 | × |
| AW20 | 17 Feb 2020 | 84 | × |  | AW21 | 16 Feb 2021 | 72 | × |
| AX20 | 24 Feb 2020 | 89 | × |  | AX21 | 23 Feb 2021 | 75 | × (Not all species) |
| AY20 | 2 Mar 2020 | 93 | × |  | AY21 | 26 Feb 2021 | 77 |  |
| AZ20 | 8 Mar 2020 | 99 | × |  |  |  |  |  |
| BA20 | 14 Mar 2020 | 101 | × (Not all species) |  |  |  |  |  |
| BB20 | 18 Mar 2020 | 105 |  |  |  |  |  |  |
| BC20 | 19 Mar 2020 | 106 |  |  |  |  |  |  |
| BD20 | 22 Mar 2020 | 107 |  |  |  |  |  |  |
| BE20 | 27 Mar 2020 | 111 |  |  |  |  |  |  |
| BF20 | 4 Apr 2020 | 117 |  |  |  |  |  |  |
| BG20 | 8 Apr 2020 | 119 |  |  |  |  |  |  |
| BH20 | 11 Apr 2020 | 122 |  |  |  |  |  |  |
| BI20 | 13 Apr 2020 | 123 |  |  |  |  |  |  |
| BJ20 | 16 Apr 2020 | 125 |  |  |  |  |  |  |
| BK20 | 20 Apr 2020 | 128 |  |  |  |  |  |  |
| BL20 | 23 Apr 2020 | 130 |  |  |  |  |  |  |
| BM20 | 26 Apr 2020 | 132 |  |  |  |  |  |  |
| BN20 | 29 Apr 2020 | 134 |  |  |  |  |  |  |
| BO20 | 2 May 2020 | 135 |  |  |  |  |  |  |
| BP20 | 4 May 2020 | 135 |  |  |  |  |  |  |
| BQ20 | 6 May 2020 | 136 |  |  |  |  |  |  |
| BR20 | 9 May 2020 | 138 |  |  |  |  |  |  |
| BS20 | 13 May 2020 | 142 |  |  |  |  |  |  |
| BT20 | 16 May 2020 | 142 |  |  |  |  |  |  |
| BU20 | 19 May 2020 | 143 |  |  |  |  |  |  |
| BV20 | 23 May 2020 | 144 |  |  |  |  |  |  |

**Supporting Information Table S3. Estimates associated with regressions of deacclimation rate in response to temperature and deacclimation in response to chill accumulation for each species.**

| Species | Temperature response ($\max\text{k}_{\text{deacc}}$)^†^ | | | | |  | Chill response (*Ψ_deacc_*)^‡^ | | | |
| --- | --- | --- | --- | --- | --- | --- | --- | --- | --- | --- |
|  | n (different temperature assays) | β_1_ | | β_2_ | β_3_ |  | n (different chill assays) | d | b | c |
| *Abies balsamea* | 14 | 0.0937 | | –0.00516 | 0.0001491 |  | 51 | 0.017 | –4.38 | 55.0 |
| *Acer rubrum* | 14 | 0.1279 | | –0.00312 | 0.0001522 |  | 50 | 0.043 | –11.20 | 70.1 |
| *Acer saccharum* | 1 | 0.0527 | | 0.00000 | 0.0000000 |  | 51 | 0.134 | –5.64 | 73.8 |
| *Cercis canadensis* | 14 | 0.0655 | | 0.00115 | –0.0000307 |  | 50 | 0.080 | –6.62 | 62.5 |
| *Cornus florida* | 7 | 0.0087 | | 0.00475 | –0.0001156 |  | 49 | 0.103 | –6.70 | 68.6 |
| *Cornus mas* | 14 | –0.0485 | | 0.01624 | –0.0002497 |  | 50 | 0.048 | –7.10 | 71.8 |
| *Fagus grandifolia* | 7 | 0.0569 | | –0.00293 | 0.0000759 |  | 50 | 0.105 | –6.53 | 72.8 |
| *Forsythia* ‘Meadowlark’ | 14 | –0.0314 | | 0.04830 | –0.0012354 |  | 50 | 0.029 | –7.57 | 70.0 |
| *Kalmia latifolia* | 7 | –0.0332 | | 0.00326 | 0.0000000 |  | 50 | 0.037 | –3.51 | 59.7 |
| *Larix kaempferi* | 14 | 0.0851 | | –0.00454 | 0.0001476 |  | 49 | 0.045 | –4.65 | 63.9 |
| *Metasequoia glyptostroboides* | 1 | 0.0606 | | 0.00000 | 0.0000000 |  | 50 | 0.118 | –3.96 | 58.2 |
| *Picea abies* | 14 | 0.0658 | | –0.00414 | 0.0002155 |  | 51 | 0.006 | –3.50 | 65.8 |
| *Prunus armeniaca* | 14 | 0.1144 | | 0.02069 | –0.0004248 |  | 50 | 0.027 | –7.74 | 72.8 |
| *Prunus nigra* | 7 | 0.2602 | | –0.02510 | 0.0007594 |  | 49 | 0.039 | –4.62 | 61.9 |
| *Rhododendron* | 4 | 0.1169 | | –0.00958 | 0.0002463 |  | 49 | 0.091 | –3.05 | 51.9 |
|  |  |  | All Species | | | | 749 | 0.065 | –4.98 | 66.0 |

**^†^**$\max k_{{deacc}_{T}}= \beta_{1} \times T+ \beta_{2} \times T^{2}+ \beta_{3} \times T^{3}$, where T is the temperature in °C.

**^‡^**$\Psi_{deacc}=d+ \frac{(1-d)}{1+ e^{[b\times(\ln\left( chill \right)-\ln\left( c \right)]}}$, where *chill* is the accumulated chill in portions.
