## Supplementary material for "Woody species do not differ in dormancy progression: differences in time to budbreak due to forcing and cold hardiness": Table S4

**Supporting Information Table S3. Estimates, fitness statistics, and displays associated with linear regressions for each collection and species.**

| Species | Collection | n | α ± error | | β ± error | | *r^2^* | *P*-value |
| --- | --- | --- | --- | --- | --- | --- | --- | --- |
| *Abies balsamea* | AA20 | 45 | -15.26 | ±1.07 | 0.00 | ±0.16 | 0.00 | 0.975 |
|  | AB20 | 50 | -14.47 | ±1.15 | -0.33 | ±0.18 | 0.07 | 0.071 |
|  | AC20 | 40 | -15.10 | ±1.12 | -0.15 | ±0.22 | 0.01 | 0.480 |
|  | AD20 | 41 | -18.13 | ±0.98 | 0.08 | ±0.16 | 0.01 | 0.591 |
|  | AE20 | 51 | -17.26 | ±0.71 | -0.33 | ±0.07 | 0.30 | <0.001 |
|  | AF20 | 61 | -18.31 | ±0.60 | -0.50 | ±0.11 | 0.26 | <0.001 |
|  | AG20 | 56 | -19.85 | ±0.52 | -0.15 | ±0.03 | 0.26 | <0.001 |
|  | AH20 | 41 | -22.91 | ±0.45 | 0.04 | ±0.03 | 0.06 | 0.132 |
|  | AI20 | 45 | -24.76 | ±0.35 | 0.01 | ±0.01 | 0.01 | 0.601 |
|  | AJ20 | 46 | -25.85 | ±0.48 | 0.00 | ±0.02 | 0.00 | 0.942 |
|  | AK20 | 43 | -25.83 | ±0.54 | 0.10 | ±0.03 | 0.26 | <0.001 |
|  | AL20 | 35 | -27.45 | ±0.76 | 0.13 | ±0.04 | 0.22 | 0.005 |
|  | AM20 | 33 | -29.35 | ±0.59 | 0.40 | ±0.03 | 0.82 | <0.001 |
|  | AN20 | 45 | -27.44 | ±0.64 | 0.35 | ±0.04 | 0.69 | <0.001 |
|  | AO20 | 41 | -29.57 | ±0.84 | 0.65 | ±0.06 | 0.77 | <0.001 |
|  | AP20 | 40 | -27.72 | ±0.63 | 0.38 | ±0.05 | 0.58 | <0.001 |
|  | AQ20 | 46 | -27.93 | ±0.52 | 0.68 | ±0.05 | 0.83 | <0.001 |
|  | AR20 | 47 | -27.88 | ±0.60 | 0.57 | ±0.05 | 0.75 | <0.001 |
|  | AS20 | 44 | -29.99 | ±0.59 | 0.98 | ±0.05 | 0.90 | <0.001 |
|  | AT20 | 48 | -27.42 | ±0.46 | 0.71 | ±0.05 | 0.82 | <0.001 |
|  | AU20 | 43 | -26.51 | ±0.46 | 0.98 | ±0.06 | 0.85 | <0.001 |
|  | AV20 | 26 | -26.83 | ±0.46 | 0.92 | ±0.07 | 0.87 | <0.001 |
|  | AW20 | 46 | -26.78 | ±0.53 | 1.08 | ±0.08 | 0.80 | <0.001 |
|  | AX20 | 42 | -26.42 | ±0.59 | 1.18 | ±0.11 | 0.73 | <0.001 |
|  | AY20 | 38 | -25.84 | ±0.66 | 0.98 | ±0.12 | 0.66 | <0.001 |
|  | AZ20 | 33 | -25.77 | ±0.42 | 1.11 | ±0.08 | 0.86 | <0.001 |
|  | BA20 | 45 | -22.87 | ±0.39 | 0.97 | ±0.11 | 0.63 | <0.001 |
|  | AA21 | 25 | -14.47 | ±0.72 | -0.26 | ±0.13 | 0.15 | 0.059 |
|  | AB21 | 38 | -15.16 | ±0.87 | -0.33 | ±0.11 | 0.20 | 0.006 |
|  | AC21 | 49 | -20.89 | ±0.68 | -0.08 | ±0.05 | 0.05 | 0.135 |
|  | AD21 | 55 | -22.19 | ±0.46 | -0.11 | ±0.03 | 0.18 | 0.001 |
|  | AE21 | 44 | -23.23 | ±0.45 | 0.09 | ±0.03 | 0.22 | 0.001 |
|  | AF21 | 41 | -24.73 | ±0.35 | 0.01 | ±0.03 | 0.01 | 0.638 |
|  | AG21 | 40 | -23.69 | ±0.38 | -0.06 | ±0.03 | 0.10 | 0.047 |
|  | AH21 | 49 | -26.45 | ±0.33 | 0.11 | ±0.02 | 0.35 | <0.001 |
|  | AI21 | 40 | -23.40 | ±0.59 | -0.12 | ±0.06 | 0.09 | 0.058 |
|  | AJ21 | 61 | -25.62 | ±0.34 | 0.04 | ±0.02 | 0.10 | 0.013 |
|  | AK21 | 53 | -26.76 | ±0.30 | 0.12 | ±0.01 | 0.56 | <0.001 |
|  | AL21 | 59 | -25.54 | ±0.30 | 0.08 | ±0.01 | 0.37 | <0.001 |
|  | AM21 | 58 | -27.08 | ±0.43 | 0.08 | ±0.02 | 0.16 | 0.002 |
|  | AN21 | 56 | -29.59 | ±0.28 | 0.18 | ±0.02 | 0.68 | <0.001 |
|  | AO21 | 47 | -29.07 | ±0.52 | 0.40 | ±0.04 | 0.66 | <0.001 |
|  | AP21 | 45 | -29.74 | ±0.75 | 0.55 | ±0.06 | 0.70 | <0.001 |
|  | AQ21 | 39 | -27.83 | ±0.40 | 0.28 | ±0.03 | 0.63 | <0.001 |
|  | AR21 | 45 | -28.00 | ±0.51 | 0.54 | ±0.04 | 0.78 | <0.001 |
|  | AS21 | 43 | -27.18 | ±0.50 | 0.42 | ±0.04 | 0.72 | <0.001 |
|  | AT21 | 43 | -30.21 | ±0.32 | 0.77 | ±0.04 | 0.89 | <0.001 |
|  | AU21 | 55 | -27.27 | ±0.57 | 0.72 | ±0.06 | 0.74 | <0.001 |
|  | AV21 | 45 | -26.84 | ±0.54 | 0.69 | ±0.08 | 0.60 | <0.001 |
|  | AW21 | 47 | -27.98 | ±0.70 | 0.97 | ±0.14 | 0.50 | <0.001 |
|  | AX21 | 48 | -26.16 | ±0.56 | 1.24 | ±0.22 | 0.40 | <0.001 |
| *Acer rubrum* | AA20 | 26 | -8.68 | ±1.18 | 0.02 | ±0.15 | 0.00 | 0.885 |
|  | AB20 | 27 | -7.06 | ±0.77 | -0.05 | ±0.12 | 0.01 | 0.716 |
|  | AC20 | 34 | -9.16 | ±1.73 | -0.01 | ±0.29 | 0.00 | 0.982 |
|  | AD20 | 29 | -9.99 | ±1.29 | 0.10 | ±0.21 | 0.01 | 0.634 |
|  | AE20 | 35 | -9.75 | ±1.02 | -0.30 | ±0.12 | 0.16 | 0.017 |
|  | AF20 | 39 | -18.84 | ±0.87 | 0.08 | ±0.11 | 0.02 | 0.455 |
|  | AG20 | 33 | -20.25 | ±0.53 | 0.09 | ±0.04 | 0.16 | 0.019 |
|  | AH20 | 25 | -21.82 | ±0.77 | 0.06 | ±0.05 | 0.06 | 0.237 |
|  | AI20 | 31 | -21.58 | ±0.50 | 0.00 | ±0.02 | 0.00 | 0.883 |
|  | AJ20 | 37 | -24.89 | ±0.89 | 0.12 | ±0.04 | 0.19 | 0.008 |
|  | AK20 | 44 | -21.99 | ±0.59 | -0.01 | ±0.03 | 0.00 | 0.686 |
|  | AL20 | 33 | -25.09 | ±0.69 | 0.10 | ±0.04 | 0.19 | 0.012 |
|  | AM20 | 26 | -24.06 | ±0.92 | 0.09 | ±0.06 | 0.09 | 0.144 |
|  | AN20 | 30 | -23.74 | ±1.34 | 0.16 | ±0.07 | 0.15 | 0.036 |
|  | AO20 | 32 | -26.51 | ±0.48 | 0.24 | ±0.04 | 0.58 | <0.001 |
|  | AP20 | 34 | -25.36 | ±1.06 | 0.28 | ±0.09 | 0.24 | 0.004 |
|  | AQ20 | 35 | -25.94 | ±1.22 | 0.89 | ±0.14 | 0.55 | <0.001 |
|  | AR20 | 42 | -23.50 | ±0.94 | 0.38 | ±0.08 | 0.36 | <0.001 |
|  | AS20 | 45 | -23.62 | ±0.78 | 0.44 | ±0.07 | 0.46 | <0.001 |
|  | AT20 | 36 | -25.52 | ±0.91 | 0.84 | ±0.13 | 0.55 | <0.001 |
|  | AU20 | 30 | -23.01 | ±0.85 | 1.65 | ±0.41 | 0.37 | <0.001 |
|  | AV20 | 47 | -25.38 | ±0.57 | 2.54 | ±0.23 | 0.74 | <0.001 |
|  | AW20 | 35 | -24.62 | ±0.59 | 2.54 | ±0.31 | 0.67 | <0.001 |
|  | AX20 | 40 | -21.99 | ±0.83 | 2.48 | ±0.31 | 0.63 | <0.001 |
|  | AY20 | 30 | -20.41 | ±0.57 | 3.41 | ±0.44 | 0.68 | <0.001 |
|  | AZ20 | 30 | -19.37 | ±0.84 | 3.20 | ±0.65 | 0.46 | <0.001 |
|  | AA21 | 12 | -4.16 | ±1.19 | -0.32 | ±0.17 | 0.25 | 0.097 |
|  | AB21 | 28 | -2.99 | ±1.68 | -0.65 | ±0.22 | 0.26 | 0.006 |
|  | AC21 | 47 | -9.08 | ±0.95 | -0.24 | ±0.08 | 0.17 | 0.004 |
|  | AD21 | 47 | -17.08 | ±0.50 | 0.11 | ±0.03 | 0.19 | 0.002 |
|  | AE21 | 45 | -19.04 | ±0.71 | 0.24 | ±0.04 | 0.46 | <0.001 |
|  | AF21 | 50 | -19.76 | ±0.64 | 0.18 | ±0.05 | 0.23 | <0.001 |
|  | AG21 | 60 | -20.13 | ±0.49 | 0.15 | ±0.02 | 0.43 | <0.001 |
|  | AH21 | 57 | -21.59 | ±0.52 | 0.15 | ±0.03 | 0.29 | <0.001 |
|  | AI21 | 69 | -21.36 | ±0.43 | 0.14 | ±0.02 | 0.49 | <0.001 |
|  | AJ21 | 70 | -22.60 | ±0.45 | 0.18 | ±0.02 | 0.53 | <0.001 |
|  | AK21 | 79 | -23.40 | ±0.49 | 0.15 | ±0.02 | 0.42 | <0.001 |
|  | AL21 | 80 | -23.83 | ±0.49 | 0.22 | ±0.02 | 0.54 | <0.001 |
|  | AM21 | 70 | -24.39 | ±0.44 | 0.19 | ±0.03 | 0.42 | <0.001 |
|  | AN21 | 70 | -24.71 | ±0.56 | 0.21 | ±0.04 | 0.32 | <0.001 |
|  | AO21 | 50 | -25.33 | ±0.67 | 0.25 | ±0.06 | 0.29 | <0.001 |
|  | AP21 | 59 | -26.40 | ±0.71 | 0.41 | ±0.05 | 0.52 | <0.001 |
|  | AQ21 | 50 | -24.78 | ±0.67 | 0.29 | ±0.05 | 0.39 | <0.001 |
|  | AR21 | 69 | -25.89 | ±0.68 | 0.30 | ±0.06 | 0.28 | <0.001 |
|  | AS21 | 58 | -25.94 | ±0.71 | 0.50 | ±0.09 | 0.34 | <0.001 |
|  | AT21 | 50 | -27.42 | ±0.72 | 0.76 | ±0.13 | 0.42 | <0.001 |
|  | AU21 | 58 | -28.03 | ±0.78 | 1.34 | ±0.11 | 0.74 | <0.001 |
|  | AV21 | 80 | -26.17 | ±0.63 | 1.59 | ±0.13 | 0.64 | <0.001 |
|  | AW21 | 79 | -26.23 | ±0.67 | 2.00 | ±0.15 | 0.69 | <0.001 |
|  | AX21 | 50 | -24.68 | ±0.70 | 2.25 | ±0.29 | 0.57 | <0.001 |
| *Acer saccharum* | AA20 | 24 | -6.56 | ±0.34 | 0.11 | ±0.05 | 0.18 | 0.039 |
|  | AB20 | 24 | -6.80 | ±0.41 | 0.20 | ±0.07 | 0.27 | 0.009 |
|  | AC20 | 20 | -5.80 | ±0.39 | 0.05 | ±0.08 | 0.02 | 0.526 |
|  | AD20 | 21 | -7.20 | ±0.97 | 0.16 | ±0.14 | 0.06 | 0.267 |
|  | AE20 | 24 | -5.56 | ±1.99 | -2.01 | ±0.39 | 0.55 | <0.001 |
|  | AF20 | 19 | -20.65 | ±1.44 | 0.32 | ±0.27 | 0.08 | 0.252 |
|  | AG20 | 27 | -24.26 | ±0.73 | 0.21 | ±0.05 | 0.43 | <0.001 |
|  | AH20 | 20 | -21.10 | ±0.51 | 0.04 | ±0.03 | 0.12 | 0.133 |
|  | AI20 | 22 | -26.19 | ±0.38 | 0.15 | ±0.02 | 0.84 | <0.001 |
|  | AJ20 | 23 | -26.43 | ±0.58 | 0.23 | ±0.03 | 0.76 | <0.001 |
|  | AK20 | 25 | -27.17 | ±0.84 | 0.22 | ±0.04 | 0.56 | <0.001 |
|  | AL20 | 18 | -28.16 | ±0.93 | 0.27 | ±0.05 | 0.64 | <0.001 |
|  | AM20 | 21 | -28.20 | ±0.71 | 0.26 | ±0.04 | 0.71 | <0.001 |
|  | AN20 | 23 | -29.14 | ±0.72 | 0.23 | ±0.04 | 0.62 | <0.001 |
|  | AO20 | 28 | -29.72 | ±0.66 | 0.30 | ±0.04 | 0.70 | <0.001 |
|  | AP20 | 23 | -30.28 | ±0.83 | 0.37 | ±0.07 | 0.56 | <0.001 |
|  | AQ20 | 23 | -31.49 | ±0.65 | 0.52 | ±0.06 | 0.78 | <0.001 |
|  | AR20 | 27 | -30.06 | ±1.31 | 0.42 | ±0.10 | 0.42 | <0.001 |
|  | AS20 | 32 | -29.77 | ±0.66 | 0.30 | ±0.06 | 0.46 | <0.001 |
|  | AT20 | 28 | -29.93 | ±0.70 | 0.24 | ±0.08 | 0.26 | 0.005 |
|  | AU20 | 32 | -30.34 | ±0.69 | 0.42 | ±0.09 | 0.42 | <0.001 |
|  | AV20 | 26 | -31.23 | ±0.44 | 0.48 | ±0.04 | 0.83 | <0.001 |
|  | AW20 | 35 | -30.32 | ±0.59 | 0.50 | ±0.11 | 0.40 | <0.001 |
|  | AX20 | 29 | -31.69 | ±0.54 | 0.92 | ±0.10 | 0.77 | <0.001 |
|  | AY20 | 23 | -29.54 | ±0.49 | 1.24 | ±0.20 | 0.65 | <0.001 |
|  | AZ20 | 37 | -29.16 | ±0.70 | 1.08 | ±0.11 | 0.74 | <0.001 |
|  | BA20 | 28 | -27.07 | ±0.63 | 1.39 | ±0.20 | 0.64 | <0.001 |
|  | AA21 | 19 | -16.68 | ±1.01 | -0.05 | ±0.14 | 0.01 | 0.714 |
|  | AB21 | 40 | -19.42 | ±0.61 | -0.05 | ±0.07 | 0.01 | 0.515 |
|  | AC21 | 30 | -19.06 | ±0.88 | 0.29 | ±0.13 | 0.15 | 0.033 |
|  | AD21 | 59 | -18.48 | ±0.47 | 0.20 | ±0.03 | 0.38 | <0.001 |
|  | AE21 | 50 | -19.50 | ±0.66 | -0.03 | ±0.04 | 0.01 | 0.453 |
|  | AF21 | 48 | -20.70 | ±0.51 | 0.14 | ±0.04 | 0.22 | 0.001 |
|  | AG21 | 50 | -20.97 | ±0.58 | 0.21 | ±0.04 | 0.34 | <0.001 |
|  | AH21 | 60 | -22.11 | ±0.37 | 0.06 | ±0.02 | 0.12 | 0.006 |
|  | AI21 | 70 | -24.79 | ±0.46 | 0.20 | ±0.02 | 0.64 | <0.001 |
|  | AJ21 | 65 | -24.40 | ±0.48 | 0.15 | ±0.03 | 0.35 | <0.001 |
|  | AK21 | 70 | -23.76 | ±0.58 | 0.13 | ±0.02 | 0.33 | <0.001 |
|  | AL21 | 70 | -26.05 | ±0.52 | 0.20 | ±0.02 | 0.52 | <0.001 |
|  | AM21 | 60 | -25.84 | ±0.50 | 0.23 | ±0.03 | 0.52 | <0.001 |
|  | AN21 | 59 | -24.77 | ±0.50 | 0.17 | ±0.03 | 0.34 | <0.001 |
|  | AO21 | 50 | -24.99 | ±0.48 | 0.21 | ±0.04 | 0.36 | <0.001 |
|  | AP21 | 48 | -26.16 | ±0.56 | 0.22 | ±0.04 | 0.38 | <0.001 |
|  | AQ21 | 50 | -26.23 | ±0.58 | 0.22 | ±0.05 | 0.31 | <0.001 |
|  | AR21 | 49 | -27.23 | ±0.51 | 0.29 | ±0.06 | 0.35 | <0.001 |
|  | AS21 | 60 | -28.39 | ±0.54 | 0.47 | ±0.05 | 0.61 | <0.001 |
|  | AT21 | 60 | -27.26 | ±0.53 | 0.35 | ±0.05 | 0.41 | <0.001 |
|  | AU21 | 67 | -28.61 | ±0.56 | 0.63 | ±0.06 | 0.65 | <0.001 |
|  | AV21 | 59 | -28.97 | ±0.50 | 0.86 | ±0.07 | 0.71 | <0.001 |
|  | AW21 | 60 | -27.24 | ±0.51 | 0.79 | ±0.10 | 0.50 | <0.001 |
|  | AX21 | 50 | -26.20 | ±0.48 | 0.79 | ±0.20 | 0.25 | <0.001 |
| *Cercis canadensis* | AA20 | 29 | -7.88 | ±1.07 | -0.15 | ±0.13 | 0.05 | 0.255 |
|  | AB20 | 38 | -10.38 | ±0.88 | 0.24 | ±0.14 | 0.08 | 0.095 |
|  | AC20 | 29 | -10.48 | ±0.87 | 0.30 | ±0.15 | 0.13 | 0.059 |
|  | AD20 | 30 | -9.61 | ±0.71 | 0.00 | ±0.11 | 0.00 | 0.968 |
|  | AE20 | 35 | -10.14 | ±0.68 | 0.02 | ±0.08 | 0.00 | 0.812 |
|  | AF20 | 43 | -13.42 | ±0.67 | 0.16 | ±0.08 | 0.09 | 0.045 |
|  | AG20 | 39 | -13.66 | ±0.58 | 0.13 | ±0.04 | 0.26 | 0.001 |
|  | AH20 | 33 | -11.71 | ±0.44 | 0.05 | ±0.02 | 0.13 | 0.038 |
|  | AI20 | 25 | -12.98 | ±0.54 | 0.08 | ±0.02 | 0.39 | 0.001 |
|  | AJ20 | 41 | -14.49 | ±0.62 | 0.14 | ±0.03 | 0.29 | <0.001 |
|  | AK20 | 32 | -15.26 | ±0.64 | 0.18 | ±0.03 | 0.57 | <0.001 |
|  | AL20 | 27 | -15.36 | ±0.58 | 0.25 | ±0.03 | 0.68 | <0.001 |
|  | AM20 | 28 | -16.03 | ±0.56 | 0.34 | ±0.03 | 0.83 | <0.001 |
|  | AN20 | 34 | -16.05 | ±0.91 | 0.38 | ±0.05 | 0.66 | <0.001 |
|  | AO20 | 38 | -17.04 | ±0.63 | 0.40 | ±0.04 | 0.74 | <0.001 |
|  | AP20 | 36 | -17.65 | ±0.66 | 0.56 | ±0.06 | 0.73 | <0.001 |
|  | AQ20 | 40 | -18.50 | ±0.57 | 0.67 | ±0.05 | 0.80 | <0.001 |
|  | AR20 | 30 | -17.55 | ±0.62 | 0.49 | ±0.05 | 0.77 | <0.001 |
|  | AS20 | 20 | -17.62 | ±0.95 | 1.09 | ±0.14 | 0.78 | <0.001 |
|  | AT20 | 26 | -17.42 | ±0.77 | 0.95 | ±0.16 | 0.59 | <0.001 |
|  | AU20 | 19 | -16.49 | ±0.78 | 1.25 | ±0.17 | 0.76 | <0.001 |
|  | AV20 | 40 | -14.41 | ±0.68 | 1.53 | ±0.27 | 0.46 | <0.001 |
|  | AW20 | 38 | -13.03 | ±0.53 | 1.92 | ±0.22 | 0.68 | <0.001 |
|  | AX20 | 44 | -12.13 | ±0.55 | 1.52 | ±0.23 | 0.51 | <0.001 |
|  | AY20 | 41 | -11.36 | ±0.42 | 1.39 | ±0.17 | 0.63 | <0.001 |
|  | AZ20 | 24 | -9.39 | ±0.72 | 1.38 | ±0.55 | 0.22 | 0.021 |
|  | BA20 | 27 | -8.51 | ±0.56 | 1.47 | ±0.45 | 0.29 | 0.003 |
|  | AA21 | 25 | -6.71 | ±0.50 | -0.02 | ±0.08 | 0.00 | 0.780 |
|  | AB21 | 35 | -6.84 | ±0.63 | -0.19 | ±0.08 | 0.15 | 0.022 |
|  | AC21 | 51 | -8.89 | ±0.40 | -0.05 | ±0.03 | 0.05 | 0.115 |
|  | AD21 | 54 | -10.14 | ±0.35 | -0.01 | ±0.02 | 0.00 | 0.663 |
|  | AE21 | 43 | -11.35 | ±0.46 | 0.13 | ±0.03 | 0.36 | <0.001 |
|  | AF21 | 50 | -13.54 | ±0.47 | 0.17 | ±0.04 | 0.33 | <0.001 |
|  | AG21 | 59 | -12.57 | ±0.37 | 0.07 | ±0.02 | 0.20 | <0.001 |
|  | AH21 | 58 | -13.71 | ±0.50 | 0.08 | ±0.03 | 0.14 | 0.004 |
|  | AI21 | 70 | -15.33 | ±0.39 | 0.12 | ±0.02 | 0.48 | <0.001 |
|  | AJ21 | 70 | -14.17 | ±0.36 | 0.15 | ±0.02 | 0.53 | <0.001 |
|  | AK21 | 80 | -16.55 | ±0.38 | 0.20 | ±0.02 | 0.53 | <0.001 |
|  | AL21 | 78 | -15.78 | ±0.43 | 0.20 | ±0.02 | 0.55 | <0.001 |
|  | AM21 | 50 | -17.19 | ±0.49 | 0.34 | ±0.04 | 0.60 | <0.001 |
|  | AN21 | 50 | -16.12 | ±0.45 | 0.36 | ±0.03 | 0.71 | <0.001 |
|  | AO21 | 60 | -18.15 | ±0.49 | 0.44 | ±0.04 | 0.72 | <0.001 |
|  | AP21 | 49 | -17.64 | ±0.54 | 0.46 | ±0.06 | 0.55 | <0.001 |
|  | AQ21 | 70 | -16.20 | ±0.42 | 0.39 | ±0.05 | 0.51 | <0.001 |
|  | AR21 | 79 | -17.59 | ±0.36 | 0.49 | ±0.04 | 0.71 | <0.001 |
|  | AS21 | 70 | -18.46 | ±0.55 | 0.70 | ±0.08 | 0.56 | <0.001 |
|  | AT21 | 70 | -17.35 | ±0.55 | 0.64 | ±0.08 | 0.50 | <0.001 |
|  | AU21 | 70 | -17.35 | ±0.53 | 1.01 | ±0.09 | 0.63 | <0.001 |
|  | AV21 | 80 | -17.16 | ±0.56 | 1.25 | ±0.12 | 0.58 | <0.001 |
|  | AW21 | 70 | -16.84 | ±0.57 | 1.71 | ±0.16 | 0.63 | <0.001 |
| *Cornus florida* | AA20 | 19 | -6.54 | ±0.29 | 0.11 | ±0.04 | 0.32 | 0.011 |
|  | AB20 | 21 | -6.31 | ±0.29 | 0.12 | ±0.05 | 0.21 | 0.038 |
|  | AC20 | 17 | -5.92 | ±0.32 | 0.07 | ±0.05 | 0.09 | 0.234 |
|  | AD20 | 16 | -5.45 | ±0.31 | -0.02 | ±0.04 | 0.01 | 0.682 |
|  | AE20 | 30 | -9.41 | ±1.20 | -0.07 | ±0.10 | 0.02 | 0.442 |
|  | AF20 | 43 | -8.73 | ±1.66 | -0.80 | ±0.24 | 0.22 | 0.002 |
|  | AG20 | 52 | -22.72 | ±0.62 | 0.35 | ±0.04 | 0.59 | <0.001 |
|  | AH20 | 47 | -23.21 | ±0.81 | 0.29 | ±0.05 | 0.45 | <0.001 |
|  | AI20 | 42 | -22.36 | ±0.96 | 0.14 | ±0.03 | 0.29 | <0.001 |
|  | AJ20 | 48 | -24.04 | ±0.51 | 0.19 | ±0.03 | 0.54 | <0.001 |
|  | AK20 | 46 | -24.86 | ±0.92 | 0.31 | ±0.04 | 0.55 | <0.001 |
|  | AL20 | 36 | -24.20 | ±0.77 | 0.13 | ±0.04 | 0.21 | 0.005 |
|  | AM20 | 36 | -24.20 | ±0.32 | 0.15 | ±0.02 | 0.66 | <0.001 |
|  | AN20 | 37 | -24.72 | ±0.82 | 0.26 | ±0.06 | 0.39 | <0.001 |
|  | AO20 | 44 | -25.25 | ±0.72 | 0.22 | ±0.05 | 0.35 | <0.001 |
|  | AP20 | 49 | -23.98 | ±0.50 | 0.22 | ±0.04 | 0.34 | <0.001 |
|  | AQ20 | 48 | -25.03 | ±0.64 | 0.36 | ±0.06 | 0.44 | <0.001 |
|  | AR20 | 42 | -26.17 | ±0.86 | 0.82 | ±0.08 | 0.70 | <0.001 |
|  | AS20 | 36 | -27.43 | ±0.86 | 0.96 | ±0.10 | 0.75 | <0.001 |
|  | AT20 | 49 | -26.70 | ±0.95 | 0.96 | ±0.10 | 0.65 | <0.001 |
|  | AU20 | 47 | -23.25 | ±0.77 | 0.31 | ±0.11 | 0.14 | 0.010 |
|  | AW20 | 48 | -25.17 | ±0.68 | 1.13 | ±0.10 | 0.72 | <0.001 |
|  | AX20 | 54 | -25.85 | ±0.69 | 1.27 | ±0.12 | 0.69 | <0.001 |
|  | AY20 | 40 | -23.11 | ±0.76 | 1.32 | ±0.14 | 0.70 | <0.001 |
|  | AZ20 | 40 | -23.27 | ±0.94 | 1.33 | ±0.19 | 0.55 | <0.001 |
|  | AA21 | 10 | -4.59 | ±0.73 | 0.01 | ±0.10 | 0.00 | 0.956 |
|  | AB21 | 34 | -10.13 | ±0.89 | 0.25 | ±0.13 | 0.11 | 0.061 |
|  | AC21 | 49 | -10.05 | ±0.76 | -0.03 | ±0.07 | 0.00 | 0.630 |
|  | AD21 | 55 | -15.88 | ±0.71 | 0.23 | ±0.05 | 0.25 | <0.001 |
|  | AE21 | 50 | -17.49 | ±0.90 | 0.09 | ±0.05 | 0.05 | 0.112 |
|  | AF21 | 49 | -19.24 | ±0.81 | 0.05 | ±0.06 | 0.01 | 0.429 |
|  | AG21 | 56 | -19.82 | ±0.78 | 0.14 | ±0.04 | 0.17 | 0.002 |
|  | AH21 | 50 | -24.10 | ±0.54 | 0.28 | ±0.05 | 0.39 | <0.001 |
|  | AI21 | 60 | -24.45 | ±0.51 | 0.26 | ±0.03 | 0.59 | <0.001 |
|  | AJ21 | 60 | -24.37 | ±0.43 | 0.18 | ±0.03 | 0.42 | <0.001 |
|  | AK21 | 60 | -23.49 | ±0.52 | 0.13 | ±0.03 | 0.28 | <0.001 |
|  | AL21 | 59 | -23.23 | ±0.53 | 0.11 | ±0.03 | 0.18 | 0.001 |
|  | AM21 | 60 | -23.93 | ±0.35 | 0.10 | ±0.02 | 0.30 | <0.001 |
|  | AN21 | 60 | -23.30 | ±0.38 | 0.06 | ±0.02 | 0.11 | 0.008 |
|  | AO21 | 50 | -22.17 | ±0.54 | -0.05 | ±0.05 | 0.02 | 0.315 |
|  | AP21 | 50 | -24.74 | ±0.38 | 0.17 | ±0.03 | 0.44 | <0.001 |
|  | AQ21 | 50 | -23.68 | ±0.53 | 0.14 | ±0.04 | 0.17 | 0.003 |
|  | AR21 | 69 | -23.79 | ±0.56 | 0.35 | ±0.05 | 0.43 | <0.001 |
|  | AS21 | 49 | -24.14 | ±0.63 | 0.29 | ±0.08 | 0.23 | <0.001 |
|  | AT21 | 60 | -24.25 | ±0.72 | 0.66 | ±0.07 | 0.58 | <0.001 |
|  | AU21 | 59 | -24.93 | ±0.67 | 0.59 | ±0.09 | 0.42 | <0.001 |
|  | AV21 | 58 | -24.18 | ±0.57 | 0.60 | ±0.08 | 0.47 | <0.001 |
|  | AW21 | 60 | -24.20 | ±0.57 | 0.40 | ±0.09 | 0.24 | <0.001 |
|  | AX21 | 50 | -24.64 | ±0.42 | 0.80 | ±0.17 | 0.30 | <0.001 |
| *Cornus mas* | AA20 | 20 | -6.09 | ±0.39 | 0.11 | ±0.05 | 0.19 | 0.058 |
|  | AB20 | 19 | -5.65 | ±0.25 | 0.04 | ±0.05 | 0.03 | 0.460 |
|  | AC20 | 11 | -5.17 | ±0.36 | 0.08 | ±0.07 | 0.13 | 0.270 |
|  | AD20 | 14 | -5.23 | ±0.26 | 0.07 | ±0.04 | 0.18 | 0.132 |
|  | AE20 | 22 | -3.69 | ±0.67 | -0.58 | ±0.05 | 0.86 | <0.001 |
|  | AF20 | 57 | -10.46 | ±0.93 | -0.53 | ±0.09 | 0.37 | <0.001 |
|  | AG20 | 59 | -18.20 | ±0.52 | 0.07 | ±0.03 | 0.08 | 0.033 |
|  | AH20 | 49 | -18.68 | ±0.45 | 0.06 | ±0.03 | 0.11 | 0.020 |
|  | AI20 | 48 | -20.81 | ±0.69 | 0.18 | ±0.03 | 0.51 | <0.001 |
|  | AJ20 | 50 | -20.61 | ±0.68 | 0.19 | ±0.03 | 0.39 | <0.001 |
|  | AK20 | 48 | -21.57 | ±0.79 | 0.31 | ±0.04 | 0.58 | <0.001 |
|  | AL20 | 40 | -21.30 | ±0.94 | 0.39 | ±0.05 | 0.58 | <0.001 |
|  | AM20 | 30 | -20.67 | ±0.84 | 0.42 | ±0.08 | 0.48 | <0.001 |
|  | AN20 | 40 | -20.89 | ±0.96 | 0.53 | ±0.07 | 0.59 | <0.001 |
|  | AO20 | 27 | -23.02 | ±1.09 | 1.28 | ±0.15 | 0.73 | <0.001 |
|  | AP20 | 36 | -21.71 | ±1.16 | 1.00 | ±0.17 | 0.49 | <0.001 |
|  | AQ20 | 39 | -22.55 | ±0.94 | 1.41 | ±0.16 | 0.69 | <0.001 |
|  | AR20 | 68 | -20.97 | ±0.73 | 1.00 | ±0.15 | 0.41 | <0.001 |
|  | AS20 | 39 | -21.49 | ±1.01 | 1.76 | ±0.40 | 0.34 | <0.001 |
|  | AT20 | 40 | -20.76 | ±1.00 | 1.22 | ±0.26 | 0.37 | <0.001 |
|  | AU20 | 40 | -20.07 | ±0.66 | 1.22 | ±0.35 | 0.24 | 0.001 |
|  | AV20 | 38 | -22.27 | ±0.90 | 3.56 | ±0.50 | 0.58 | <0.001 |
|  | AW20 | 30 | -22.09 | ±0.80 | 3.26 | ±0.62 | 0.49 | <0.001 |
|  | AX20 | 39 | -21.58 | ±0.99 | 4.03 | ±0.53 | 0.61 | <0.001 |
|  | AY20 | 30 | -22.22 | ±0.67 | 3.59 | ±0.52 | 0.63 | <0.001 |
|  | AZ20 | 20 | -20.19 | ±1.02 | 5.36 | ±1.44 | 0.43 | 0.002 |
|  | AA21 | 32 | -8.30 | ±0.94 | -0.54 | ±0.14 | 0.34 | <0.001 |
|  | AB21 | 44 | -13.07 | ±1.06 | -0.24 | ±0.13 | 0.08 | 0.063 |
|  | AC21 | 60 | -12.53 | ±0.80 | -0.12 | ±0.07 | 0.05 | 0.083 |
|  | AD21 | 54 | -12.92 | ±0.81 | -0.17 | ±0.06 | 0.16 | 0.003 |
|  | AE21 | 45 | -16.54 | ±0.70 | 0.04 | ±0.04 | 0.03 | 0.271 |
|  | AF21 | 47 | -19.26 | ±0.70 | 0.10 | ±0.05 | 0.07 | 0.063 |
|  | AG21 | 60 | -19.50 | ±0.51 | 0.12 | ±0.02 | 0.30 | <0.001 |
|  | AH21 | 60 | -19.08 | ±0.46 | 0.09 | ±0.03 | 0.18 | 0.001 |
|  | AI21 | 70 | -19.52 | ±0.50 | 0.10 | ±0.03 | 0.14 | 0.001 |
|  | AJ21 | 69 | -20.44 | ±0.47 | 0.12 | ±0.03 | 0.18 | <0.001 |
|  | AK21 | 79 | -20.39 | ±0.48 | 0.22 | ±0.03 | 0.45 | <0.001 |
|  | AL21 | 80 | -19.84 | ±0.54 | 0.18 | ±0.03 | 0.26 | <0.001 |
|  | AM21 | 60 | -20.33 | ±0.61 | 0.28 | ±0.05 | 0.32 | <0.001 |
|  | AN21 | 49 | -20.87 | ±0.58 | 0.31 | ±0.07 | 0.31 | <0.001 |
|  | AO21 | 40 | -20.92 | ±0.97 | 0.68 | ±0.13 | 0.41 | <0.001 |
|  | AP21 | 69 | -21.21 | ±0.75 | 0.63 | ±0.10 | 0.36 | <0.001 |
|  | AQ21 | 78 | -21.00 | ±0.59 | 0.73 | ±0.11 | 0.37 | <0.001 |
|  | AR21 | 79 | -20.12 | ±0.68 | 0.82 | ±0.15 | 0.28 | <0.001 |
|  | AS21 | 60 | -20.89 | ±0.75 | 1.12 | ±0.19 | 0.37 | <0.001 |
|  | AT21 | 70 | -21.80 | ±0.61 | 1.27 | ±0.17 | 0.45 | <0.001 |
|  | AU21 | 49 | -21.21 | ±0.62 | 1.08 | ±0.26 | 0.27 | <0.001 |
|  | AV21 | 70 | -20.55 | ±0.84 | 1.60 | ±0.23 | 0.41 | <0.001 |
|  | AW21 | 49 | -24.00 | ±0.76 | 2.65 | ±0.31 | 0.60 | <0.001 |
|  | AX21 | 39 | -22.07 | ±0.72 | 1.72 | ±0.38 | 0.35 | <0.001 |
| *Fagus grandifolia* | AA20 | 35 | -7.30 | ±0.25 | 0.08 | ±0.03 | 0.14 | 0.026 |
|  | AB20 | 33 | -8.18 | ±0.40 | 0.31 | ±0.07 | 0.42 | <0.001 |
|  | AC20 | 31 | -9.59 | ±0.68 | 0.44 | ±0.12 | 0.33 | 0.001 |
|  | AD20 | 32 | -7.91 | ±0.36 | 0.10 | ±0.05 | 0.10 | 0.081 |
|  | AE20 | 33 | -8.33 | ±0.36 | -0.12 | ±0.04 | 0.26 | 0.002 |
|  | AF20 | 47 | -10.53 | ±0.27 | -0.05 | ±0.03 | 0.06 | 0.094 |
|  | AG20 | 31 | -12.85 | ±0.32 | 0.05 | ±0.02 | 0.16 | 0.028 |
|  | AH20 | 23 | -13.43 | ±0.43 | -0.01 | ±0.03 | 0.01 | 0.727 |
|  | AI20 | 27 | -13.83 | ±0.47 | -0.01 | ±0.02 | 0.01 | 0.696 |
|  | AJ20 | 22 | -14.62 | ±0.49 | 0.03 | ±0.03 | 0.05 | 0.335 |
|  | AK20 | 26 | -14.66 | ±0.29 | 0.01 | ±0.01 | 0.04 | 0.355 |
|  | AL20 | 21 | -14.81 | ±0.67 | 0.03 | ±0.04 | 0.04 | 0.374 |
|  | AM20 | 21 | -18.71 | ±0.91 | 0.25 | ±0.05 | 0.54 | <0.001 |
|  | AN20 | 25 | -15.56 | ±0.72 | 0.02 | ±0.04 | 0.01 | 0.633 |
|  | AO20 | 28 | -16.87 | ±0.74 | 0.18 | ±0.05 | 0.36 | 0.001 |
|  | AP20 | 29 | -16.71 | ±0.43 | 0.14 | ±0.04 | 0.30 | 0.002 |
|  | AQ20 | 27 | -16.02 | ±0.27 | 0.11 | ±0.02 | 0.46 | <0.001 |
|  | AR20 | 32 | -16.02 | ±0.43 | 0.08 | ±0.04 | 0.13 | 0.045 |
|  | AS20 | 38 | -16.87 | ±0.51 | 0.18 | ±0.05 | 0.30 | <0.001 |
|  | AT20 | 38 | -18.19 | ±0.44 | 0.33 | ±0.05 | 0.58 | <0.001 |
|  | AU20 | 49 | -18.11 | ±0.64 | 0.42 | ±0.08 | 0.37 | <0.001 |
|  | AV20 | 22 | -17.23 | ±0.73 | 0.57 | ±0.29 | 0.16 | 0.063 |
|  | AW20 | 59 | -18.01 | ±0.65 | 0.48 | ±0.06 | 0.52 | <0.001 |
|  | AX20 | 31 | -16.02 | ±0.57 | 0.23 | ±0.15 | 0.08 | 0.120 |
|  | AY20 | 34 | -17.17 | ±0.52 | 0.80 | ±0.15 | 0.45 | <0.001 |
|  | AZ20 | 35 | -14.78 | ±0.43 | 0.46 | ±0.09 | 0.46 | <0.001 |
|  | BA20 | 47 | -13.40 | ±0.52 | 0.73 | ±0.16 | 0.33 | <0.001 |
|  | AA21 | 32 | -5.00 | ±0.67 | -0.07 | ±0.10 | 0.02 | 0.482 |
|  | AB21 | 34 | -8.08 | ±0.45 | 0.06 | ±0.06 | 0.03 | 0.322 |
|  | AC21 | 53 | -9.87 | ±0.30 | 0.16 | ±0.03 | 0.40 | <0.001 |
|  | AD21 | 57 | -9.83 | ±0.29 | 0.00 | ±0.02 | 0.00 | 0.816 |
|  | AE21 | 50 | -10.60 | ±0.51 | -0.02 | ±0.03 | 0.01 | 0.534 |
|  | AF21 | 46 | -11.71 | ±0.34 | -0.03 | ±0.03 | 0.03 | 0.213 |
|  | AG21 | 44 | -12.57 | ±0.36 | -0.05 | ±0.03 | 0.09 | 0.054 |
|  | AH21 | 43 | -14.03 | ±0.32 | 0.01 | ±0.03 | 0.00 | 0.659 |
|  | AI21 | 54 | -14.43 | ±0.27 | 0.02 | ±0.02 | 0.02 | 0.272 |
|  | AJ21 | 61 | -15.21 | ±0.41 | 0.07 | ±0.02 | 0.17 | 0.001 |
|  | AK21 | 56 | -15.18 | ±0.36 | 0.02 | ±0.02 | 0.03 | 0.203 |
|  | AL21 | 60 | -14.73 | ±0.40 | 0.01 | ±0.02 | 0.00 | 0.676 |
|  | AM21 | 51 | -15.38 | ±0.31 | 0.04 | ±0.02 | 0.10 | 0.025 |
|  | AN21 | 46 | -16.20 | ±0.39 | 0.06 | ±0.02 | 0.13 | 0.015 |
|  | AO21 | 38 | -17.00 | ±0.51 | 0.08 | ±0.04 | 0.09 | 0.067 |
|  | AP21 | 42 | -16.62 | ±0.42 | 0.21 | ±0.03 | 0.57 | <0.001 |
|  | AQ21 | 40 | -16.41 | ±0.46 | 0.17 | ±0.04 | 0.38 | <0.001 |
|  | AR21 | 43 | -18.32 | ±0.31 | 0.26 | ±0.02 | 0.74 | <0.001 |
|  | AS21 | 55 | -16.90 | ±0.51 | 0.23 | ±0.04 | 0.34 | <0.001 |
|  | AT21 | 53 | -17.08 | ±0.51 | 0.16 | ±0.05 | 0.16 | 0.003 |
|  | AU21 | 65 | -17.48 | ±0.47 | 0.32 | ±0.05 | 0.42 | <0.001 |
|  | AV21 | 52 | -17.37 | ±0.49 | 0.26 | ±0.07 | 0.22 | <0.001 |
|  | AW21 | 43 | -16.78 | ±0.39 | 0.33 | ±0.08 | 0.27 | <0.001 |
| *Forsythia* ‘Meadowlark’ | AA20 | 29 | -7.14 | ±0.57 | 0.13 | ±0.08 | 0.09 | 0.107 |
|  | AB20 | 27 | -6.06 | ±0.29 | 0.04 | ±0.05 | 0.03 | 0.381 |
|  | AC20 | 24 | -6.06 | ±0.39 | -0.02 | ±0.08 | 0.00 | 0.808 |
|  | AD20 | 23 | -7.80 | ±0.85 | 0.16 | ±0.13 | 0.06 | 0.245 |
|  | AE20 | 25 | -7.55 | ±0.93 | -0.14 | ±0.10 | 0.09 | 0.157 |
|  | AF20 | 42 | -15.69 | ±1.00 | -0.26 | ±0.13 | 0.08 | 0.065 |
|  | AG20 | 51 | -20.07 | ±0.51 | 0.01 | ±0.04 | 0.00 | 0.816 |
|  | AH20 | 40 | -20.82 | ±0.92 | 0.29 | ±0.05 | 0.43 | <0.001 |
|  | AI20 | 30 | -23.28 | ±0.81 | 0.40 | ±0.04 | 0.80 | <0.001 |
|  | AJ20 | 38 | -22.75 | ±0.53 | 0.18 | ±0.03 | 0.49 | <0.001 |
|  | AK20 | 28 | -23.86 | ±0.73 | 0.49 | ±0.04 | 0.86 | <0.001 |
|  | AL20 | 26 | -25.47 | ±1.28 | 0.66 | ±0.08 | 0.75 | <0.001 |
|  | AM20 | 21 | -24.63 | ±1.48 | 0.75 | ±0.15 | 0.56 | <0.001 |
|  | AN20 | 26 | -24.37 | ±1.42 | 0.83 | ±0.11 | 0.72 | <0.001 |
|  | AO20 | 22 | -25.56 | ±1.39 | 1.41 | ±0.20 | 0.72 | <0.001 |
|  | AP20 | 18 | -24.89 | ±1.37 | 1.83 | ±0.25 | 0.78 | <0.001 |
|  | AQ20 | 29 | -25.13 | ±1.06 | 2.29 | ±0.19 | 0.85 | <0.001 |
|  | AR20 | 42 | -24.92 | ±1.13 | 2.66 | ±0.31 | 0.64 | <0.001 |
|  | AS20 | 35 | -24.71 | ±1.09 | 3.94 | ±0.44 | 0.71 | <0.001 |
|  | AT20 | 15 | -22.93 | ±0.95 | 5.52 | ±0.43 | 0.93 | <0.001 |
|  | AU20 | 21 | -23.66 | ±0.72 | 8.04 | ±0.63 | 0.90 | <0.001 |
|  | AV20 | 24 | -24.83 | ±1.26 | 8.11 | ±0.97 | 0.76 | <0.001 |
|  | AW20 | 19 | -22.60 | ±0.71 | 7.83 | ±0.58 | 0.92 | <0.001 |
|  | AX20 | 30 | -19.81 | ±1.51 | 4.36 | ±0.86 | 0.48 | <0.001 |
|  | AY20 | 18 | -20.92 | ±1.56 | 12.49 | 2.21 | 0.67 | <0.001 |
|  | AZ20 | 12 | -15.10 | ±1.39 | 9.50 | ±1.96 | 0.70 | 0.001 |
|  | AA21 | 30 | -10.87 | ±1.82 | -0.03 | ±0.28 | 0.00 | 0.920 |
|  | AB21 | 44 | -15.00 | ±0.96 | -0.42 | ±0.12 | 0.21 | 0.002 |
|  | AC21 | 42 | -20.42 | ±0.81 | 0.75 | ±0.08 | 0.69 | <0.001 |
|  | AD21 | 43 | -16.91 | ±0.68 | 0.04 | ±0.06 | 0.01 | 0.510 |
|  | AE21 | 38 | -20.66 | ±0.75 | 0.19 | ±0.05 | 0.29 | <0.001 |
|  | AF21 | 45 | -19.85 | ±0.56 | 0.07 | ±0.04 | 0.05 | 0.126 |
|  | AG21 | 35 | -20.69 | ±1.31 | 0.42 | ±0.10 | 0.35 | <0.001 |
|  | AH21 | 40 | -22.14 | ±1.02 | 0.48 | ±0.09 | 0.41 | <0.001 |
|  | AI21 | 56 | -21.46 | ±0.50 | 0.17 | ±0.03 | 0.37 | <0.001 |
|  | AJ21 | 56 | -20.59 | ±0.41 | 0.08 | ±0.03 | 0.15 | 0.004 |
|  | AK21 | 64 | -23.24 | ±0.65 | 0.33 | ±0.04 | 0.57 | <0.001 |
|  | AL21 | 76 | -24.20 | ±0.54 | 0.35 | ±0.03 | 0.58 | <0.001 |
|  | AM21 | 50 | -21.60 | ±0.50 | 0.15 | ±0.05 | 0.16 | 0.004 |
|  | AN21 | 42 | -22.75 | ±0.47 | 0.15 | ±0.05 | 0.15 | 0.010 |
|  | AO21 | 41 | -23.91 | ±1.21 | 0.72 | ±0.13 | 0.45 | <0.001 |
|  | AP21 | 79 | -25.07 | ±0.66 | 0.48 | ±0.07 | 0.40 | <0.001 |
|  | AQ21 | 70 | -26.66 | ±0.91 | 1.66 | ±0.14 | 0.68 | <0.001 |
|  | AR21 | 58 | -26.04 | ±0.86 | 1.82 | ±0.18 | 0.64 | <0.001 |
|  | AS21 | 54 | -26.91 | ±1.00 | 1.82 | ±0.23 | 0.54 | <0.001 |
|  | AT21 | 57 | -25.77 | ±1.14 | 2.20 | ±0.27 | 0.55 | <0.001 |
|  | AU21 | 49 | -25.71 | ±1.41 | 2.92 | ±0.46 | 0.46 | <0.001 |
|  | AV21 | 46 | -26.52 | ±1.02 | 3.61 | ±0.38 | 0.67 | <0.001 |
|  | AW21 | 42 | -28.62 | ±1.23 | 5.51 | ±0.49 | 0.76 | <0.001 |
|  | AX21 | 40 | -24.29 | ±1.22 | 3.57 | ±0.48 | 0.59 | <0.001 |
| *Kalmia latifolia* | AA20 | 22 | -6.05 | ±0.35 | 0.01 | ±0.05 | 0.00 | 0.860 |
|  | AB20 | 14 | -5.93 | ±0.35 | -0.03 | ±0.06 | 0.02 | 0.607 |
|  | AC20 | 14 | -6.04 | ±0.28 | 0.08 | ±0.05 | 0.19 | 0.115 |
|  | AD20 | 18 | -6.70 | ±0.70 | 0.06 | ±0.11 | 0.02 | 0.594 |
|  | AE20 | 22 | -4.07 | ±0.73 | -0.64 | ±0.06 | 0.87 | <0.001 |
|  | AF20 | 28 | -5.31 | ±1.68 | -0.76 | ±0.13 | 0.55 | <0.001 |
|  | AG20 | 39 | -15.68 | ±1.69 | -0.19 | ±0.09 | 0.10 | 0.047 |
|  | AH20 | 49 | -20.50 | ±0.50 | 0.07 | ±0.03 | 0.12 | 0.016 |
|  | AI20 | 49 | -23.91 | ±0.49 | 0.05 | ±0.02 | 0.15 | 0.007 |
|  | AJ20 | 49 | -24.41 | ±0.57 | 0.17 | ±0.03 | 0.42 | <0.001 |
|  | AK20 | 48 | -22.76 | ±0.68 | 0.11 | ±0.03 | 0.20 | 0.001 |
|  | AL20 | 37 | -22.68 | ±0.93 | 0.28 | ±0.05 | 0.46 | <0.001 |
|  | AM20 | 39 | -23.35 | ±0.87 | 0.34 | ±0.05 | 0.54 | <0.001 |
|  | AN20 | 38 | -22.14 | ±0.93 | 0.21 | ±0.05 | 0.29 | <0.001 |
|  | AO20 | 47 | -24.71 | ±0.67 | 0.35 | ±0.04 | 0.58 | <0.001 |
|  | AP20 | 44 | -25.74 | ±0.98 | 0.54 | ±0.08 | 0.50 | <0.001 |
|  | AQ20 | 44 | -21.93 | ±0.97 | 0.43 | ±0.09 | 0.38 | <0.001 |
|  | AR20 | 45 | -22.36 | ±0.82 | 0.38 | ±0.07 | 0.41 | <0.001 |
|  | AS20 | 50 | -21.49 | ±0.89 | 0.29 | ±0.08 | 0.21 | 0.001 |
|  | AT20 | 48 | -21.11 | ±1.01 | 0.49 | ±0.11 | 0.30 | <0.001 |
|  | AU20 | 44 | -22.63 | ±0.82 | 0.48 | ±0.11 | 0.31 | <0.001 |
|  | AV20 | 30 | -21.51 | ±0.97 | 0.37 | ±0.16 | 0.17 | 0.024 |
|  | AW20 | 64 | -22.28 | ±0.66 | 0.54 | ±0.06 | 0.54 | <0.001 |
|  | AX20 | 49 | -23.09 | ±0.82 | 0.89 | ±0.12 | 0.55 | <0.001 |
|  | AY20 | 29 | -20.17 | ±1.06 | 0.93 | ±0.42 | 0.15 | 0.035 |
|  | AZ20 | 39 | -21.25 | ±0.96 | 0.71 | ±0.20 | 0.26 | 0.001 |
|  | BA20 | 40 | -18.95 | ±0.65 | 0.70 | ±0.20 | 0.24 | 0.001 |
|  | AA21 | 6 | -3.74 | ±0.61 | -0.08 | ±0.12 | 0.11 | 0.519 |
|  | AB21 | 15 | -5.00 | ±0.35 | 0.02 | ±0.07 | 0.01 | 0.738 |
|  | AC21 | 20 | -5.66 | ±0.69 | -0.03 | ±0.09 | 0.01 | 0.738 |
|  | AD21 | 26 | -3.54 | ±0.90 | -0.53 | ±0.06 | 0.79 | <0.001 |
|  | AE21 | 37 | -7.67 | ±1.08 | -0.33 | ±0.06 | 0.46 | <0.001 |
|  | AF21 | 47 | -13.04 | ±0.83 | -0.17 | ±0.06 | 0.14 | 0.009 |
|  | AG21 | 39 | -13.69 | ±0.97 | -0.27 | ±0.07 | 0.30 | <0.001 |
|  | AH21 | 47 | -20.61 | ±0.94 | 0.28 | ±0.09 | 0.19 | 0.002 |
|  | AI21 | 50 | -17.96 | ±0.75 | 0.00 | ±0.07 | 0.00 | 0.989 |
|  | AJ21 | 60 | -20.06 | ±0.64 | 0.02 | ±0.04 | 0.00 | 0.625 |
|  | AK21 | 66 | -21.77 | ±0.61 | 0.07 | ±0.02 | 0.12 | 0.004 |
|  | AL21 | 58 | -22.30 | ±0.56 | 0.17 | ±0.03 | 0.37 | <0.001 |
|  | AM21 | 52 | -23.75 | ±0.69 | 0.11 | ±0.04 | 0.15 | 0.005 |
|  | AN21 | 58 | -22.04 | ±0.74 | 0.04 | ±0.04 | 0.02 | 0.326 |
|  | AO21 | 50 | -23.68 | ±0.75 | 0.11 | ±0.06 | 0.06 | 0.088 |
|  | AP21 | 49 | -23.54 | ±0.75 | 0.15 | ±0.06 | 0.14 | 0.008 |
|  | AQ21 | 50 | -23.47 | ±0.88 | 0.37 | ±0.07 | 0.37 | <0.001 |
|  | AR21 | 59 | -22.47 | ±0.73 | 0.14 | ±0.04 | 0.15 | 0.003 |
|  | AS21 | 49 | -24.95 | ±0.77 | 0.56 | ±0.09 | 0.43 | <0.001 |
|  | AT21 | 49 | -23.90 | ±0.84 | 0.40 | ±0.11 | 0.23 | 0.001 |
|  | AU21 | 49 | -23.98 | ±0.81 | 0.73 | ±0.13 | 0.39 | <0.001 |
|  | AV21 | 59 | -23.51 | ±0.73 | 0.56 | ±0.11 | 0.32 | <0.001 |
|  | AW21 | 50 | -23.14 | ±0.77 | 0.86 | ±0.16 | 0.39 | <0.001 |
| *Larix kaempferi* | AA20 | 34 | -9.95 | ±0.98 | 0.00 | ±0.13 | 0.00 | 0.989 |
|  | AB20 | 30 | -11.69 | ±1.30 | 0.05 | ±0.21 | 0.00 | 0.798 |
|  | AC20 | 28 | -14.06 | ±1.68 | 0.29 | ±0.29 | 0.04 | 0.312 |
|  | AD20 | 26 | -12.28 | ±1.30 | -0.21 | ±0.18 | 0.05 | 0.252 |
|  | AE20 | 38 | -15.39 | ±0.98 | -0.20 | ±0.10 | 0.10 | 0.058 |
|  | AF20 | 43 | -21.77 | ±0.74 | 0.17 | ±0.08 | 0.09 | 0.046 |
|  | AG20 | 39 | -21.75 | ±0.44 | -0.07 | ±0.03 | 0.13 | 0.025 |
|  | AH20 | 32 | -22.48 | ±0.64 | 0.02 | ±0.04 | 0.01 | 0.588 |
|  | AI20 | 27 | -23.97 | ±0.45 | 0.02 | ±0.02 | 0.05 | 0.246 |
|  | AJ20 | 29 | -25.24 | ±0.45 | 0.09 | ±0.02 | 0.40 | <0.001 |
|  | AK20 | 29 | -24.19 | ±0.74 | 0.06 | ±0.04 | 0.09 | 0.108 |
|  | AL20 | 20 | -26.38 | ±0.87 | 0.36 | ±0.06 | 0.70 | <0.001 |
|  | AM20 | 23 | -25.54 | ±0.50 | 0.23 | ±0.03 | 0.78 | <0.001 |
|  | AN20 | 34 | -26.62 | ±1.16 | 0.37 | ±0.06 | 0.54 | <0.001 |
|  | AO20 | 27 | -24.95 | ±1.35 | 0.45 | ±0.11 | 0.39 | <0.001 |
|  | AP20 | 36 | -23.21 | ±0.95 | 0.30 | ±0.08 | 0.29 | 0.001 |
|  | AQ20 | 40 | -23.64 | ±0.79 | 0.53 | ±0.07 | 0.60 | <0.001 |
|  | AR20 | 41 | -22.70 | ±0.87 | 0.43 | ±0.07 | 0.47 | <0.001 |
|  | AS20 | 43 | -22.46 | ±0.91 | 0.46 | ±0.08 | 0.44 | <0.001 |
|  | AT20 | 43 | -22.03 | ±0.83 | 0.44 | ±0.09 | 0.37 | <0.001 |
|  | AU20 | 34 | -23.06 | ±0.97 | 0.81 | ±0.16 | 0.45 | <0.001 |
|  | AV20 | 30 | -24.41 | ±0.89 | 1.18 | ±0.25 | 0.45 | <0.001 |
|  | AW20 | 39 | -24.52 | ±0.78 | 1.09 | ±0.12 | 0.69 | <0.001 |
|  | AX20 | 51 | -23.57 | ±0.59 | 0.78 | ±0.09 | 0.61 | <0.001 |
|  | AY20 | 34 | -22.67 | ±0.63 | 1.11 | ±0.18 | 0.53 | <0.001 |
|  | AZ20 | 40 | -21.24 | ±0.54 | 0.78 | ±0.12 | 0.52 | <0.001 |
|  | AA21 | 10 | -4.37 | ±0.57 | -0.06 | ±0.09 | 0.05 | 0.554 |
|  | AB21 | 22 | -12.50 | ±1.66 | -0.32 | ±0.21 | 0.11 | 0.137 |
|  | AC21 | 48 | -17.68 | ±0.65 | -0.01 | ±0.06 | 0.00 | 0.922 |
|  | AD21 | 47 | -17.19 | ±0.68 | -0.18 | ±0.05 | 0.25 | <0.001 |
|  | AE21 | 41 | -19.34 | ±0.71 | 0.00 | ±0.04 | 0.00 | 0.912 |
|  | AF21 | 44 | -23.69 | ±0.64 | 0.17 | ±0.05 | 0.23 | 0.001 |
|  | AG21 | 50 | -20.63 | ±0.53 | 0.03 | ±0.03 | 0.03 | 0.262 |
|  | AH21 | 41 | -24.06 | ±0.49 | 0.12 | ±0.04 | 0.16 | 0.009 |
|  | AI21 | 62 | -23.98 | ±0.25 | 0.07 | ±0.01 | 0.43 | <0.001 |
|  | AJ21 | 57 | -24.09 | ±0.25 | 0.07 | ±0.01 | 0.37 | <0.001 |
|  | AK21 | 58 | -24.07 | ±0.34 | 0.08 | ±0.01 | 0.41 | <0.001 |
|  | AL21 | 55 | -24.70 | ±0.40 | 0.09 | ±0.02 | 0.37 | <0.001 |
|  | AM21 | 50 | -24.80 | ±0.26 | 0.16 | ±0.02 | 0.69 | <0.001 |
|  | AN21 | 49 | -23.70 | ±0.29 | 0.03 | ±0.02 | 0.05 | 0.115 |
|  | AO21 | 43 | -24.65 | ±0.43 | 0.08 | ±0.04 | 0.12 | 0.026 |
|  | AP21 | 40 | -24.92 | ±0.64 | 0.29 | ±0.05 | 0.51 | <0.001 |
|  | AQ21 | 46 | -25.47 | ±0.59 | 0.29 | ±0.05 | 0.47 | <0.001 |
|  | AR21 | 56 | -26.20 | ±0.64 | 0.33 | ±0.05 | 0.42 | <0.001 |
|  | AS21 | 58 | -26.25 | ±0.54 | 0.47 | ±0.05 | 0.63 | <0.001 |
|  | AT21 | 46 | -29.05 | ±0.75 | 0.92 | ±0.10 | 0.67 | <0.001 |
|  | AU21 | 56 | -25.39 | ±0.61 | 0.53 | ±0.06 | 0.59 | <0.001 |
|  | AV21 | 57 | -25.37 | ±0.58 | 0.68 | ±0.08 | 0.55 | <0.001 |
|  | AW21 | 55 | -29.08 | ±0.62 | 1.61 | ±0.12 | 0.77 | <0.001 |
| *Metasequoia glyptostroboides* | AA20 | 34 | -9.48 | ±0.72 | 0.05 | ±0.11 | 0.01 | 0.628 |
|  | AB20 | 30 | -8.21 | ±1.10 | -0.12 | ±0.19 | 0.01 | 0.552 |
|  | AC20 | 21 | -9.71 | ±0.88 | 0.27 | ±0.16 | 0.13 | 0.111 |
|  | AD20 | 26 | -9.21 | ±1.04 | 0.02 | ±0.16 | 0.00 | 0.897 |
|  | AE20 | 36 | -12.31 | ±1.03 | 0.18 | ±0.13 | 0.06 | 0.165 |
|  | AF20 | 58 | -15.47 | ±0.69 | 0.14 | ±0.08 | 0.06 | 0.073 |
|  | AG20 | 51 | -19.70 | ±0.49 | 0.31 | ±0.03 | 0.65 | <0.001 |
|  | AH20 | 36 | -20.54 | ±0.45 | 0.17 | ±0.03 | 0.54 | <0.001 |
|  | AI20 | 34 | -21.22 | ±0.58 | 0.15 | ±0.02 | 0.56 | <0.001 |
|  | AJ20 | 38 | -22.38 | ±0.63 | 0.18 | ±0.03 | 0.48 | <0.001 |
|  | AK20 | 31 | -22.91 | ±0.90 | 0.44 | ±0.05 | 0.75 | <0.001 |
|  | AL20 | 26 | -20.94 | ±1.04 | 0.22 | ±0.06 | 0.34 | 0.002 |
|  | AM20 | 25 | -24.02 | ±0.60 | 0.60 | ±0.05 | 0.88 | <0.001 |
|  | AN20 | 31 | -22.34 | ±1.12 | 0.50 | ±0.07 | 0.63 | <0.001 |
|  | AO20 | 31 | -23.53 | ±1.01 | 0.69 | ±0.07 | 0.76 | <0.001 |
|  | AP20 | 32 | -23.76 | ±1.02 | 0.83 | ±0.09 | 0.74 | <0.001 |
|  | AQ20 | 35 | -25.05 | ±0.80 | 0.88 | ±0.07 | 0.83 | <0.001 |
|  | AR20 | 38 | -24.03 | ±0.82 | 1.04 | ±0.07 | 0.85 | <0.001 |
|  | AS20 | 37 | -24.39 | ±0.65 | 1.16 | ±0.06 | 0.92 | <0.001 |
|  | AT20 | 34 | -23.72 | ±0.76 | 0.91 | ±0.08 | 0.80 | <0.001 |
|  | AU20 | 47 | -24.20 | ±0.73 | 1.27 | ±0.09 | 0.81 | <0.001 |
|  | AV20 | 35 | -22.58 | ±0.65 | 0.85 | ±0.07 | 0.81 | <0.001 |
|  | AW20 | 47 | -23.35 | ±0.62 | 1.11 | ±0.08 | 0.82 | <0.001 |
|  | AX20 | 43 | -23.00 | ±0.85 | 1.36 | ±0.13 | 0.71 | <0.001 |
|  | AY20 | 37 | -22.32 | ±0.47 | 1.31 | ±0.09 | 0.85 | <0.001 |
|  | AZ20 | 42 | -21.90 | ±0.73 | 1.40 | ±0.13 | 0.75 | <0.001 |
|  | AA21 | 12 | -4.23 | ±0.25 | -0.08 | ±0.03 | 0.34 | 0.047 |
|  | AB21 | 16 | -4.21 | ±0.54 | 0.00 | ±0.07 | 0.00 | 0.947 |
|  | AC21 | 36 | -8.08 | ±1.04 | 0.10 | ±0.11 | 0.02 | 0.373 |
|  | AD21 | 46 | -12.09 | ±0.82 | 0.15 | ±0.06 | 0.13 | 0.013 |
|  | AE21 | 43 | -18.68 | ±0.69 | 0.37 | ±0.04 | 0.67 | <0.001 |
|  | AF21 | 47 | -20.60 | ±0.50 | 0.48 | ±0.04 | 0.78 | <0.001 |
|  | AG21 | 52 | -18.87 | ±0.55 | 0.16 | ±0.03 | 0.41 | <0.001 |
|  | AH21 | 54 | -20.63 | ±0.45 | 0.22 | ±0.03 | 0.58 | <0.001 |
|  | AI21 | 63 | -19.95 | ±0.38 | 0.16 | ±0.02 | 0.62 | <0.001 |
|  | AJ21 | 63 | -20.47 | ±0.43 | 0.19 | ±0.02 | 0.60 | <0.001 |
|  | AK21 | 65 | -19.97 | ±0.53 | 0.18 | ±0.02 | 0.55 | <0.001 |
|  | AL21 | 61 | -20.43 | ±0.76 | 0.22 | ±0.03 | 0.41 | <0.001 |
|  | AM21 | 57 | -21.33 | ±0.56 | 0.24 | ±0.03 | 0.50 | <0.001 |
|  | AN21 | 56 | -22.04 | ±0.69 | 0.27 | ±0.04 | 0.43 | <0.001 |
|  | AO21 | 46 | -22.65 | ±0.34 | 0.28 | ±0.03 | 0.69 | <0.001 |
|  | AP21 | 46 | -21.52 | ±0.30 | 0.21 | ±0.02 | 0.67 | <0.001 |
|  | AQ21 | 44 | -21.60 | ±0.86 | 0.52 | ±0.07 | 0.58 | <0.001 |
|  | AR21 | 55 | -23.22 | ±0.84 | 0.65 | ±0.07 | 0.64 | <0.001 |
|  | AS21 | 52 | -22.50 | ±0.60 | 0.66 | ±0.06 | 0.74 | <0.001 |
|  | AT21 | 50 | -21.83 | ±0.78 | 0.69 | ±0.07 | 0.64 | <0.001 |
|  | AU21 | 59 | -22.57 | ±0.49 | 0.77 | ±0.05 | 0.80 | <0.001 |
|  | AV21 | 51 | -22.26 | ±0.56 | 0.81 | ±0.08 | 0.68 | <0.001 |
|  | AW21 | 51 | -22.45 | ±0.52 | 0.70 | ±0.08 | 0.59 | <0.001 |
|  | AX21 | 44 | -21.14 | ±0.26 | 0.63 | ±0.11 | 0.44 | <0.001 |
| *Picea abies* | AA20 | 39 | -15.98 | ±1.07 | 0.01 | ±0.15 | 0.00 | 0.935 |
|  | AB20 | 38 | -14.32 | ±1.06 | -0.14 | ±0.17 | 0.02 | 0.403 |
|  | AC20 | 36 | -15.92 | ±1.24 | -0.23 | ±0.22 | 0.03 | 0.297 |
|  | AD20 | 34 | -15.77 | ±0.97 | -0.24 | ±0.14 | 0.09 | 0.091 |
|  | AE20 | 40 | -17.84 | ±0.65 | -0.22 | ±0.08 | 0.17 | 0.008 |
|  | AF20 | 46 | -18.85 | ±0.53 | -0.22 | ±0.06 | 0.23 | 0.001 |
|  | AG20 | 35 | -21.09 | ±0.61 | -0.02 | ±0.04 | 0.01 | 0.600 |
|  | AH20 | 29 | -20.59 | ±0.53 | -0.05 | ±0.03 | 0.11 | 0.082 |
|  | AI20 | 27 | -23.07 | ±0.73 | 0.05 | ±0.03 | 0.11 | 0.089 |
|  | AJ20 | 25 | -21.19 | ±1.21 | 0.04 | ±0.06 | 0.02 | 0.481 |
|  | AK20 | 29 | -22.49 | ±0.76 | 0.18 | ±0.04 | 0.45 | <0.001 |
|  | AL20 | 21 | -23.56 | ±0.74 | 0.13 | ±0.04 | 0.37 | 0.003 |
|  | AM20 | 21 | -25.54 | ±1.00 | 0.57 | ±0.06 | 0.83 | <0.001 |
|  | AN20 | 27 | -23.09 | ±1.08 | 0.35 | ±0.06 | 0.58 | <0.001 |
|  | AO20 | 29 | -25.37 | ±0.75 | 0.63 | ±0.05 | 0.86 | <0.001 |
|  | AP20 | 28 | -23.47 | ±1.27 | 0.59 | ±0.11 | 0.54 | <0.001 |
|  | AQ20 | 29 | -25.19 | ±0.91 | 0.83 | ±0.08 | 0.80 | <0.001 |
|  | AR20 | 30 | -23.62 | ±0.65 | 0.65 | ±0.06 | 0.81 | <0.001 |
|  | AS20 | 38 | -23.58 | ±1.02 | 0.60 | ±0.09 | 0.55 | <0.001 |
|  | AT20 | 35 | -23.52 | ±0.57 | 0.63 | ±0.06 | 0.75 | <0.001 |
|  | AU20 | 35 | -23.39 | ±0.72 | 0.70 | ±0.10 | 0.58 | <0.001 |
|  | AV20 | 30 | -24.48 | ±0.54 | 0.99 | ±0.10 | 0.78 | <0.001 |
|  | AW20 | 31 | -25.27 | ±0.49 | 0.72 | ±0.08 | 0.75 | <0.001 |
|  | AX20 | 35 | -24.79 | ±0.35 | 1.36 | ±0.07 | 0.93 | <0.001 |
|  | AY20 | 22 | -25.41 | ±0.63 | 2.44 | ±0.25 | 0.83 | <0.001 |
|  | AZ20 | 32 | -23.09 | ±0.66 | 0.86 | ±0.14 | 0.55 | <0.001 |
|  | BA20 | 41 | -23.73 | ±0.45 | 1.40 | ±0.14 | 0.72 | <0.001 |
|  | AA21 | 20 | -18.46 | ±0.56 | -0.06 | ±0.09 | 0.02 | 0.515 |
|  | AB21 | 40 | -17.39 | ±0.70 | -0.37 | ±0.09 | 0.33 | <0.001 |
|  | AC21 | 48 | -17.97 | ±0.60 | -0.22 | ±0.06 | 0.26 | <0.001 |
|  | AD21 | 44 | -19.96 | ±0.58 | -0.16 | ±0.06 | 0.15 | 0.008 |
|  | AE21 | 38 | -19.50 | ±0.65 | -0.06 | ±0.04 | 0.06 | 0.155 |
|  | AF21 | 37 | -21.24 | ±0.62 | 0.04 | ±0.04 | 0.02 | 0.435 |
|  | AG21 | 36 | -22.07 | ±0.43 | 0.06 | ±0.03 | 0.10 | 0.056 |
|  | AH21 | 43 | -21.17 | ±0.50 | 0.03 | ±0.03 | 0.04 | 0.211 |
|  | AI21 | 31 | -20.82 | ±0.54 | -0.09 | ±0.05 | 0.10 | 0.091 |
|  | AJ21 | 44 | -21.25 | ±0.33 | 0.00 | ±0.02 | 0.00 | 0.822 |
|  | AK21 | 47 | -22.91 | ±0.34 | 0.06 | ±0.02 | 0.24 | <0.001 |
|  | AL21 | 56 | -22.22 | ±0.61 | 0.17 | ±0.03 | 0.44 | <0.001 |
|  | AM21 | 40 | -23.70 | ±0.84 | 0.22 | ±0.05 | 0.36 | <0.001 |
|  | AN21 | 50 | -22.40 | ±0.82 | 0.18 | ±0.05 | 0.22 | 0.001 |
|  | AO21 | 44 | -22.61 | ±0.60 | 0.33 | ±0.05 | 0.52 | <0.001 |
|  | AP21 | 47 | -23.80 | ±0.69 | 0.34 | ±0.05 | 0.50 | <0.001 |
|  | AQ21 | 42 | -23.24 | ±0.90 | 0.48 | ±0.07 | 0.54 | <0.001 |
|  | AR21 | 37 | -23.61 | ±0.79 | 0.50 | ±0.07 | 0.60 | <0.001 |
|  | AS21 | 45 | -23.48 | ±0.94 | 0.63 | ±0.08 | 0.57 | <0.001 |
|  | AT21 | 48 | -24.57 | ±0.62 | 0.77 | ±0.08 | 0.67 | <0.001 |
|  | AU21 | 47 | -24.83 | ±0.99 | 0.74 | ±0.14 | 0.38 | <0.001 |
|  | AV21 | 43 | -25.50 | ±1.09 | 1.18 | ±0.16 | 0.57 | <0.001 |
|  | AW21 | 46 | -25.13 | ±0.68 | 1.19 | ±0.14 | 0.63 | <0.001 |
|  | AX21 | 43 | -24.26 | ±0.81 | 1.74 | ±0.32 | 0.43 | <0.001 |
| *Prunus armeniaca* | AA20 | 21 | -5.54 | ±0.62 | 0.01 | ±0.09 | 0.00 | 0.951 |
|  | AB20 | 24 | -4.87 | ±0.62 | 0.08 | ±0.10 | 0.03 | 0.410 |
|  | AC20 | 19 | -4.62 | ±0.75 | -0.01 | ±0.13 | 0.00 | 0.934 |
|  | AD20 | 28 | -4.75 | ±0.88 | -0.30 | ±0.13 | 0.18 | 0.025 |
|  | AE20 | 39 | -6.75 | ±0.89 | -0.27 | ±0.08 | 0.24 | 0.001 |
|  | AF20 | 60 | -10.71 | ±0.69 | -0.18 | ±0.07 | 0.09 | 0.021 |
|  | AG20 | 56 | -15.18 | ±0.56 | 0.11 | ±0.04 | 0.14 | 0.004 |
|  | AH20 | 45 | -16.13 | ±0.65 | 0.13 | ±0.04 | 0.22 | 0.001 |
|  | AI20 | 34 | -17.22 | ±0.68 | 0.18 | ±0.03 | 0.47 | <0.001 |
|  | AJ20 | 35 | -19.09 | ±0.59 | 0.27 | ±0.04 | 0.61 | <0.001 |
|  | AK20 | 25 | -20.71 | ±0.75 | 0.45 | ±0.08 | 0.60 | <0.001 |
|  | AL20 | 16 | -21.12 | ±0.96 | 0.57 | ±0.08 | 0.78 | <0.001 |
|  | AM20 | 20 | -22.00 | ±0.94 | 0.54 | ±0.08 | 0.72 | <0.001 |
|  | AN20 | 27 | -22.17 | ±1.10 | 0.62 | ±0.10 | 0.63 | <0.001 |
|  | AO20 | 27 | -22.07 | ±0.49 | 0.65 | ±0.05 | 0.85 | <0.001 |
|  | AP20 | 39 | -21.71 | ±0.65 | 0.62 | ±0.07 | 0.68 | <0.001 |
|  | AQ20 | 43 | -21.08 | ±0.67 | 0.65 | ±0.06 | 0.73 | <0.001 |
|  | AR20 | 27 | -19.21 | ±0.97 | 1.14 | ±0.19 | 0.59 | <0.001 |
|  | AS20 | 18 | -20.84 | ±0.87 | 2.70 | ±0.31 | 0.83 | <0.001 |
|  | AT20 | 17 | -18.25 | ±0.77 | 3.61 | ±0.40 | 0.84 | <0.001 |
|  | AU20 | 21 | -18.84 | ±1.27 | 4.72 | ±0.91 | 0.59 | <0.001 |
|  | AV20 | 20 | -18.12 | ±1.07 | 5.80 | ±0.75 | 0.77 | <0.001 |
|  | AW20 | 46 | -12.81 | ±1.19 | 2.06 | ±0.49 | 0.29 | <0.001 |
|  | AX20 | 28 | -13.74 | ±1.09 | 4.41 | ±0.71 | 0.59 | <0.001 |
|  | AY20 | 20 | -14.89 | ±1.00 | 7.59 | ±1.25 | 0.67 | <0.001 |
|  | AZ20 | 11 | -15.60 | ±0.36 | 11.00 | ±0.85 | 0.95 | <0.001 |
|  | AA21 | 10 | -4.74 | ±0.54 | 0.11 | ±0.08 | 0.21 | 0.183 |
|  | AB21 | 24 | -2.29 | ±1.18 | -0.67 | ±0.14 | 0.49 | <0.001 |
|  | AC21 | 37 | -14.47 | ±1.14 | 0.28 | ±0.10 | 0.20 | 0.006 |
|  | AD21 | 43 | -10.67 | ±1.15 | 0.11 | ±0.08 | 0.04 | 0.175 |
|  | AE21 | 39 | -13.73 | ±0.89 | 0.15 | ±0.05 | 0.20 | 0.005 |
|  | AF21 | 38 | -13.98 | ±0.77 | 0.18 | ±0.05 | 0.24 | 0.002 |
|  | AG21 | 55 | -15.63 | ±0.52 | 0.12 | ±0.04 | 0.16 | 0.002 |
|  | AH21 | 50 | -18.24 | ±0.67 | 0.27 | ±0.06 | 0.28 | <0.001 |
|  | AI21 | 56 | -17.90 | ±0.74 | 0.21 | ±0.07 | 0.15 | 0.003 |
|  | AJ21 | 69 | -16.88 | ±0.50 | 0.31 | ±0.05 | 0.42 | <0.001 |
|  | AK21 | 65 | -20.61 | ±0.54 | 0.55 | ±0.05 | 0.69 | <0.001 |
|  | AL21 | 82 | -18.78 | ±0.43 | 0.23 | ±0.03 | 0.35 | <0.001 |
|  | AM21 | 55 | -21.00 | ±0.47 | 0.53 | ±0.04 | 0.73 | <0.001 |
|  | AN21 | 56 | -21.83 | ±0.65 | 0.59 | ±0.06 | 0.68 | <0.001 |
|  | AO21 | 53 | -21.78 | ±0.73 | 0.73 | ±0.06 | 0.74 | <0.001 |
|  | AP21 | 89 | -21.14 | ±0.51 | 0.67 | ±0.05 | 0.65 | <0.001 |
|  | AQ21 | 89 | -20.92 | ±0.44 | 0.72 | ±0.06 | 0.62 | <0.001 |
|  | AR21 | 85 | -20.35 | ±0.54 | 0.91 | ±0.08 | 0.62 | <0.001 |
|  | AS21 | 65 | -18.85 | ±0.83 | 1.21 | ±0.15 | 0.50 | <0.001 |
|  | AT21 | 63 | -18.52 | ±0.80 | 1.52 | ±0.18 | 0.54 | <0.001 |
|  | AU21 | 56 | -18.31 | ±1.05 | 2.49 | ±0.35 | 0.49 | <0.001 |
|  | AV21 | 42 | -18.69 | ±1.16 | 3.52 | ±0.47 | 0.58 | <0.001 |
|  | AW21 | 48 | -18.58 | ±1.07 | 3.65 | ±0.44 | 0.60 | <0.001 |
|  | AX21 | 27 | -18.16 | ±0.86 | 7.65 | ±0.69 | 0.83 | <0.001 |
| *Prunus nigra* | AA20 | 26 | -5.23 | ±1.00 | -0.08 | ±0.13 | 0.01 | 0.560 |
|  | AB20 | 22 | -7.69 | ±0.92 | 0.49 | ±0.16 | 0.32 | 0.006 |
|  | AC20 | 18 | -5.34 | ±0.44 | 0.18 | ±0.09 | 0.20 | 0.063 |
|  | AD20 | 14 | -4.49 | ±0.68 | 0.02 | ±0.11 | 0.00 | 0.848 |
|  | AE20 | 15 | -4.26 | ±0.30 | 0.05 | ±0.04 | 0.09 | 0.266 |
|  | AF20 | 21 | -4.14 | ±0.22 | 0.07 | ±0.03 | 0.18 | 0.057 |
|  | AG20 | 11 | -3.19 | ±0.20 | -0.03 | ±0.02 | 0.23 | 0.135 |
|  | AH20 | 14 | -6.64 | 2.17 | -0.32 | ±0.10 | 0.46 | 0.008 |
|  | AI20 | 24 | -15.69 | ±1.29 | -0.08 | ±0.05 | 0.12 | 0.100 |
|  | AJ20 | 19 | -20.83 | ±1.10 | 0.13 | ±0.06 | 0.26 | 0.025 |
|  | AK20 | 22 | -25.19 | ±1.38 | 0.27 | ±0.06 | 0.47 | <0.001 |
|  | AL20 | 15 | -13.94 | ±1.15 | 0.27 | ±0.07 | 0.51 | 0.003 |
|  | AM20 | 13 | -23.01 | ±1.25 | 0.33 | ±0.09 | 0.55 | 0.004 |
|  | AN20 | 31 | -26.96 | ±1.00 | 0.44 | ±0.06 | 0.64 | <0.001 |
|  | AO20 | 29 | -28.51 | ±0.89 | 0.45 | ±0.05 | 0.72 | <0.001 |
|  | AP20 | 29 | -26.00 | ±0.87 | 0.39 | ±0.08 | 0.48 | <0.001 |
|  | AQ20 | 27 | -29.47 | ±0.72 | 0.78 | ±0.08 | 0.79 | <0.001 |
|  | AR20 | 26 | -26.35 | ±1.24 | 0.73 | ±0.10 | 0.71 | <0.001 |
|  | AS20 | 33 | -26.93 | ±1.07 | 1.11 | ±0.08 | 0.85 | <0.001 |
|  | AT20 | 24 | -27.43 | ±1.23 | 0.85 | ±0.12 | 0.70 | <0.001 |
|  | AU20 | 23 | -24.67 | ±1.01 | 0.81 | ±0.19 | 0.46 | <0.001 |
|  | AV20 | 21 | -23.81 | ±0.80 | 1.24 | ±0.31 | 0.46 | 0.001 |
|  | AW20 | 31 | -22.55 | ±1.06 | 1.23 | ±0.21 | 0.54 | <0.001 |
|  | AX20 | 26 | -21.49 | ±0.69 | 1.92 | ±0.26 | 0.69 | <0.001 |
|  | AY20 | 23 | -20.41 | ±0.52 | 1.25 | ±0.21 | 0.62 | <0.001 |
|  | AZ20 | 22 | -14.98 | ±1.46 | 1.55 | ±0.46 | 0.36 | 0.003 |
|  | AA21 | 9 | -3.90 | ±0.50 | 0.00 | ±0.07 | 0.00 | 0.958 |
|  | AB21 | 15 | -3.61 | ±1.65 | -0.26 | ±0.22 | 0.09 | 0.275 |
|  | AC21 | 18 | -5.19 | ±0.64 | 0.10 | ±0.07 | 0.11 | 0.173 |
|  | AD21 | 19 | -3.59 | ±0.92 | -0.18 | ±0.07 | 0.30 | 0.016 |
|  | AE21 | 19 | -6.01 | ±1.13 | -0.54 | ±0.06 | 0.82 | <0.001 |
|  | AF21 | 14 | -2.12 | 2.91 | -0.89 | ±0.19 | 0.65 | 0.001 |
|  | AG21 | 21 | -9.61 | ±1.89 | -0.43 | ±0.12 | 0.41 | 0.002 |
|  | AH21 | 20 | -18.83 | ±1.18 | -0.06 | ±0.10 | 0.02 | 0.580 |
|  | AI21 | 31 | -19.46 | ±1.04 | -0.05 | ±0.08 | 0.01 | 0.525 |
|  | AJ21 | 34 | -18.73 | ±1.06 | -0.12 | ±0.09 | 0.06 | 0.168 |
|  | AK21 | 57 | -21.24 | ±0.65 | 0.07 | ±0.03 | 0.09 | 0.021 |
|  | AL21 | 48 | -24.37 | ±0.73 | 0.24 | ±0.03 | 0.56 | <0.001 |
|  | AM21 | 41 | -22.60 | ±0.85 | 0.22 | ±0.05 | 0.35 | <0.001 |
|  | AN21 | 46 | -22.37 | ±0.79 | 0.19 | ±0.05 | 0.27 | <0.001 |
|  | AO21 | 50 | -24.93 | ±0.84 | 0.40 | ±0.06 | 0.46 | <0.001 |
|  | AP21 | 49 | -23.94 | ±0.89 | 0.34 | ±0.07 | 0.32 | <0.001 |
|  | AQ21 | 45 | -24.90 | ±0.94 | 0.56 | ±0.07 | 0.57 | <0.001 |
|  | AR21 | 51 | -26.85 | ±0.74 | 0.55 | ±0.06 | 0.66 | <0.001 |
|  | AS21 | 51 | -24.34 | ±0.96 | 0.53 | ±0.12 | 0.28 | <0.001 |
|  | AT21 | 56 | -26.48 | ±0.85 | 0.75 | ±0.12 | 0.41 | <0.001 |
|  | AU21 | 44 | -28.91 | ±1.04 | 1.35 | ±0.17 | 0.60 | <0.001 |
|  | AV21 | 51 | -24.66 | ±0.78 | 0.79 | ±0.13 | 0.44 | <0.001 |
|  | AW21 | 69 | -25.62 | ±0.67 | 1.84 | ±0.18 | 0.60 | <0.001 |
| *Rhododendron calendulaceum* | AB20 | 16 | -9.27 | ±0.95 | 0.12 | ±0.14 | 0.05 | 0.428 |
|  | AC20 | 23 | -8.43 | ±0.73 | 0.02 | ±0.14 | 0.00 | 0.863 |
|  | AD20 | 48 | -14.43 | ±0.83 | 0.08 | ±0.12 | 0.01 | 0.525 |
|  | AE20 | 59 | -15.01 | ±0.62 | 0.01 | ±0.06 | 0.00 | 0.857 |
|  | AF20 | 75 | -17.47 | ±0.55 | 0.01 | ±0.06 | 0.00 | 0.933 |
|  | AG20 | 59 | -17.76 | ±0.63 | -0.02 | ±0.04 | 0.01 | 0.576 |
|  | AH20 | 39 | -20.14 | ±0.74 | 0.10 | ±0.06 | 0.07 | 0.106 |
|  | AI20 | 30 | -23.58 | ±0.43 | 0.12 | ±0.03 | 0.42 | <0.001 |
|  | AJ20 | 40 | -24.81 | ±0.42 | 0.11 | ±0.03 | 0.25 | 0.001 |
|  | AK20 | 40 | -22.48 | ±0.55 | 0.10 | ±0.03 | 0.19 | 0.005 |
|  | AL20 | 38 | -25.76 | ±0.55 | 0.27 | ±0.03 | 0.68 | <0.001 |
|  | AM20 | 30 | -26.86 | ±0.50 | 0.31 | ±0.05 | 0.59 | <0.001 |
|  | AN20 | 48 | -25.53 | ±0.60 | 0.20 | ±0.03 | 0.43 | <0.001 |
|  | AO20 | 49 | -28.53 | ±0.59 | 0.41 | ±0.04 | 0.71 | <0.001 |
|  | AP20 | 39 | -26.42 | ±0.52 | 0.35 | ±0.06 | 0.52 | <0.001 |
|  | AQ20 | 48 | -25.60 | ±0.46 | 0.25 | ±0.04 | 0.44 | <0.001 |
|  | AR20 | 39 | -24.63 | ±0.61 | 0.22 | ±0.07 | 0.19 | 0.005 |
|  | AS20 | 37 | -26.19 | ±0.67 | 0.34 | ±0.08 | 0.35 | <0.001 |
|  | AT20 | 49 | -25.41 | ±0.53 | 0.22 | ±0.06 | 0.23 | <0.001 |
|  | AU20 | 50 | -26.41 | ±0.62 | 0.40 | ±0.09 | 0.31 | <0.001 |
|  | AV20 | 40 | -26.06 | ±0.45 | 0.36 | ±0.08 | 0.34 | <0.001 |
|  | AW20 | 30 | -26.91 | ±0.58 | 0.54 | ±0.07 | 0.66 | <0.001 |
|  | AX20 | 50 | -27.98 | ±0.39 | 0.51 | ±0.06 | 0.63 | <0.001 |
|  | AY20 | 40 | -27.18 | ±0.30 | 0.43 | ±0.06 | 0.62 | <0.001 |
|  | AZ20 | 40 | -25.69 | ±0.42 | 0.52 | ±0.09 | 0.48 | <0.001 |
|  | AA21 | 35 | -10.32 | ±0.58 | -0.12 | ±0.09 | 0.05 | 0.189 |
|  | AB21 | 50 | -10.66 | ±0.90 | -0.07 | ±0.11 | 0.01 | 0.527 |
|  | AC21 | 50 | -14.26 | ±0.98 | -0.04 | ±0.12 | 0.00 | 0.744 |
|  | AD21 | 60 | -18.08 | ±0.60 | 0.13 | ±0.04 | 0.14 | 0.003 |
|  | AE21 | 50 | -16.92 | ±0.65 | 0.08 | ±0.04 | 0.08 | 0.047 |
|  | AF21 | 50 | -18.60 | ±0.81 | 0.14 | ±0.06 | 0.11 | 0.020 |
|  | AG21 | 50 | -19.10 | ±0.55 | 0.03 | ±0.04 | 0.01 | 0.476 |
|  | AH21 | 50 | -22.62 | ±0.61 | 0.22 | ±0.06 | 0.24 | <0.001 |
|  | AI21 | 50 | -20.73 | ±0.63 | 0.01 | ±0.06 | 0.00 | 0.830 |
|  | AJ21 | 60 | -21.31 | ±0.51 | 0.04 | ±0.03 | 0.03 | 0.201 |
|  | AK21 | 59 | -21.79 | ±0.44 | 0.07 | ±0.02 | 0.13 | 0.004 |
|  | AL21 | 60 | -21.44 | ±0.47 | 0.06 | ±0.03 | 0.08 | 0.033 |
|  | AM21 | 59 | -23.39 | ±0.49 | 0.18 | ±0.03 | 0.39 | <0.001 |
|  | AN21 | 60 | -23.22 | ±0.50 | 0.13 | ±0.03 | 0.23 | <0.001 |
|  | AO21 | 49 | -24.82 | ±0.72 | 0.22 | ±0.06 | 0.22 | 0.001 |
|  | AP21 | 49 | -25.11 | ±0.57 | 0.25 | ±0.04 | 0.41 | <0.001 |
|  | AQ21 | 47 | -24.21 | ±0.62 | 0.27 | ±0.05 | 0.40 | <0.001 |
|  | AR21 | 50 | -25.54 | ±0.72 | 0.33 | ±0.06 | 0.38 | <0.001 |
|  | AS21 | 49 | -25.23 | ±0.58 | 0.30 | ±0.07 | 0.30 | <0.001 |
|  | AT21 | 50 | -26.61 | ±0.63 | 0.40 | ±0.08 | 0.33 | <0.001 |
|  | AU21 | 50 | -26.85 | ±0.65 | 0.55 | ±0.11 | 0.36 | <0.001 |
|  | AV21 | 50 | -27.00 | ±0.71 | 0.53 | ±0.14 | 0.24 | <0.001 |
|  | AW21 | 50 | -28.13 | ±0.61 | 0.60 | ±0.12 | 0.33 | <0.001 |
|  | AX21 | 50 | -27.12 | ±0.76 | 0.36 | ±0.31 | 0.03 | 0.248 |
