## Supplementary material for "Woody species do not differ in dormancy progression: differences in time to budbreak due to forcing and cold hardiness": Table S5

**Supporting Information Table S4. Estimates, fitness statistics, and displays associated with linear regressions for each temperature and species.**

| Species | Chill | T (ºC) | n | α ± error | | β ± error | | *r^2^* | *P*-value |
| --- | --- | --- | --- | --- | --- | --- | --- | --- | --- |
| *Abies balsamea* | 54 | 2 | 26 | -28.22 | ±0.34 | 0.09 | ±0.01 | 0.66 | <0.001 |
|  |  | 4 | 34 | -25.34 | ±1.44 | 0.42 | ±0.06 | 0.63 | <0.001 |
|  |  | 7 | 34 | -25.50 | ±2.32 | 0.27 | ±0.12 | 0.14 | 0.027 |
|  |  | 11 | 33 | -23.75 | ±2.72 | 0.51 | ±0.19 | 0.19 | 0.010 |
|  |  | 15 | 38 | -24.39 | ±2.32 | 0.71 | ±0.19 | 0.29 | 0.001 |
|  |  | 22 | 45 | -28.00 | ±0.51 | 0.54 | ±0.04 | 0.78 | <0.001 |
|  |  | 30 | 31 | -25.81 | ±0.69 | 1.15 | ±0.15 | 0.66 | <0.001 |
|  | 82 | 2 | 39 | -27.93 | ±0.41 | 0.04 | ±0.02 | 0.09 | 0.070 |
|  |  | 4 | 38 | -26.99 | ±0.25 | 0.02 | ±0.01 | 0.04 | 0.252 |
|  |  | 7 | 49 | -27.08 | ±0.30 | 0.01 | ±0.02 | 0.00 | 0.765 |
|  |  | 11 | 48 | -27.52 | ±0.60 | 0.19 | ±0.03 | 0.43 | <0.001 |
|  |  | 15 | 46 | -27.65 | ±0.69 | 0.68 | ±0.08 | 0.64 | <0.001 |
|  |  | 22 | 52 | -26.10 | ±0.63 | 0.75 | ±0.08 | 0.64 | <0.001 |
|  |  | 30 | 65 | -28.12 | ±0.53 | 1.49 | ±0.13 | 0.67 | <0.001 |
| *Acer rubrum* | 54 | 2 | 30 | -25.86 | ±0.67 | -0.02 | ±0.03 | 0.01 | 0.539 |
|  |  | 4 | 40 | -26.89 | ±0.70 | 0.04 | ±0.03 | 0.07 | 0.090 |
|  |  | 7 | 50 | -25.90 | ±0.87 | 0.04 | ±0.03 | 0.03 | 0.226 |
|  |  | 11 | 60 | -25.36 | ±0.85 | 0.10 | ±0.04 | 0.11 | 0.009 |
|  |  | 15 | 50 | -26.44 | ±0.95 | 0.27 | ±0.04 | 0.48 | <0.001 |
|  |  | 22 | 70 | -25.56 | ±0.71 | 0.28 | ±0.06 | 0.23 | <0.001 |
|  |  | 30 | 70 | -25.27 | ±0.56 | 0.56 | ±0.07 | 0.50 | <0.001 |
|  | 82 | 2 | 70 | -25.36 | ±0.70 | 0.19 | ±0.03 | 0.34 | <0.001 |
|  |  | 4 | 78 | -24.50 | ±0.46 | 0.19 | ±0.02 | 0.50 | <0.001 |
|  |  | 7 | 100 | -24.58 | ±0.59 | 0.40 | ±0.03 | 0.59 | <0.001 |
|  |  | 11 | 88 | -24.67 | ±0.44 | 0.72 | ±0.05 | 0.72 | <0.001 |
|  |  | 15 | 88 | -24.00 | ±0.59 | 1.09 | ±0.09 | 0.63 | <0.001 |
|  |  | 22 | 90 | -23.54 | ±0.61 | 1.48 | ±0.11 | 0.67 | <0.001 |
|  |  | 30 | 58 | -23.05 | ±0.95 | 2.95 | ±0.31 | 0.62 | <0.001 |
| *Cercis canadensis* | 54 | 2 | 30 | -16.41 | ±0.61 | -0.02 | ±0.02 | 0.02 | 0.430 |
|  |  | 4 | 40 | -15.74 | ±0.42 | -0.02 | ±0.02 | 0.05 | 0.150 |
|  |  | 7 | 47 | -18.25 | ±0.75 | 0.15 | ±0.03 | 0.43 | <0.001 |
|  |  | 11 | 59 | -17.11 | ±0.84 | 0.14 | ±0.04 | 0.21 | <0.001 |
|  |  | 15 | 40 | -16.45 | ±0.53 | 0.24 | ±0.03 | 0.58 | <0.001 |
|  |  | 22 | 79 | -17.59 | ±0.36 | 0.49 | ±0.04 | 0.71 | <0.001 |
|  |  | 30 | 89 | -15.17 | ±0.26 | 0.50 | ±0.04 | 0.70 | <0.001 |
|  | 82 | 2 | 66 | -15.43 | ±0.46 | 0.13 | ±0.02 | 0.40 | <0.001 |
|  |  | 4 | 63 | -13.99 | ±0.32 | 0.14 | ±0.01 | 0.60 | <0.001 |
|  |  | 7 | 72 | -14.72 | ±0.33 | 0.26 | ±0.02 | 0.71 | <0.001 |
|  |  | 11 | 60 | -14.29 | ±0.37 | 0.43 | ±0.04 | 0.67 | <0.001 |
|  |  | 15 | 80 | -14.74 | ±0.34 | 0.59 | ±0.05 | 0.67 | <0.001 |
|  |  | 22 | 66 | -14.30 | ±0.42 | 0.88 | ±0.07 | 0.69 | <0.001 |
|  |  | 30 | 55 | -14.32 | ±0.57 | 1.25 | ±0.15 | 0.55 | <0.001 |
| *Cornus florida* | 54 | 2 | 30 | -23.69 | ±0.49 | 0.01 | ±0.02 | 0.02 | 0.457 |
|  |  | 4 | 30 | -23.68 | ±0.49 | 0.03 | ±0.02 | 0.07 | 0.148 |
|  |  | 7 | 50 | -24.34 | ±0.41 | -0.01 | ±0.01 | 0.00 | 0.636 |
|  |  | 11 | 60 | -23.06 | ±0.69 | -0.01 | ±0.03 | 0.00 | 0.596 |
|  |  | 15 | 50 | -23.80 | ±0.61 | 0.10 | ±0.03 | 0.23 | <0.001 |
|  |  | 22 | 70 | -23.87 | ±0.63 | 0.37 | ±0.05 | 0.41 | <0.001 |
|  |  | 30 | 60 | -23.33 | ±0.61 | 0.42 | ±0.07 | 0.38 | <0.001 |
| *Cornus mas* | 54 | 2 | 40 | -20.88 | ±0.84 | 0.05 | ±0.04 | 0.05 | 0.172 |
|  |  | 4 | 40 | -20.90 | ±0.62 | 0.03 | ±0.03 | 0.03 | 0.310 |
|  |  | 7 | 70 | -21.10 | ±0.61 | 0.06 | ±0.02 | 0.08 | 0.019 |
|  |  | 11 | 70 | -21.19 | ±0.70 | 0.23 | ±0.04 | 0.33 | <0.001 |
|  |  | 15 | 90 | -20.75 | ±0.77 | 0.23 | ±0.08 | 0.09 | 0.004 |
|  |  | 22 | 80 | -20.30 | ±0.72 | 0.91 | ±0.16 | 0.30 | <0.001 |
|  |  | 30 | 80 | -20.09 | ±0.68 | 1.14 | ±0.13 | 0.50 | <0.001 |
|  | 82 | 2 | 50 | -21.27 | ±0.52 | 0.12 | ±0.03 | 0.30 | <0.001 |
|  |  | 4 | 90 | -21.37 | ±0.64 | 0.12 | ±0.03 | 0.17 | <0.001 |
|  |  | 7 | 110 | -21.19 | ±0.47 | 0.22 | ±0.03 | 0.33 | <0.001 |
|  |  | 11 | 80 | -19.73 | ±0.63 | 0.30 | ±0.10 | 0.10 | 0.004 |
|  |  | 15 | 90 | -18.40 | ±0.91 | 0.69 | ±0.16 | 0.17 | <0.001 |
|  |  | 22 | 48 | -19.70 | ±0.91 | 1.81 | ±0.26 | 0.52 | <0.001 |
|  |  | 30 | 39 | -18.94 | ±1.11 | 2.98 | ±0.60 | 0.40 | <0.001 |
| *Fagus grandifolia* | 54 | 2 | 25 | -18.53 | ±0.43 | 0.06 | ±0.02 | 0.42 | <0.001 |
|  |  | 4 | 28 | -18.59 | ±0.32 | 0.07 | ±0.01 | 0.55 | <0.001 |
|  |  | 7 | 30 | -18.62 | ±0.32 | 0.12 | ±0.01 | 0.71 | <0.001 |
|  |  | 11 | 35 | -17.57 | ±0.79 | 0.14 | ±0.04 | 0.26 | 0.002 |
|  |  | 15 | 44 | -17.54 | ±0.36 | 0.06 | ±0.02 | 0.28 | <0.001 |
|  |  | 22 | 43 | -18.32 | ±0.31 | 0.26 | ±0.02 | 0.74 | <0.001 |
|  |  | 30 | 46 | -17.46 | ±0.37 | 0.45 | ±0.04 | 0.74 | <0.001 |
| *Forsythia* ‘Meadowlark’ | 54 | 2 | 34 | -22.94 | ±0.68 | 0.02 | ±0.03 | 0.02 | 0.435 |
|  |  | 4 | 43 | -23.11 | ±0.52 | 0.07 | ±0.02 | 0.21 | 0.002 |
|  |  | 7 | 55 | -24.52 | ±1.20 | 0.29 | ±0.06 | 0.32 | <0.001 |
|  |  | 11 | 56 | -23.15 | ±1.70 | 0.66 | ±0.13 | 0.33 | <0.001 |
|  |  | 15 | 60 | -23.57 | ±1.19 | 0.90 | ±0.14 | 0.42 | <0.001 |
|  |  | 22 | 65 | -23.82 | ±1.18 | 1.43 | ±0.25 | 0.34 | <0.001 |
|  |  | 30 | 81 | -22.59 | ±1.19 | 1.33 | ±0.21 | 0.34 | <0.001 |
|  | 82 | 2 | 92 | -21.42 | ±1.18 | 0.20 | ±0.05 | 0.15 | <0.001 |
|  |  | 4 | 121 | -20.46 | ±0.76 | 0.21 | ±0.04 | 0.20 | <0.001 |
|  |  | 7 | 75 | -22.80 | ±1.24 | 1.26 | ±0.18 | 0.41 | <0.001 |
|  |  | 11 | 56 | -23.57 | ±1.13 | 2.20 | ±0.27 | 0.56 | <0.001 |
|  |  | 15 | 31 | -23.61 | ±1.28 | 4.06 | ±0.72 | 0.52 | <0.001 |
|  |  | 22 | 27 | -20.69 | ±2.00 | 6.58 | ±1.48 | 0.44 | <0.001 |
|  |  | 30 | 28 | -20.10 | ±1.76 | 6.35 | ±1.09 | 0.57 | <0.001 |
| *Kalmia latifolia* | 54 | 2 | 29 | -22.71 | ±1.07 | -0.03 | ±0.04 | 0.02 | 0.506 |
|  |  | 4 | 30 | -22.00 | ±0.86 | -0.06 | ±0.03 | 0.10 | 0.089 |
|  |  | 7 | 49 | -23.49 | ±0.84 | -0.02 | ±0.03 | 0.02 | 0.394 |
|  |  | 11 | 35 | -23.10 | ±1.00 | 0.02 | ±0.05 | 0.01 | 0.635 |
|  |  | 15 | 50 | -23.83 | ±0.90 | 0.17 | ±0.04 | 0.30 | <0.001 |
|  |  | 22 | 59 | -22.47 | ±0.73 | 0.14 | ±0.04 | 0.15 | 0.003 |
|  |  | 30 | 51 | -22.62 | ±0.78 | 0.31 | ±0.09 | 0.18 | 0.002 |
| *Larix kaempferi* | 54 | 2 | 25 | -27.94 | ±0.59 | 0.10 | ±0.02 | 0.46 | <0.001 |
|  |  | 4 | 29 | -27.46 | ±0.48 | 0.09 | ±0.02 | 0.46 | <0.001 |
|  |  | 7 | 44 | -26.65 | ±0.47 | 0.02 | ±0.02 | 0.05 | 0.134 |
|  |  | 11 | 33 | -26.70 | ±0.36 | 0.06 | ±0.02 | 0.28 | 0.002 |
|  |  | 15 | 48 | -27.62 | ±0.47 | 0.18 | ±0.02 | 0.64 | <0.001 |
|  |  | 22 | 56 | -26.20 | ±0.64 | 0.33 | ±0.05 | 0.42 | <0.001 |
|  |  | 30 | 49 | -26.76 | ±0.51 | 0.85 | ±0.07 | 0.78 | <0.001 |
|  | 82 | 2 | 50 | -25.32 | ±0.63 | 0.12 | ±0.03 | 0.29 | <0.001 |
|  |  | 4 | 65 | -25.33 | ±0.45 | 0.16 | ±0.02 | 0.48 | <0.001 |
|  |  | 7 | 58 | -25.34 | ±0.48 | 0.26 | ±0.03 | 0.63 | <0.001 |
|  |  | 11 | 67 | -24.19 | ±0.44 | 0.37 | ±0.03 | 0.64 | <0.001 |
|  |  | 15 | 46 | -23.40 | ±0.62 | 0.50 | ±0.12 | 0.30 | <0.001 |
|  |  | 22 | 59 | -23.76 | ±0.56 | 0.64 | ±0.11 | 0.36 | <0.001 |
|  |  | 30 | 48 | -23.99 | ±0.56 | 0.97 | ±0.14 | 0.50 | <0.001 |
| *Picea abies* | 54 | 2 | 24 | -25.16 | ±0.54 | 0.08 | ±0.02 | 0.46 | <0.001 |
|  |  | 4 | 20 | -25.39 | ±1.33 | 0.21 | ±0.05 | 0.47 | 0.001 |
|  |  | 7 | 30 | -24.09 | ±0.97 | 0.07 | ±0.04 | 0.08 | 0.122 |
|  |  | 11 | 42 | -23.29 | ±1.19 | 0.14 | ±0.06 | 0.14 | 0.014 |
|  |  | 15 | 41 | -25.41 | ±0.79 | 0.43 | ±0.05 | 0.68 | <0.001 |
|  |  | 22 | 39 | -23.60 | ±1.02 | 0.50 | ±0.09 | 0.48 | <0.001 |
|  |  | 30 | 37 | -23.05 | ±0.66 | 1.24 | ±0.11 | 0.80 | <0.001 |
|  | 82 | 2 | 59 | -22.89 | ±0.69 | -0.01 | ±0.03 | 0.00 | 0.635 |
|  |  | 4 | 56 | -21.52 | ±1.37 | 0.09 | ±0.06 | 0.04 | 0.139 |
|  |  | 7 | 68 | -22.91 | ±0.81 | 0.10 | ±0.04 | 0.09 | 0.013 |
|  |  | 11 | 72 | -23.12 | ±0.73 | 0.23 | ±0.04 | 0.34 | <0.001 |
|  |  | 15 | 68 | -23.30 | ±0.60 | 0.38 | ±0.04 | 0.57 | <0.001 |
|  |  | 22 | 42 | -23.98 | ±0.58 | 0.77 | ±0.08 | 0.68 | <0.001 |
|  |  | 30 | 40 | -25.62 | ±0.86 | 2.13 | ±0.29 | 0.59 | <0.001 |
| *Prunus armeniaca* | 54 | 2 | 37 | -21.31 | ±0.41 | 0.07 | ±0.02 | 0.24 | 0.002 |
|  |  | 4 | 50 | -20.99 | ±0.39 | 0.13 | ±0.02 | 0.60 | <0.001 |
|  |  | 7 | 60 | -20.15 | ±0.95 | 0.23 | ±0.05 | 0.30 | <0.001 |
|  |  | 11 | 68 | -20.40 | ±0.72 | 1.09 | ±0.10 | 0.62 | <0.001 |
|  |  | 15 | 40 | -18.18 | ±1.26 | 0.55 | ±0.09 | 0.50 | <0.001 |
|  |  | 22 | 87 | -20.23 | ±0.59 | 0.93 | ±0.09 | 0.58 | <0.001 |
|  |  | 30 | 73 | -17.76 | ±0.95 | 1.11 | ±0.17 | 0.39 | <0.001 |
|  | 82 | 2 | 61 | -17.73 | ±0.40 | 0.10 | ±0.02 | 0.25 | <0.001 |
|  |  | 4 | 96 | -14.32 | ±0.86 | 0.10 | ±0.06 | 0.02 | 0.132 |
|  |  | 7 | 68 | -16.09 | ±1.12 | 1.19 | ±0.20 | 0.35 | <0.001 |
|  |  | 11 | 39 | -16.30 | ±1.05 | 1.90 | ±0.32 | 0.49 | <0.001 |
|  |  | 15 | 29 | -15.89 | ±1.17 | 5.49 | ±0.90 | 0.58 | <0.001 |
|  |  | 22 | 25 | -14.82 | ±1.20 | 5.19 | ±0.90 | 0.59 | <0.001 |
|  |  | 30 | 24 | -18.03 | ±0.79 | 7.40 | ±0.70 | 0.83 | <0.001 |
| *Prunus nigra* | 54 | 2 | 23 | -29.57 | ±1.09 | 0.13 | ±0.04 | 0.34 | 0.003 |
|  |  | 4 | 23 | -27.74 | ±1.35 | 0.10 | ±0.01 | 0.16 | 0.061 |
|  |  | 7 | 43 | -27.75 | ±1.11 | 0.22 | ±0.06 | 0.47 | <0.001 |
|  |  | 11 | 60 | -27.93 | ±0.82 | 0.23 | ±0.12 | 0.48 | <0.001 |
|  |  | 15 | 49 | -22.82 | ±1.60 | 0.16 | ±0.19 | 0.10 | 0.025 |
|  |  | 22 | 74 | -22.69 | ±1.22 | 0.36 | ±0.19 | 0.14 | 0.001 |
|  |  | 30 | 48 | -25.90 | ±1.31 | 1.29 | ±0.04 | 0.58 | <0.001 |
| *Rhododendron calendulaceum* | 54 | 11 | 40 | -24.26 | ±1.36 | 0.29 | ±0.15 | 0.34 | <0.001 |
|  |  | 15 | 50 | -24.88 | ±0.89 | 0.17 | ±0.02 | 0.30 | <0.001 |
|  |  | 22 | 50 | -25.54 | ±0.72 | 0.33 | ±0.01 | 0.38 | <0.001 |
|  |  | 30 | 69 | -24.59 | ±0.66 | 0.82 | ±0.02 | 0.59 | <0.001 |
